# Sequence Dependent UV Damage of Complete Pools of Oligonucleotides

**DOI:** 10.1101/2022.08.01.502267

**Authors:** Corinna L. Kufner, Stefan Krebs, Marlis Fischaleck, Julia Philippou-Massier, Helmut Blum, Dominik B. Bucher, Dieter Braun, Wolfgang Zinth, Christof B. Mast

## Abstract

Understanding the sequence-dependent DNA damage formation requires to probe a complete pool of sequences over a wide dose range of the damage causing exposure. We used high throughput sequencing to simultaneously obtain the dose dependence and quantum yields for oligonucleotide damages for all possible 4096 DNA sequences with hexamer length. We exposed the DNA with ultraviolet radiation at 266 nm and doses of up to 500 photons per base. At the dimer level our results confirm existing literature values, whereas we now quantified the susceptibility of sequence motifs to UV irradiation up to previously inaccessible polymer lengths. This revealed the protective effect of the sequence context in preventing the formation of UV-lesions. For example, the rate to form dipyrimidine lesions is strongly reduced by nearby guanine bases. Our results provide a complete picture of the sensitivity of oligonucleotides to UV irradiation and allow to predict their survival chances in high-UV environments.

---

The genome plays a central role as a blueprint for all known life, therefore damage of the information carrier DNA and the related faulty readout by polymerases is a major threat. In severe cases DNA modifications can result in cell death or in defective organisms. DNA damage may be caused by reactive compounds, for example by reactive oxygen species, or by energetic radiation. In the latter case, energetic particles can break the backbone of DNA strands. Softer radiation in the ultraviolet (UV) range can alter the structure of DNA bases, for example by sequence-specific reactions such as the formation of cyclobutane pyrimidine dimers (CPD). UV-radiation damage of DNA should not only be considered harmful as a threat for the genetic integrity but is currently used in medical application such as cancer treatment or sterilization.

For better protection against DNA damage and for optimizing medical efficacy, a detailed understanding of damage formation and its mode of action is necessary. Information on a molecular level has been obtained especially for short DNA strands by chemical and physical methods. The damage quantum yields and molecular structures of damages in short oligomers and single dimers have been investigated for several decades^1–4^. Important information on the molecular mechanisms of damage formation was obtained. For instance, it has been shown that the CPD lesion is formed predominantly via the singlet channel within sub-picoseconds and molecular reasons for the frequency of damage formation have been discovered^5–11^. The combination of experimental results supplied important knowledge on the damage of short DNA strands^12^.

For long DNA strands e.g. genomic DNA, biological methods such as next-generation/high-throughput sequencing (NGS, HTS), combined with tailored enzyme assays, have provided significant insights^13^. Examples are Excision-seq^14^, CPD-seq^15^, HS-damage-seq^16^, (t)XR-seq^17^ or Adduct-seq^18^. Most of them involve a cascade of enzymatic preparation steps such as fragmentation, repair, ligation, cleavage or digestion. Lesions caused by UV radiation or reactive chemicals such as cisplatin, were successfully excised from the genomic DNA in a region around the damage and identified using specific endonucleases or additionally used lesion-specific antibodies. Enzymatic removal of the damage then allowed for next generation sequencing. The genome-wide characterization of damage formation thus made it possible to pursue previously unknown possibilities for building a platform for DNA damage-induced mutagenesis^16,17,19–23^. Additionally, significant experience in tracking dynamic changes in sequence has been made with techniques such as kinetic sequencing, applied for example to measure the activity of ribozymes^24^.

Here, we demonstrate an approach which determines UV dose dependence of damage formation in DNA simultaneously for all 4096 possible hexamer sequences within a single experiment, only utilizing a commercial sequencing library preparation after irradiation and subsequent data analysis. Despite its simplicity, our technique reveals the bi-exponential properties of several damage states, resolves the reduction in the damage rate of di-pyrimidine lesions by nearby guanines, and provides previously unavailable damage rates for DNA lengths all the way to the hexamer. The damage is detected via the specific termination of polymerase function at the lesion site during the preparation step for next-generation sequencing (NGS). By using a suitable polymerase for damage detection, the biological relevance of the corresponding damage is thus implicitly taken into account. The relative change in frequency per sequence obtained from NGS reflects the damage process. Data analysis then reveals the sequence-dependent damaging for all possible sequences and allows to determine the respective damage quantum yields.

## Results

In order to obtain the full sequence and dose dependence of damage formation from UV-irradiation our procedure comprises five steps (see Figure 1 and Supplementary Information SI-1):

**Figure 1.**
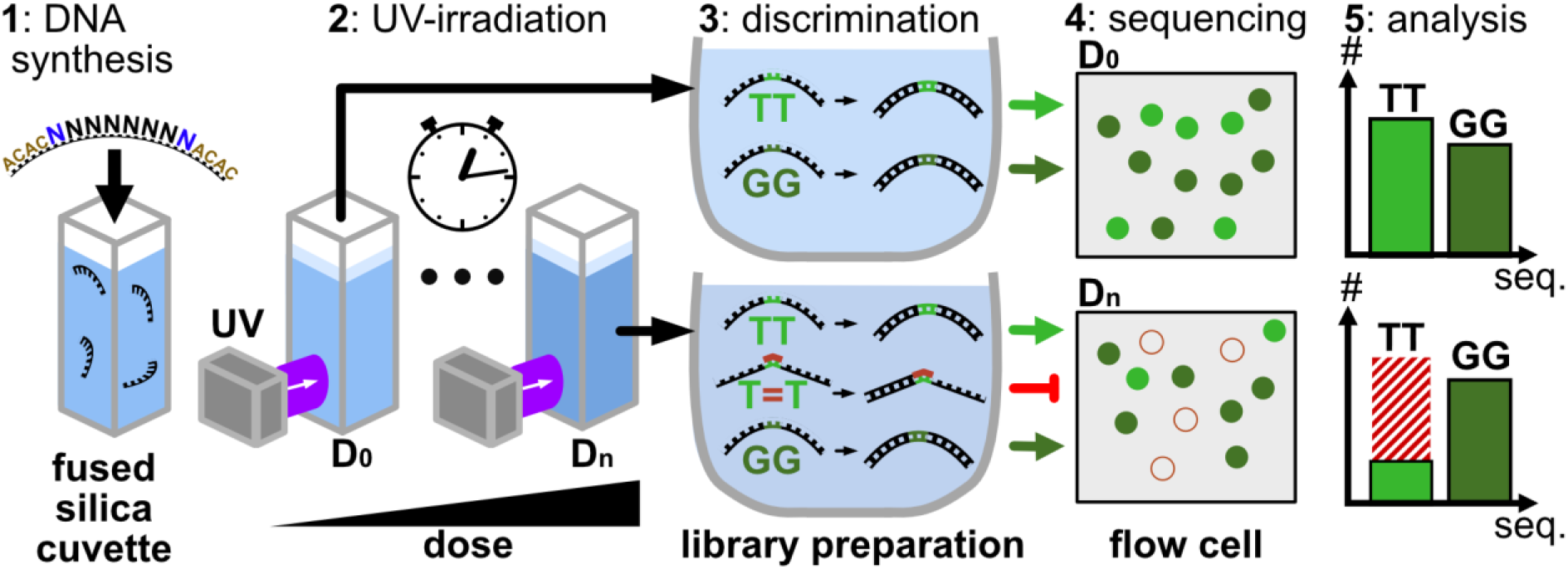
Experiment. Step 1: Short synthetic DNA strands with a central random hexamer (black) flanked by a tag sequence (brown) and spacer random nucleotide (blue) are synthesized to form a naïve sequence library. Step 2: The DNA strands in a cuvette irradiated with UV laser light at 266 nm to induce photolesions. Step 3: The irradiated sample is split into a number of samples with defined irradiation doses. Subsequent library preparation for high throughput sequencing selects and amplifies only intact sequences without UV lesions for next generation sequencing (Step 4). Step 5: Analysis of the data, normalization and correction yields the survival frequencies of submers up to hexamer length for subsequent analysis of the sequence dependent damage process.

1. *DNA synthesis*. A complete pool of oligomers with randomly distributed DNA sequences is synthesized with multiple copies of each sequence. We combine a central part, which contains the randomized nucleotides with tail sequences at the 5’ as well as at the 3’ end to facilitate analysis. In the present example the randomized part has a length of eight nucleotides and both tails have the sequence ACAC.
2. *Exposure to damage inducing process*. The randomer pool is exposed to UV-radiation (266 nm) to induce damaging photoreactions. At defined irradiation doses portions of the DNA-pool are extracted, to produce a set of samples with ascending exposure doses up to ca. 500 photons/base. The different samples are identically handled in the subsequent steps.
3. *Discrimination*. These samples undergo a discrimination step, which allows only intact DNA-oligomers to be processed in the subsequent sequencing step. In the present example, the discrimination occurs via the library preparation, which attaches pre-defined adapter strands to the DNA and amplifies the products in a polymerase chain reaction. The polymerases used in this process stop at previously generated UV lesions. Consequently, only intact strands can be successfully prepared for sequencing.
4. *Sequencing*. Only the undamaged oligomers can be successfully processed, which allows the determination of their abundance in the sequencing process. Strands, where a large fraction is damaged by the UV-irradiation will yield only small relative readout frequencies. Since the presented technique relies on the number of surviving oligomers, the sequencing procedure should be performed in a way to yield a sufficiently large number of reads for each sequence in the randomized pool.
5. *Analysis and quantification*. The frequencies of each sequence are finally analyzed and evaluated for each dose level so that the obtained two-dimensional result space allows for quantitative statements about the sequence-dependent damage. The analysis step is used to eliminate interference by possible intrinsic sequence dependencies in the synthesis, discrimination, or sequencing steps.

Due to the high number of extractions at well-defined dose values, our approach allows the investigation of complex damage models for complete sequence spaces. In the presented case, the complex sequence space covers all possible 4096 hexamers. In doing so, we can either analyse the influence of the sequence context by analysing the full strands, or by summing up all sequence frequencies of shorter sequences in the central hexamer, in order to investigate its damage properties at a higher dynamic range. Different types of DNA-damages may be addressed by an appropriate choice of the polymerase used in the discrimination step. The technique thus combines the advantages of high-throughput sequencing, the parallel measurement of damage for a complete pool of DNA sequences, with those of quantitative methods in the determination of damage processes and quantum yields.

### Detection of CPD-lesions

In a first test-experiment we studied the ability of our approach to recognize the most prominent DNA-photolesion, the cyclobutane pyrimidine dimer (CPD) between two adjacent thymidines, in which the atoms C5 and C6 from one thymine form covalent bonds to the corresponding atoms of an adjacent thymine^8,9^ (see Figure 2a). Upon absorption of photons in the dominant UV-C absorption band of DNA around 260 nm the CPD-lesion is formed predominantly via the singlet channel^5–7,11^ with a quantum yield in the range of ca. 2 · 10^−2^ ^2,4,25^. This CPD-lesion causes a distortion of single and double-stranded DNA^26,27^, which leads to a termination of the reading process for some polymerases^28,29^. The recognition of the CPD-lesion by the library preparation kit used in the discrimination step (see Materials and Methods) was tested on a number of DNA-hexamers containing a central native TT sequence or a central CPD lesion (T=T). The application of Step 3, discrimination and Step 4, sequencing, yielded the absolute frequencies of the different hexamers as given in Figure 2b. Starting from the same amount of DNA, typical absolute frequencies of 3 · 10^5^ were obtained for the intact oligomers. For all oligomers which contained T=T lesions, the absolute frequencies are strongly reduced. We found small frequencies in the range of several ten to a few hundred. Apparently, the presence of the T=T-lesion causes an approximately thousand-fold reduction of the absolute frequencies. This experiment clearly demonstrates that the procedures used in Step 3 and Step 4, namely in library preparation and NGS are able to recognize the CPD damage with high sensitivity. The molecular modifications upon formation of the T=T lesion prevent the execution of at least one step in the library preparation procedure and prohibit the readout of the original sequence^30^. The similarity of the molecular changes induced by CPD-lesions for the different di-pyrimidine lesions supports the notion that also T=C, C=T and C=C CPD-lesion are recognized by the procedure used. Here, the sensitivity of the employed technique might be deduced from a comparison of the results obtained by the present technique with literature values (see below). This approach may also be used to address the sensitivity of the presented technique towards other DNA-damages, particularly as the majority of dimer lesions alter the molecular structure of the bases significantly^25,31–33^. UV-induced strand breaks^34^ are also recognized as the fragments are rejected in the library preparation and sequencing steps 3 and 4.

**Figure 2.**
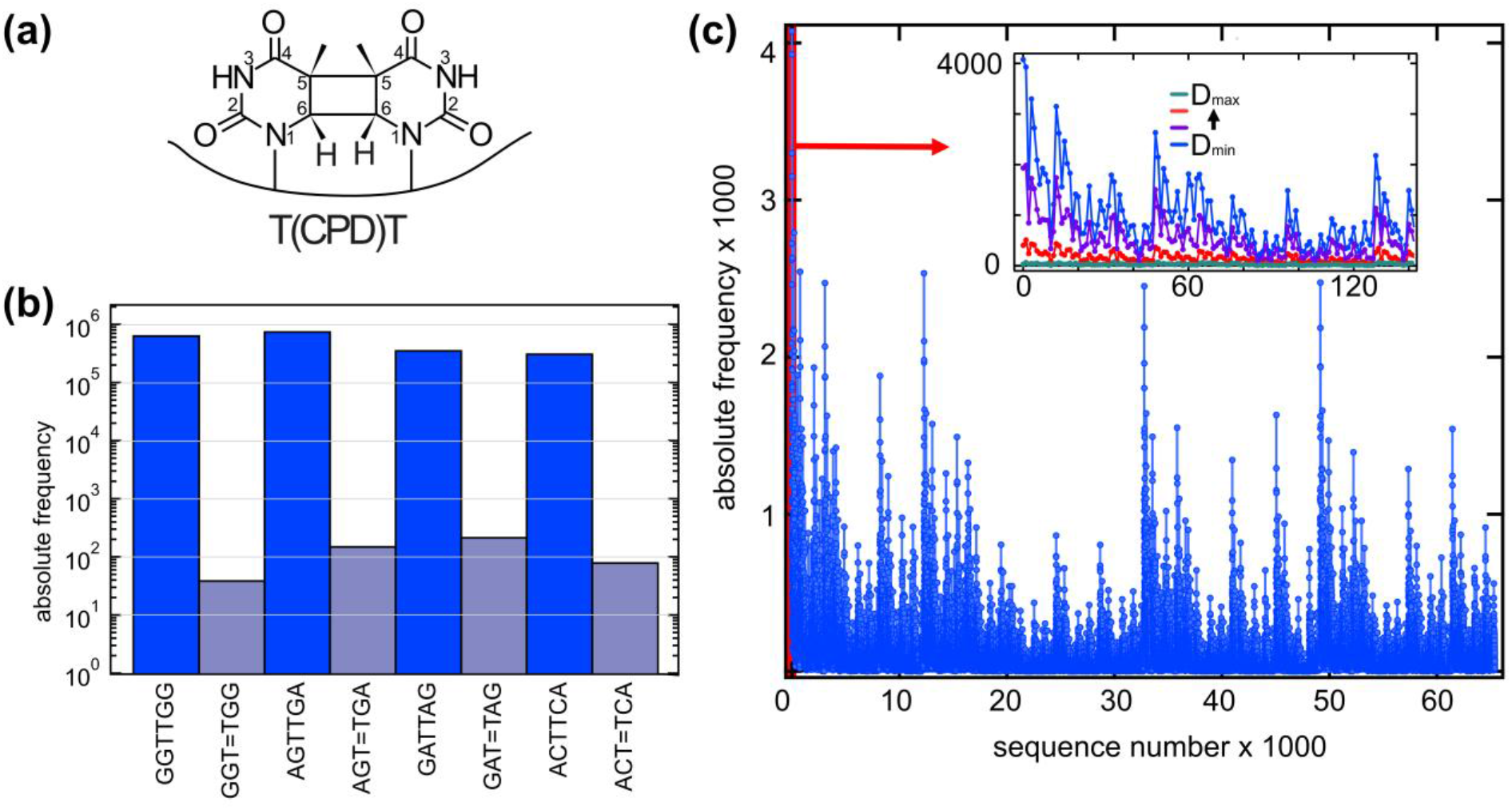
Cyclobutane Thymine Dimer (T=T) lesion and its detection by next generation sequencing. (a) Molecular scheme of the T=T-lesion with the cyclobutane ring formed with atoms C5 and C6 of the adjacent thymines. (b) Comparison of different hexamers containing an intact central TT sequence in comparison to the corresponding hexamers containing a T=T lesion at the same position. The test experiment clearly shows that the hexamers containing a T=T lesion show much smaller (more than 1000-fold) absolute frequencies than the intact hexamers. The large discrimination demonstrates that the proposed technique (Step 3 and 4) is well suited to detect the T=T damage by allowing only fully intact strands to be successfully sequenced. (c) Frequencies of the 65536 sequences from a pool of DNA strands with eight central randomers (raw data). The synthesis of the sample, the preparation of the library and the sequencing result in sequence dependencies for the unexposed sample, which must be considered in the further analysis. Insert: Frequencies for a selection of sequences taken with increasing exposure (blue: unexposed, to green: maximum dose) showing sequence dependent decrease of the abundances.

Figure 2c shows the raw sequenced reads from strands comprising the randomized octamer and ACAC tails that are irradiated by different doses of UV-radiation (Step 2). Subsequently discrimination (Step 3) and sequencing (Step 4) are performed, which yield the frequencies of the different 16-mers as raw data. These frequencies are plotted against the sequence number of the central octamer. The figure shows that the frequencies vary strongly with the sequence. For non-irradiated oligomers (D = 0, blue) the largest frequency (4075 counts) is found for AAAAAAAA while many oligomers have frequencies close to the average value of 98 count. Nearly all the 65536 possible sequences show non-vanishing frequencies. The insert of Figure 2c illustrates a narrow selection of oligomer sequences for different dose values. It is evident that the frequencies decay with increasing dose and that this decay may depend on the sequence. However, the strong variations of the frequencies of the raw data demonstrate that elaborate normalization procedures are required to obtain consistent frequencies for the different oligomers. For this purpose, we used reference sequences (poly-G oligomers with weak dose dependence). Details of the procedures for normalization and analysis used in step 5 are presented in the Supporting Information (SI-1 to SI-9, SI-15).

### Sequence Dependent Damage

The following paragraph focuses on the dependence of the frequencies of different oligomers on the applied UV irradiation dose D_j_. All presented data were obtained for ACAC-tailed random octamers of type ACAC-NNNNNNNN-ACAC after application of Step 1 to Step 5 of the measurement procedure. Since the ACAC-tails have an influence on the frequencies of the directly neighboring randomers via nearest neighbor interactions, we will only consider the frequencies of the 4096 oligomers of the central hexamer obtained upon summing over the outmost random nucleotides neighboring the tail parts. The relative frequencies of oligomers of shorter length can be calculated using the methods presented in the Supplementary chapters SI-2 to SI-4. These relative frequencies refer to the given oligomer with randomized neighbors.

In Figure 3, we compare the evaluation of polymers of different lengths. Figure 3a shows the relative frequencies (color coded) for hexamers with sequence numbers (for an example of the numbering see SI-t 3 and SI-t8, 9) between i = 3500 (TCGGTA) and i = 4095 (TTTTTT) as a function of exposure dose. The displayed sequences comprise oligomers with small decrease of frequency with dose (see e.g. around i = 3820, for oligomers containing a leading GTGT tetramer). Sequences rich in TT dimers (see i = 4095 (TTTTTT) or i = 3584 (TCTTTT), have rapidly decreasing relative frequencies which decay to < 0.2 on a dose range of 20 photon/base. Figure 3b gives quantitative information for the dose dependences of the relative frequencies for the poly-A oligomers (symbols). The mono-exponential fit functions exp(-µD_j_) (solid curves) describe well the dose dependences. The decay constants of the fit functions are μ_AAAAAA_ = 0.015 base/photon, μ_AAAA_ = 0.0098 base/photon and μ_AA_ = 0.0039 base/photon. A typical relative error of these decay constants is in the order of 20%. The inverse of these numbers indicates how many photons are absorbed in each base to decrease the relative frequency of the respective oligomer to 1/e. For the hexamer, this number is ca. 67 photons absorbed in each base. Directly related is the quantum yield Φ, i. e. the inverse of the total number of absorbed photons required to produce one damage. Since also photons absorbed in bases directly adjacent to the oligomer may damage (see SI-9), the quantum yield becomes Φ = μ_polymer_ /(i+1) = 2.1 · 10^−3^ (-AAAAAA-), 1.96 · 10^−3^ (-AAAA-) and 1.3 · 10^−3^ (-AA-). As expected, less photons are required (higher quantum yield) to damage the longer oligomers since they contain a larger number of damageable base pairs.

**Figure 3:**
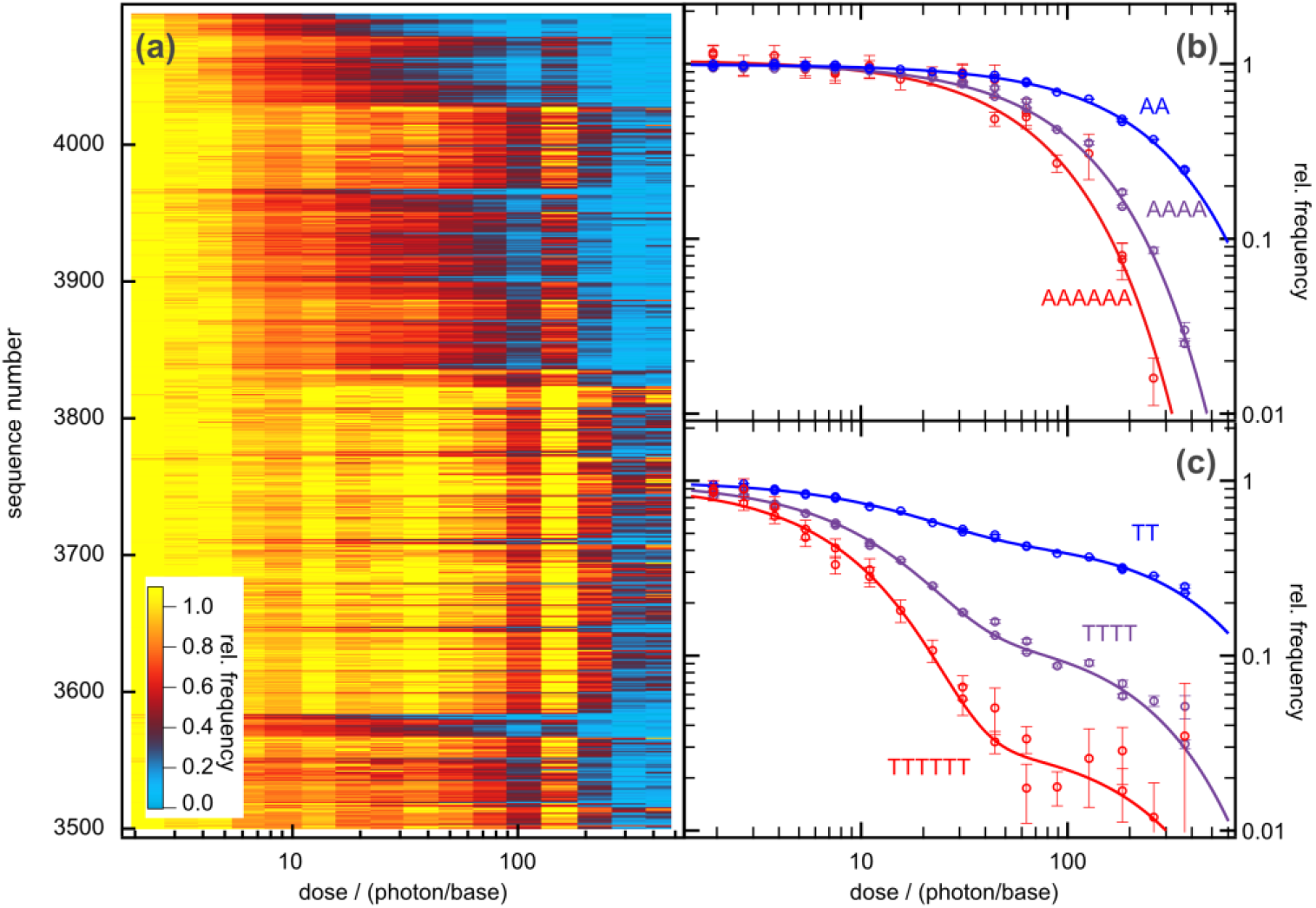
Relative frequencies as a function of UV-irradiation dose. (a) 2-D-plot for hexamers with sequence numbers between 3500 and 4095. (b) and (c) Relative frequencies for poly-A and poly-T sub-sequences as a function of UV-irradiation dose for different oligomer lengths (symbols). For poly-A (b) they follow closely mono-exponential fit-functions (solid curves). For the poly-T sequences (c) a bi-exponential behavior is observed. For poly-A and poly-T sequences the initial decay accelerates with oligomer length.

The poly-T oligomers (see Figure 3c) show a behavior clearly different from that of poly-A. The thymine sequences exhibit a much faster initial decay in the range of ten photons/base. It is striking that the oligomer frequencies do not decrease completely with the initial decay. There is a secondary slower component. While the relative amplitude of the fast decay is in the range of 0.5 for TT, it becomes much larger for the tetramer and the hexamer. From this qualitative inspection of the data, it becomes evident that the frequency of poly-T oligomers can not be described by a mono-exponential function. As discussed in the Supplementary Information SI-8, deviations from the mono-exponentiality point to a more complex reaction scheme. For instance, the photolesion may reconvert by secondary UV-absorption to the initial state (repair process) and an additional (irreversible) decay process may exist^35^. For the given precision of the experimental data, a bi-exponential fit can lead to large uncertainties in fit-parameters (amplitudes and decay constants). However, the initial slope of the relative frequency, which contains information on the damage quantum yield can often be obtained with reasonable precision.

For the initial decay of the frequencies of the poly-T oligomers we find a strong acceleration with increasing number of bases. The corresponding damage coefficients range from μ_TT_ = 0.034 base/photon, μ_TTTT_ = 0.077 base/photon to μ_TTTTTT_ = 0.11 base/photon. Thus the damage quantum yield of the poly-T oligomers becomes: Φ = μ_polymer_/(i+1) = 0.011 (-TT-), 0.0154 (-TTTT-) and 0.016 (-TTTTTT-). These numbers indicates that the damage process in poly-T is at least 10-times more efficient than in poly-A.

Figures 4 gives information on the UV-damage of tetramers. Figure 4 a to c present the dose dependence for a selection of tetramers together with fit curves. In Figure 4d a 2D-plot shows the relative frequencies of all tetramers, while Figure 4e and 4f give the data for narrow selections of sequences. The overview plot of 4d reveals ranges with very different dose dependences. There are sequences with negligible decay (see e.g. the yellow stripe around i = 170, where GT, GC or GG dimers form the tetramers). Rapid decays show up in the plot via an early appearance of a blue range. This happens for tetramers containing exclusively pyrimidines with several thymines. Details of the dose dependences are well represented by the plots on the left part of Figure 4. Figure 4a shows the results for oligomers which contain the bases T and G. For TTTT (see Figure 3b) we observe the rapid decay due to efficient damage and the deviation from the mono-exponential function which points to secondary photoreactions. For one or several thymines being replaced by guanines the survival probabilities show a much slower initial decay and the biexponentiality is less pronounced. Oligomers without the possibility to form internal bipyrimidine lesions have slower decay and show nearly monoexponential curves. The behavior found for GTGT, GGGT, and GGGG is notable: For GTGT the decay of the relative frequencies with dose nearly vanishes, while for GGGT a weak initial decay has a decay constant μ_GGGT_ = 0.0017 base/photon. Interestingly, the poly-G oligomer with μ_GGGG_ = 0.00077 base/photon (value determined by the normalization procedure, see SI) lies in between.

**Figure 4:**
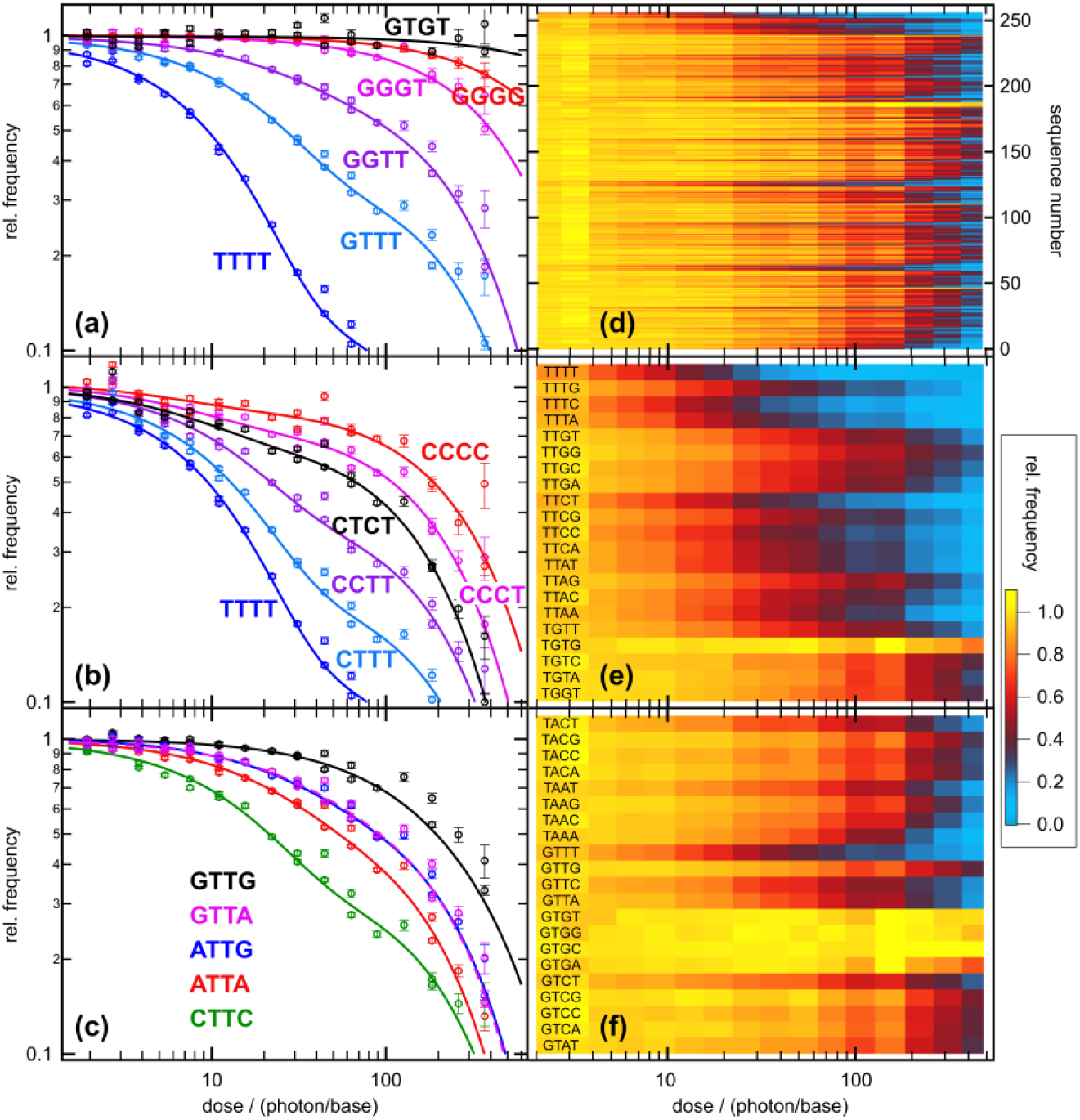
Relative frequencies of tetramer sequences as a function of UV-irradiation dose. (a) Sequences from TTTT to GTGT show the large range of decay constants. When several TT-dimers in the sequence exist, the possibilities to form T=T lesions lead to rapid decay of the survival probability. (b) For combinations of thymine and cytosine bases in the tetramers there is always the possibility to form several CPD-lesions. There are clear indications for a bi-exponential behavior. (c) Relative frequencies for a selection of tetramer sequences containing a central TT dimer plotted as a function of UV-irradiation dose. When the TT-dimer is flanked by purines A or G the decay is much slower than for flanking C, where CPD-lesions are possible between all bases of the tetramer. The decay of GTTG is surprisingly slow, presumably due to the possible charge-transfer reaction between G and T, acting as an additional decay channel reducing damage frequuency. (d) 2D-plot of the relative frequencies for all 256 tetramers. (e) and (f) expanded views for tetramers with a large fraction of fast decaying sequences (e) containing TT dimers and pyrimidine trimers. When GT or GC dimers are present (f) the decay is very slow.

Different combinations of the pyrimidine bases thymine and cytosine are included in the oligomers shown in Figure 4b. The poly-C data reveal a weak bi-exponentiality where the initial decay has only a small amplitude. The dominant part of the damage occurs at higher doses. In the mixed oligomers the most efficient damages occur when at least one TT-dimer is present. However, combinations of C and T without TT or CC dimers also lead to considerably fast lesion formation. For a comparison of oligomers showing more or less pronounced bi-exponentialities we consider the dose values D_50%_ where 50% of the oligomers are damaged. These dose values decrease from D_50%_ = ca. 190 (CCCC), via 110 (CCCT), 64 (CTCT), 21 (CCTT), 13(CTTT) to 9.6 photon/base (TTTT).

In Figure 4c we compare tetramers with a central TT sequence. When TT is flanked by C, pyrimidine dimer lesions become possible between TT, CT and TC and lead to a rapid decay of the survival probability. Only ca. 21 photons/base are required to damage 50% of the CTTC tetramers. When purine bases are flanking the central TT dimer, there is a considerable reduction of the damage efficiency. For ATTA D_50%_ becomes 53 photon/base. With flanking G the damage resistivity improves even more. For GTTG, we observe a much higher damage resistivity D_50%_ = 200 photon/base. The oligomers ATTG and GTTA have similar survival probabilities with D_50%_ = 87and 92 photon/base, respectively.

Figure 3 and 4 contain the information on the dose dependence for a limited number of tetramers. An overview of all tetramers is given in the Supporting Information in Table SI-t17, where the total damaging coefficients μ_olig_ and the related quantum yields are given together with the D_50%_ values for all 256 sequences of the tetramer pool. Discrepancies between D_50%_ and D_50%,mono_ = ln2 /μ_olig_ occur for sequences with strong deviations from the simple mono-exponential decay.

One remark should be added regarding the precision of the deduced damage coefficients, which amounts to ca. 25% relative error. One source for these uncertainties is the statistical error of the relative frequencies of the fit procedure. In most cases the uncertainty is dominated by the limited precision in the determination of the molecular concentrations and thus of the number of absorbed photons per base.

### Molecular Damage and Quantum Yield

The experimental results on the dose dependence of the oligomer frequencies, e. g. the initial decay constants μ_olig_ give important qualitative insight into the role of the sequence for the initial steps in UV-damage formation. Quantitative information on the underlying molecular processes and a comparison with the quantum yields given in the literature may be obtained when combining the observations with a molecular model for damage formation. Here, we use a simple model of the initial step of damage formation, which is based on dimeric damage processes and nearest neighbor interactions (see scheme in Figure 5, insert and SI-9 for details). We assume the same excitation of all bases in the strand, where the excitation of one base X in a sequence WXY is equally shared with the two neighboring bases W and Y leading to excited dimer states (WX)* and (XY)* which cause damages of the respective dimers with the dimeric quantum yields φ_WX_ and φ_XY_.

**Figure 5:**
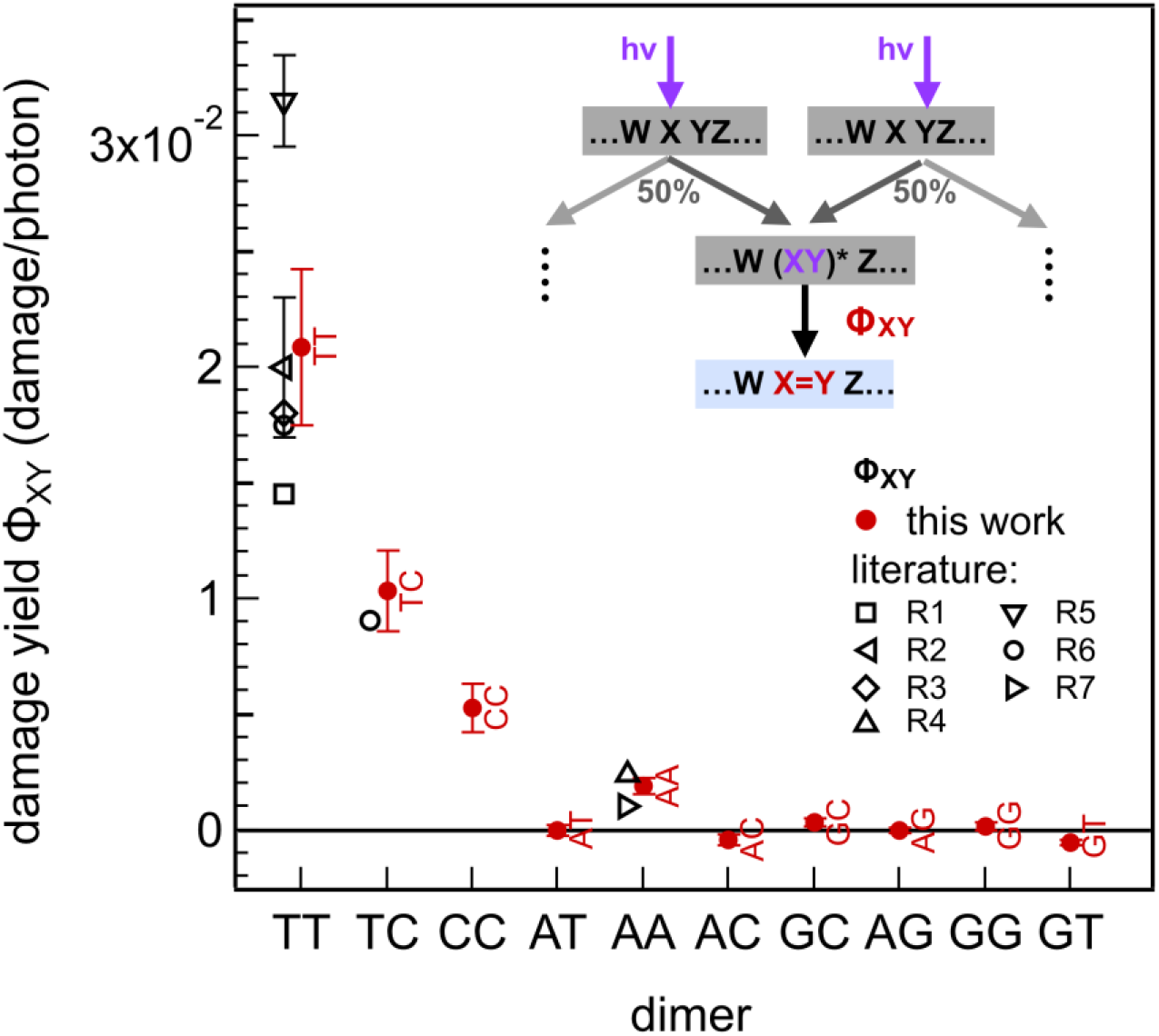
Quantum yields for the damage determined according to the molecular model with dimeric damage processes (see insert and Equation 1). The quantum yields (red dots) agree well with the literature (black, open symbols) for the bi-pyrimidines: R1:^2^, R2:^4^, R3:^12^, R4:^36^, R5:^37^, R6:^25^, R7:^33^. For most purine containing dimers only small values of the damage quantum yields are found below the experimental sensitivity of ca. 0.5 10^−3^.

From the tetramer data we obtained a set of quantum yields with values for the strongly damaging di-pyrimidine dimers in agreement with literature. However, for the guanine containing dimers GT and GC negative values for the quantum efficiencies, e. g. μ_GT_ = −5 10^−3^, resulted which require an extension of the molecular model by processes which reduce the formation of di-pyrimidine dimer lesions when G is adjacent. One explanation could be that the formation of charge transfer states G^+^T^-^ or G^+^C^-^ between the strong electron donor G and a pyrimidine deactivates the damage^7,9,38–43^. In a very recent study the role of charge transfer quenching has been studies for tetramers^44^.

From this complete model one obtains the dimeric quantum yields given in Figure 5, filled circles. The most efficient damages occur for the pyrimidine dimers. The largest damage quantum yield is found for the TT dimer with *Φ*_*TT*_ = 0.021 ± 0.003. For CT (or TC) a somewhat smaller value is obtained, *Φ*_*CT*_ = 0.011 ± 0.002. For CC the quantum yield is again smaller, *Φ*_*CC*_ = 0.006 ± 0.001 and shows a considerable uncertainty. The purine dimer AA has a quantum yield around *Φ*_*AA*_ = 0.0019 ± 0.0003. These values are within the ranges found in the literature (see Figure 5 open symbols). For the other purine containing dimers quantum yields are in the range of < 0.5 · 10^−3^. In other words, they are zero within the present precision of the experiment. The analysis of the frequencies from the dimer pool leads, within the given uncertainties, to the same quantum yields. Since the frequencies in the hexamer pool are much smaller than for dimers and tetramers, larger uncertainties arise and prevent a reliable bi-exponential fit, the prerequisite for a successful analysis.

## Discussion

In this work we measured the dependence of oligomer survival on UV-exposure for the complete sequence space of DNA hexamers. Our measurements reveal the sensitivity of the employed technique for dimeric DNA lesions (see Figure 2). For complete hexamer, tetramer and dimer pools the sequence dependent survival was determined and results are presented in Figures 3 to 5, and SI-t 8 and SI-t 9. There are oligomers with high UV-resistivity, where even dose values of > 2000 absorbed photons per hexamer (or 500 photon/base) scarcely harm the survival. These oligomers often contain a large fraction of guanines and single pyrimidines (see Table SI-t 9) but no TT dimers. For oligomers with longer pyrimidine parts, especially poly-T parts, even small doses in the range of 10 photon/base are sufficient to damage most of the oligomers. The role of pyrimidine dimers for the DNA-damage becomes evident when the data are analyzed using the simple molecular model presented above. It reveals that the most important damage quantum yield in the range of 2% occurs for TT dimers. The quantum yields for CT (TC) or CC-dimers are somewhat smaller with ca. 1% and 0.5%, respectively. These values agree with the literature, proving that the presented technique with the discrimination step using the Swift Library Kit (see Materials and Methods) is able to detect the CPD lesion. Most of the purine containing dimers have quantum yields below our detection limit of ≤10^−3^. These small values are again in agreement with the literature (TA^45^, GC^31^, AA^36^). Only the AA-dimer shows a detectable damage quantum yield of ca. 0.2% in the range of the literature values. Thus, the used library preparation kit is also able to unravel AA damage sites. When we inspect the presence of A in the hexamers showing strong irradiation damage (see Table SI-t 9, lower part) we often find poly-A parts. Apparently longer poly-A parts can be taken as an indicator for increased susceptibility towards UV-irradiation damage.

The DNA in our experiments was mainly single-stranded. By a special design of the tail sequences and by suitable temperature and salt conditions it would be possible to increase the probability for double strand formation and to record the corresponding UV-sensitivity. Further interesting extensions of the presented technique could involve the study of damage induced by other irradiation wavelengths, by photosensitization or to investigate the action of denaturing and reactive compounds.

The sensitivity of the technique depends on the readout possibilities of the sequencing procedure, because it is based on the frequency of surviving oligomers. This requires high count numbers for all sequences of the oligomer pool, restricting the maximal strand length. In future experiments, longer sequences could be addresses by strategies combining randomized parts with parts of defined sequences, to limit to total length of the randomized parts.

The integrity of the genetic information of organisms living today is maintained by sophisticated enzymes. Our approach allows to examine damages of single stranded DNA without the need for additional enzymatic repair processes. The resulting information on irradiation-induced damage represents situations with (i) UV-exposure by high-intensity irradiation, where severe damage is established prior to the start of repair, (ii) for organisms with defective repair or (iii) for short DNA strands of early (molecular) life^46^, before the establishment of efficient enzymatic repair mechanisms. Because of the major role of RNA in the origins of life, the extension of our technique to RNA opens promising perspectives. Investigation of UV irradiation-induced damage of RNA strands would require an adaptation of the discrimination Step 3 by using reverse transcriptases. This modification appears to be straight forward since suitable RNA library preparation kits exist for NGS. The potential for the recognition of RNA damages remains to be investigated.

**In conclusion**, we characterized the sequence dependent irradiation damage in large, randomized pools of DNA in a highly parallel fashion. Our approach combines different standard processing steps, such as synthesis of oligomers with randomized sequences, UV-irradiation, discrimination against damaged DNA strands and the identification of the intact oligomers by NGS with appropriate analysis. Using DNA-strands with centrally randomized octamers, we could characterize the sensitivity towards UV irradiation of the complete sequence pool up to hexamers which consists of 4096 different sequences. The evaluation of the damage response by means of a simple model with dimeric damage formation was used to obtain the damage quantum yields of bi-pyrimidine dimers in agreement with the literature. However, the comparison of the dimeric model with the experimental data also revealed a protecting action of the sequence context. For example, the rate to form TT lesions is strongly reduced by a nearby guanine. In this work, the proposed technique was demonstrated on the radiation-induced lesion formation of single stranded DNA. Extensions of the method while keeping the underlying principles are possible to study other DNA or RNA structures or to record the response of the nucleotide strands upon exposure to damaging reagents.

## Materials and Methods

DNA 16-mers with random parts of the sequence ACACNNNNNNNNACAC were synthesized by Biomers, (Germany), dissolved in PBS-buffer and diluted to a base concentration of ca. 0.7 mM (concentration of bases, checked by UV-absorption, Shimadzu UV1800). Oligonucleotides containing the CPD-damage T=T were purchased from IBA GmbH, Germany. A starting volume of 3.45 ml of the 16-mer sample was irradiated at 266 nm pulses from a Nd-based laser system (AOT-YVO-25QSP/MOPA from Advanced Optical Technology, UK) in a fused silica cuvette (path length 10mm) under stirring. Monitoring of the laser power allowed to determine the power absorbed in the sample (for details of the determination of the absorbed dose see SI-7, relative error in the determination ca. 16%). At each dose level 50 µl sample was taken from the cuvette and stored at low temperatures for further handling. Subsequently the samples were prepared for sequencing (Step 3) according to the provided protocol of the Swift Accel-NGS 1S DNA Library Kit (Swift Bioscience, US). The sequencing (Step 4) was performed using a Hi-Seq sequencer (Illumina, US) and the data was then filtered using the framing sequence ACAC with a quality score larger then 20 and analyzed (Step 5) using the methods given in the Supplementary Information SI-1 to SI-14. The main parts of data analysis involve normalization with respect to the unirradiated sample and to the oligomer containing poly-G in the central part. Further analysis steps involve the numerical fitting of the data by bi-exponential functions and the computation of the dimeric quantum yields via the molecular model described in SI-9. Further details on the methods used can be found in SI-1.

## Acknowledgements

This work was supported by the Deutsche Forschungsgemeinschaft (DFG, German Research Foundation) through the Clusters of Excellence “Center of Integrated Protein Science Munich” (W.Z.) and Project-ID 364653263 – TRR 235 (CRC235), Project P08 (C.B.M). Funding from the Simons Foundation (327125 to D.B.), Volkswagen Initiative ‘Life? – A Fresh Scientific Approach to the Basic Principles of Life’ (C.B.M., D.B.), ERC ADV 2018 Grant 834225 (EAVESDROP) (D.B.) and from ERC-2017-ADG from the European Research Council (D.B.) is gratefully acknowledged. Funded by the Deutsche Forschungsgemeinschaft (DFG, German Research Foundation) under Germany’
ss Excellence Strategy – EXC-2094 – 390783311 (D.B., C.B.M). The work is supported by the Center for Nanoscience Munich (CeNS).

## Author Contributions

CLK, SK, MF, JPM, WZ, CBM performed the experiments. DB, CLK, DBB, HB, WZ, CBM conceived and designed the experiments. WZ, CBM analyzed the data. CLK, DBB, DB, WZ, CBM wrote the paper. All authors discussed the results and commented on the manuscript.

## Competing interests

The authors declare no competing interests.

## Data availability

The data supporting the findings of this study are available within the paper and its Supplementary Information. Additional information and files are available from the corresponding author upon reasonable request.

## Code availability

The operations discussed in the Supplementary Information and Materials and Methods for evaluating and analyzing the data have been implemented using standard methods in Labview and Igor Pro and are available on request at any time.

## Notes

### Competing Interest Statement

The authors have declared no competing interest.

