## Supplementary Text for "Sequence Dependent UV Damage of Complete Pools of Oligonucleotides"

### Supplementary Information

#### Table of contents:

**Chapters**:

SI-1: Methods and Materials
SI-2: Normalization of the raw data
SI-3: Improving statistics by averaging over tail sequences and positions
SI-4: Exponential length dependence of the survival probability
SI-5: Validation for sequence-averaging over all positions
SI-6: Sequence symmetry of survival probabilities
SI-7: Absorbed Dose
SI-8: Molecular Model for Dose-Dependence
SI-9: Molecular Model for the Damage Coefficient of Oligomers
SI-10: Fitting of damage rates
SI-11: Error propagation
SI-12: Example for data handling
SI-13: Detection of outliers
SI-14: Analysis for the predominant occurrence of single-stranded DNA
SI-15: Determination of the damage yield for GG-dimers by ultraviolet spectroscopy
SI-16: Assessment of the sensitivity of the method for detection of 8-oxoguanine
SI-17 Characteristic data on the dose dependences for tetramers and hexamers

**Figures**:

SI-f 1: Raw data from sequencing.
SI-f 2: Raw data from sequencing with dose dependence.
SI-f 3: Plot of the averaged absolute frequency of the 16-mer data versus dose
SI-f 4: Factorization of survival probabilities according to equation SI-e 12
SI-f 5: The result of Equation SI-e 17
SI-f 6: The sum shown in the left-hand side of Equation SI-e 20
SI-f 7: The deviation of isolated survival probabilities of dimers
SI-f 8: Comparison of survival probabilities of sequences and their reverse counterparts
SI-f 9: Different reaction models used in the description of the UV-induced damage
SI-f 10: The experimentally obtained isolated survival probabilities for dimers
SI-f 11: Data for the selection of outliers
SI-f 12: Calculated statistics on the binding probability in a randomized DNA
SI-f 13: Plot of the absorption spectra of the GG-dimer sample in the near UV and visible range for different illumination values
SI-f 14: Change in the relative frequency of a target sequence ACAC AGAGAG ACAC upon insertion of an 8-oxo-guanine at position 8

**Tables**:

SI-t 1: Values of extinction coefficients used for the calculation of the absorbed dose
SI-t 2: Absolute frequencies for the raw data set
SI-t 3: Absolute frequencies of the central hexamer
SI-t 4: Absolute frequencies of the dimers
SI-t 5: Relative frequencies of the dimers

SI-t 6: Relative frequencies of the dimers
SI-t 7. Relative frequencies of the dimers after correction
SI-t 8. Decay parameters for tetramers
SI-t 9. Hexamer sequences with largest and smallest survival probabilities

#### SI-1. Methods and Materials

##### Description of the method

The proposed technique aims to quantify the sequence dependent response of oligonucleotides to external stimuli like UV-irradiation (the focus of this paper), which modify or damage the molecular structure of the nucleic acids. The technique allows the simultaneous characterization of a wide variety of sequences by high-throughput quantification. The method uses the combination of five work steps.

**Step 1: DNA-Synthesis**: A pool of randomly distributed DNA sequences of length n and comprising all 4 canonical bases was created. During every extension step of the synthesis, all four nucleotide types were provided at equal concentrations. Deviations from a completely uniform base distribution are corrected in the data analysis in step 5.
At the 5' and the 3' ends, the random sequences are flanked by defined tag sequences, which make it possible to identify intact oligomers against poorly prepared sequences and sequence fragments present in the sample after the end of the preparation steps. In addition, the tag sequences may be designed to cause the formation of complexes or as done in this work, to ensure the DNA to remain single stranded. In the present investigation of the UV-sensitivity of DNA-oligomers we used n = 8 randomized bases flanked by ACAC sequences on both ends, yielding sequences of the pattern 5'-ACAC-NNNNNNNN-ACAC-3' (N is any one of the 4 canonical dNTPs: A, G, C, T).

**Step 2: Exposure to damage inducing process (1$^{st}$ dimension, here UV-irradiation**). The pH-buffered sample solution is irradiated for a defined time at constant temperature while the radiation intensity is continuously monitored to achieve the desired radiation dose $D_j$. After each irradiation process, a part of the solution (sample j) is extracted and stored for analysis at -80°C while the rest of the sample is further irradiated. Subsequent irradiation steps lead to a set of samples with defined irradiation doses. The irradiation dose is defined here as the average number of UV-photons absorbed per base in the sample j. The applied dose was up to 500 photons per base to record damage quantum yields down the $10^{-3}$ range. The dose dependent alteration of the oligomers with different sequences will be visualized in the subsequent work steps.

**Step 3: Discrimination.** The most critical step is the quantitative detection of oligomers that have been altered by the UV-irradiation in the previous step. To avoid time-consuming chemical analysis of the UV-damaged strands, we determined the sequences of all intact oligomers before and after exposure using the inbuilt polymerase stop functionality implemented by the high-fidelity polymerase included in the library preparation process. From the difference of reads per sequence between these two measurements, the frequency of damaged strands can then be deduced. In principle, this step can also be performed with any other technique that can select for intact strands prior to sequencing. However, since commercially available library preparation kits for Next-Generation Sequencing (NGS) use high fidelity polymerases, they are expected to be sensitive for major structural and chemical modifications of the DNA strands[1]. In the present experiments we used these inbuilt stopping capabilities instead of adapting the library preparation Kit to contain polymerases with known error detection rates. In further investigations the sensitivity of the enzymes used to detect the desired type of damage should be studied. After the discrimination step, only undamaged DNA should be successfully indexed for sequencing.

**Step 4: Sequencing, determination of the oligomer frequency (2nd dimension).** Only completely prepared DNA, i.e. strands with fully ligated index adapters, can be detected quantitatively by NGS. More frequent sequences generate more readout events, which allows the determination of the relative number of all sequences within a sample. Since the technique relies on the number of surviving oligomers, the sequencing procedure should be performed in a way to yield a sufficiently large number of counts for the different sequences in the pool.

**Step 5: Analysis and quantification.** The different steps in sample preparation, amplification and analysis may be subject to fluctuations in relevant parameters such as pipetting error or sequence dependence of used enzymes. These effects alter the total and relative number of oligomers with certain sequences. In addition, the synthesis, library preparation and the sequencing itself can be sequence dependent. These effects have to be considered before the dose dependence of the survival of oligomers can be quantified. An example is given in Figure SI-f1, where the average number of oligomers is plotted for the unirradiated sample, i.e. $D = 0$, without any corrections. The observed frequencies vary by several orders of magnitude with a preference of oligomers containing mainly adenosine. To correct these variations, the frequencies of the sequences from the irradiated sample should be normalized with the corresponding frequencies of the sequences from the unirradiated sample (see next Chapter). Sequence-independent deviations between different samples can additionally be corrected by further normalization to a supposedly known frequency of a certain "lighthouse sequence". This could be, for example, a sequence that is not damaged by UV light at all, or whose damage rate could be determined by an independent measurement using e.g. UV spectroscopy. Details of the correction process and the underlying equations are presented in the next chapter.

**Materials and techniques**

DNA randomers of the sequence ACACNNNNNNNNACAC and 8-oxoG modified sequences (see SI-16) were synthesized by a commercial provider (Biomers, Germany), whereby all four canonical nucleotides were used in equal amounts in the synthesis steps of the central 8-mer. Upon receipt, the samples were dissolved in 1x PBS (137 mM NaCl, 2.7 mM KCl, 10 mM Na2HPO4, and 1.8 mM KH2PO), diluted to a base concentration of 1 mM, vortexed, and centrifuged. The final DNA concentration was checked by Nanodrop (Thermofischer, US) and UV-vis measurement (Shimadzu UV1800). Due to the low absolute concentration per sequence of 1 nM and the fact that Watson-Crick base pairing can occur mainly in the comparably short central 8-mer, less than 0.1 % DNA duplexes are expected at room temperature. Therefore, the expected effect of reducing the susceptibility to damage in the double strand compared to a single strand is negligible (see SI-13).

Oligonucleotides GGT=TGG, AGT=TGA, GAT=TAG and ACT=TCA containing the CPD-damage T=T were purchased from IBA GmbH, Germany. These oligonucleotides (and the corresponding intact oligomers at the same concentrations) are directly introduced in Step 3 of the procedure and the absolute frequencies of the intact sequences are recorded after Step 4.

For the UV-exposure, 3.45 ml of the sample were filled into a fused silica cuvette (Type 117100F-10-40, Hellma, Germany) with a pathlength 10 mm, which was then placed in a temperature-controlled socket. Magnetic stirring was used to ensure a homogeneous sample at a temperature of 22°C. The exposure itself was carried out with a Nd-based laser system (AOT-YVO-25QSP/MOPA) from Advanced Optical Technology, UK, (repetition rate of 6.5 kHz, average power ca. 20 mW, wavelength 266 nm, beam diameter in the sample ca. 1 mm). The laser power before and after the sample cuvette was monitored by power meters (Ophir, Israel) to determine the power absorbed in the sample (for details of the determination of the absorbed dose see SI-7). After a desired dose was reached, the exposure was stopped briefly and 50 µl sample was taken from the cuvette, frozen in and stored at -80°C. Illumination and sample collection was repeated until a maximum average dose of ca 500 photons per base was reached. The samples were then prepared for sequencing according to the protocol of the Swift Accel-NGS 1S DNA Library Kit (Swift Bioscience, US), taking care to process the samples immediately after thawing. The sequencing was then performed using a Hi-Seq sequencer (Illumina, US) and the data was then analyzed using the methods given in the main text and in the Supplementary Information SI-1 to SI-14.

The determination of the dose, i.e. of the number of absorbed photons per base (see SI-7) requires knowledge on the number of oligomers in the sample cell and on the extinction coefficients of the different bases in the oligomers at the irradiation wavelength of 266 nm. For this wavelength the extinction coefficients of the four bases A, C, G and T are similar. For simplicity of the data evaluation we assume that the extinction coefficients are the same. This assumption together with the uncertainty of the determination of the absolute extinction coefficients leads to a relative error in the determination of the irradiation dose of $\Delta D/D = 0.16$. When decay coefficients of the fit curves or quantum yields $\Phi_{XY}$ are determined, the total error is due to the error of the fit procedures and the error in the determination of the dose values.

#### SI-2. Normalization of the raw data

The presented technique in the main document and in SI-1 records the action of an external exposure - in the present case UV irradiation - on oligonucleotides in a highly parallel way. The parallel nature is obtained by combining oligomers with defined length but arbitrary sequences and second-generation sequencing leading to a 2D-data set. Details of the different steps of the data handling procedure are (see also main text and SI-1):

(1) DNA oligomers of a defined length with tails of (sequence ACAC, in some cases GTTG) at the 5' and the 3' end contain in the center (eight) bases with arbitrary sequences. I. e. the DNA synthesis is performed with equal concentration of the four nucleotides A, C, G or T. Possible sequence biases during synthesis will be addressed (see below). The tails are used to identify the intact oligomers after the sequencing process. In addition, they may serve to avoid double-strands in the oligomer solutions. The sequence ACAC for the tails is chosen since it contains no damage-prone bi-pyrimidine and should survive UV-irradiation doses of considerable strength. The complete 16-mer solution is split after the irradiation into $J = 18$ individual samples. These sample have the same volume and oligomer concentration. For an arbitrary sequence i, within the precision of the synthesis and the splitting process, they should contain the same number $H^{(0)}(i)$ of oligomers with central sequence $i$.

(2) In the second step of the experiment the different aliquots are treated by specific amounts of UV-irradiation. In this process a fraction of the oligomers undergoes radiation damage, depending on the absorbed dose $D_j$ (in our case dose is defined as absorbed photons per base). In this process the number of intact oligomers with sequence i in sample j decreases due to absorption of dose $D_j$ to $H^{(1)}(i,j)$

(SI-e 1) $$H^{(1)}(i,j) = S(i,D_j)H^{(0)}(i)$$

Here $S(i,D_j)$ is the survival probability of oligomer $i$ exposed to dose $D_j$. This quantity contains the information on the dose dependence of the oligomers with sequence $i$.

(3) In the subsequent treatment steps (library preparation leading to discrimination), intact oligomers are transformed into structures which enter the subsequent amplification and analysis steps (PCR and bridge amplification) of the Illumina process. The damaged structures are assumed to be rejected in these processes. The parameters of rejection and amplification may vary from sample to sample leading to an efficiency factor $p_j$, which is assumed to be independent of the specific sequence. Simultaneously the amplification steps may also depend on the sequence, which is considered by the sequence dependent efficiency factor $q_i$ and is the same for all samples. Thus, the final number $H^{(F)}(i,j)$ of intact oligomers after amplification should become:

(SI-e 2) $$H^{(F)}(i,j) = H^{(1)}(i,j) \cdot p_j \cdot q_i = S(i,D_j) \cdot H^{(0)}(i) \cdot p_j \cdot q_i$$

It should be noted that $H^{(F)}(i,j)$ refers here to the complete 16-mer with eight bases in the tails and the central randomer. The sought-after quantity of the investigation is the survival probability as a function of the sequence and the dose, $S(i,D_j)$. However, the efficiency factors $p_j$ and $q_i$, which may exhibit considerable variations with $i$ and $j$, also enter Equation (SI-e 2). They may hide the relation between the original oligomer frequencies $H^{(0)}(i)$ and $S(i,D_j)$. Thus $S(i,D_j)$ can only be obtained after elimination of $p_j$, $q_i$ and the sequence dependence of original oligomer frequencies $H^{(0)}(i)$ by appropriate normalization procedures. The pronounced variations with $i$ and $j$ are evident from the plot of the recorded data $H^{(F)}(i,j)$ plotted versus index $i$ representing the sequence of the central octamer (Figure SI-f 1 or Table SI-t 2).

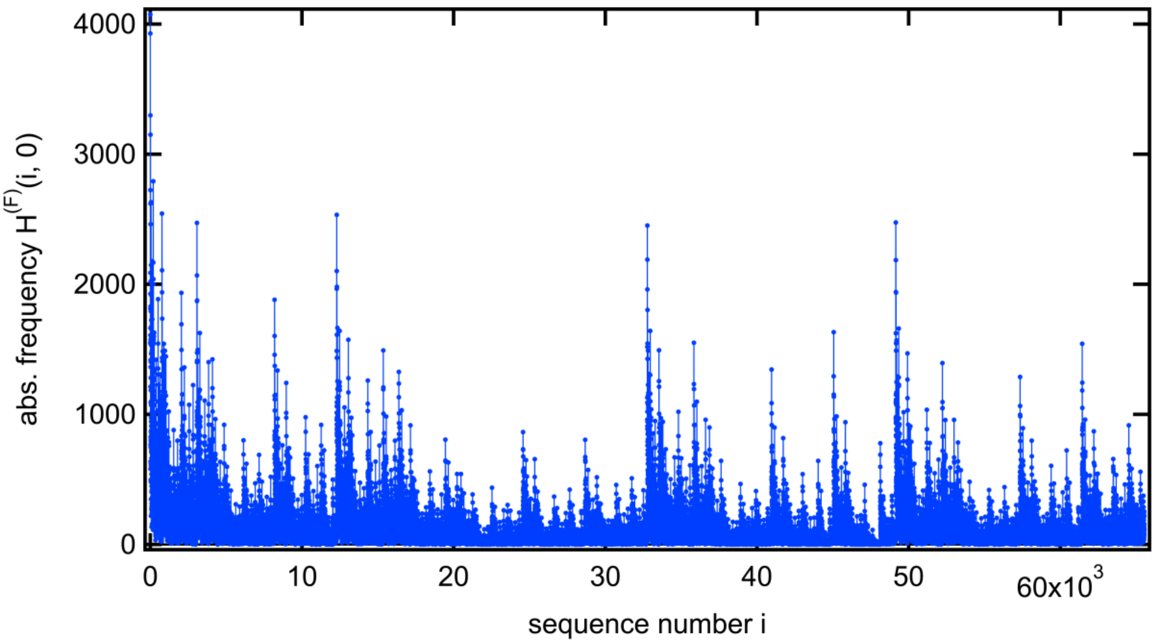

**Figure SI-f 1.** Raw data $H^{(F)}(i,0)$ versus sequence of the central octamer for the sample with dose $D = 0$.

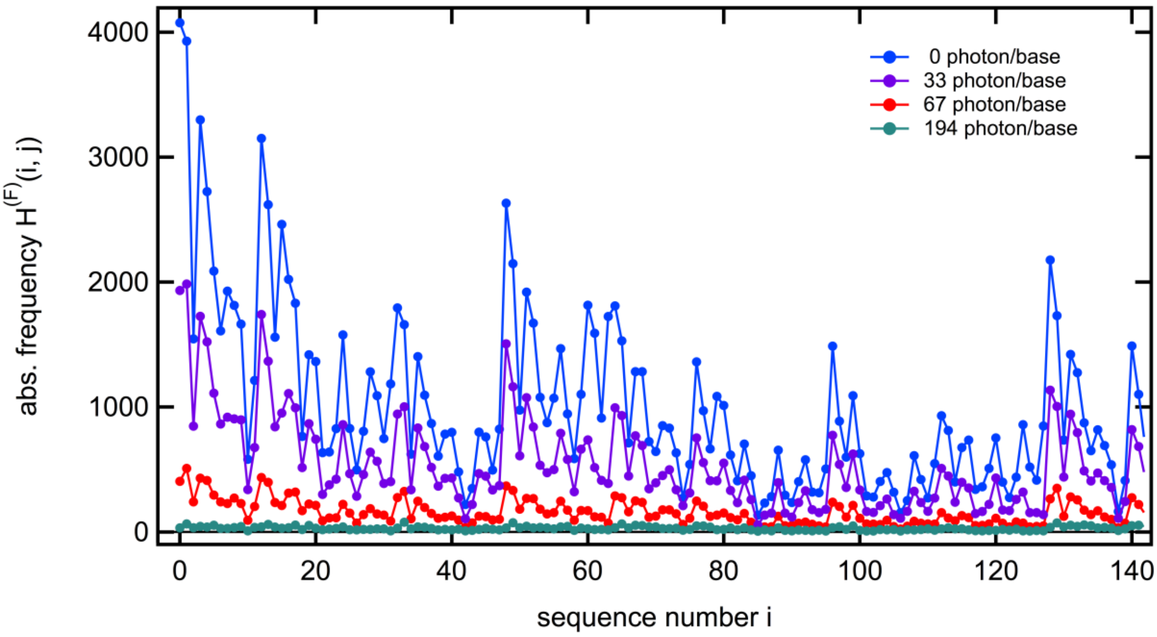

**Figure SI-f 2.** Raw data $H^{(F)}(i,j)$ for small values of the sequence index $i$ for samples with the dose values $D_j$ given in the Figure.

For the not-irradiated sample (Figure SI-f 1 and Figure SI-f 2 blue trace), one observes strong variations of the recorded numbers $H^{(F)}(i,0)$. A similar pattern of the sequence dependence is found for other samples. These strong variations may be related to sequence dependencies of the term $H^{(0)}(i) \cdot q_i$. For higher doses values the term $S(i,D_j)$ may also gain importance.

Samples subject to remarkably high irradiation doses show the expected strong reduction in oligomer numbers (Figure SI-f 2). This damage can be caused by the central, randomized bases as well as by the bases in the tails. In order to demonstrate the dose dependence, we plot in Figure SI-f 3 the absolute frequencies averaged over all sequences as a function of the applied dose. The data points at the higher dose values (red squares, lower scale) clearly show a strong decay with increasing dose values. The over-all decay is not mono-exponential and occurs on the dose-range of 50 photon/base. The points at smaller doses values (violet open squares, top scale) exhibit that in addition to the general trend (decay with increasing dose) there are fluctuations, which may result from the variations in the preparation/analysis conditions for the different samples, represented by the efficiency factor $p_j$. These results indicate that normalization procedures are required to obtain the correct survival probability $S(i, D_j)$ from the recorded data $H^{(F)}(i,j)$.

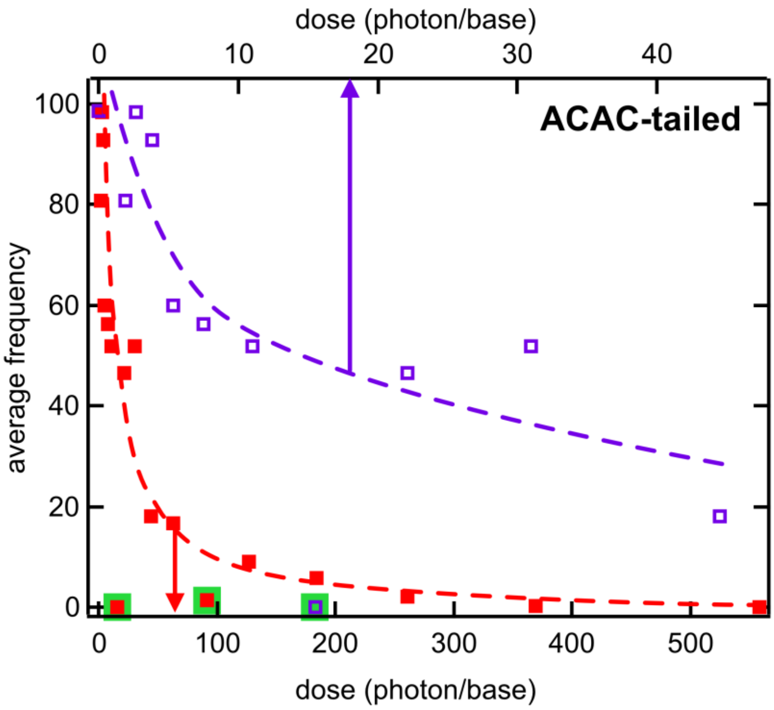

**Figure SI-f 3.** Plot of the averaged absolute frequency of the 16-mer data versus dose. Red squares, lower scale, violet open squares, top scale. Outliers marked with green. The data clearly show the overall decay of the frequency with dose together with fluctuations around the general trend due to preparation/analysis artifacts.

For the elimination of the general sequence dependent efficiency factor (product $H^{(0)}(i) \cdot q_i$) we divide the recorded oligomer numbers $H^{(F)}(i,j)$ by the oligomer numbers obtained at dose $D_0 = 0$, where the survival probability is, by definition, set to $S(i,0) = 1$.

(SI-e 3) $$H^{(Norm-i)}(i,j) := \frac{H^{(F)}(i,j)}{H^{(F)}(i,0)} = \frac{S(i,D_j)H^{(0)}(i)p_j q_i}{H^{(0)}(i)p_0 q_i} = \frac{S(i,D_j)p_j}{p_0}$$

These normalized data directly reflect the sequence dependence of the quantity $S(i, D_j)$. After this division, the remaining preparation/analysis dependent factor $\frac{p_j}{p_0}$ may be eliminated when additional information is available for the dose dependence of a specific sequence $i_0$ or of a set of sequences. In the presented case of UV-irradiation damage one knows from literature that strong damage occurs for sequences containing di-pyrimidine steps (especially TT) while all-purine sequences should show weak dose dependences. An inspection of the data shows that poly-G sequences are among the sequences with the slowest decay (for additional information see chapter SI-15). When we assume for a certain sequence $i_0$ (e. g. for poly-G in the central octamer) a defined survival probability $S(i_0, D_j) = F_{i_0}(D_j)$ we can eliminate $\frac{p_j}{p_0}$ in SI-e 3 and obtain:

(SI-e 4) $$\frac{H^{(Norm-i)}(i,j)}{H^{(Norm-i)}(i_0,j)} = \frac{S(i,D_j)p_j}{p_0}\frac{p_0}{S(i_0,D_j)p_j} = \frac{S(i,D_j)}{F_{i_0}(D_j)}$$

(SI-e 5) $$S(i, D_j) = \frac{H^{(Norm-i)}(i,j)}{H^{(Norm-i)}(i_0,j)} F_{i_0}(D_j)$$

Equation SI-e 5 shows one possibility to perform the data correction for the preparation/analysis variations. It requires knowledge on the dose dependence of a specific sequence.

Until now we treated the absolute and relative frequencies of the complete 16-mers. For a better understanding of sequence selective radiation damage, we want to consider shorter oligomers where all possible sequences are present in a chain with random contacts. In addition, we want to improve the quality of the data by averaging over the remaining parts. Therefor we split our investigated 16-mer sequence $i$ into 3 parts. The left and the right regions (sequence are $i_L$ and $i_R$) and the central region $i_C$.

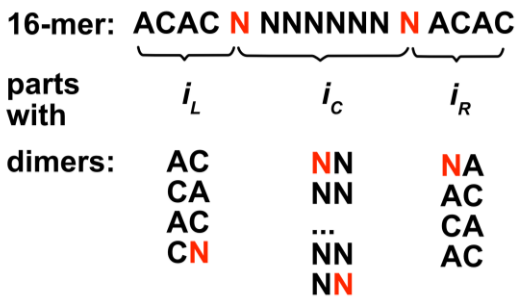

The central part $i_C$ which will be considered in the analysis shall only contain up to 6 random bases N, which are not adjacent to the nucleotides of the ACAC tails. Thus, the ACAC tails should have negligible influence on the central part.

For the elimination of parts of sequences (e. g. left or right part), which should not be considered in the analysis, we use a factorization of the survival probability of the 16-mer sequence.

(SI-e 6) $$S(i, D_j) = S_L(i_L, D_j) \cdot S_C(i_C, D_j) \cdot S_R(i_R, D_j)$$

Such a factorization is correct in situations where the survival probabilities of the different part are independent. Within the molecular model of damage formation with monomeric and dimeric damage (see SI-9) factorization is also possible. We apply (SI-e6) also for the reference sequence $i_0$, which is chosen in a way that $i$ and $i_0$ differ only in the central parts $i_C$ and $i_{C,0}$, but have the same sequences in the tails, $i_L = i_{L,0}$ and $i_R = i_{R,0}$ . When these relations are used in equation SI-e 4 the survival probabilities of the tail parts, which do not depend on the sequence of the central part cancel and a relation for the survival probability of the central part results, where $F_{i_{C,0}}(D_j) = S_C(i_{C,0}, D_j)$ refers to the survival probability of the central part of the reference sequence.

(SI-e 7) $$\frac{H^{(Norm-i)}(i,j)}{H^{(Norm-i)}(i_0,j)} = \frac{S(i,D_j)p_j}{p_0}\frac{p_0}{S(i_0,D_j)p_j} =$$
$$\left(\frac{S_L(i_L,D_j)\cdot S_C(i_C,D_j)\cdot S_R(i_R,D_j)p_j}{p_0}\right)\left(\frac{p_0}{S_L(i_L,D_j)\cdot S_C(i_{C,0},D_j)\cdot S_R(i_R,D_j)p_j}\right)$$

and therefore

(SI-e 8) $$S_C(i_C, D_j) = \frac{H^{(Norm-i)}(i,j)}{H^{(Norm-i)}(i_0,j)} F_{i_{C,0}}(D_j)$$

This equation allows to obtain the survival probability of a sequence within the central hexamer directly from the experimental data with only the assumption on the survival probability of the central part of the reference sequence. One remark should be added regarding the definition of the central part within the molecular model. Assuming that UV-damage involves only monomer and dimer lesions (see chapter SI-9) the central sequence remains intact only if all possible dimers involving bases from the central sequence remain undamaged. Therefore, the survival probability does not only refer to dimers that only involve bases from the central part, but also to dimers that may be formed with the random base directly adjacent to the central sequence (see above).

In practical applications the recorded frequencies $H^{(F)}(i,j)$ are rather small, which leads to considerable uncertainties in the values $H^{(Norm-i)}(i,j)$. To improve the statistics, equation SI-e 8 can be averaged over sequences of the left and right tail. For this reason, we sum just $H^{(F)}(i,j)$ over the tails or over all possible positions of the sequence $i_C$ (see next chapter) before computing $H^{(Norm-i)}(i,j)$. This procedure strongly reduces the uncertainties but leads to an increased influence of tail sequences with large frequencies. However, since the tails are chosen to yield survival probabilities independent of the central sequence, the procedure should not lead to systematic changes of $S_C(i_C, D_j)$.

The procedure described in this chapter directly provides the survival probability as a function of the applied dose for a sequence embedded in random bases. When the damage quantum yield $\Phi_{XY}$ for an isolated dimer XY is required, the fit of the survival probabilities $S_C(i_C, D_j)$ of the different sequences can be used to determine all required values $\Phi_{XY}$ (see chapter SI-9, equation SI- e 47).

In the next chapters we present an alternative method to determine the survival probabilities for isolated oligomers. One will find that both methods lead to the same quantum yields for dimeric damage.

#### SI-3 Improving statistics by averaging over tail sequences and positions

As already mentioned in the last chapter, the statistical error that arises in the case of small absolute frequencies can be reduced by averaging. In this chapter we consider two possible approaches. If we look at a certain center sequence of interest $i_c$ of length *k* within our total strand at position $l$, we can on the one hand average over all sequences that have this sequence $i_c$ at position $l$. On the other hand, it is worth considering extending this by averaging over different positions $l$ if it can be assumed that the absolute position of the sequence $i_c$ within our overall strand is irrelevant. We start with averaging over all possible sequences $i_L$ and $i_R$ in the tails but keep constant the central sequence $i_c$ at a position $l$ . The values labelled with an asterisk indicate this frame-averaging procedure.

(SI-e 9)

$$H^{(F,*)}(i_c, j, l) := \sum_{i \; with \; i_c \; at \; pos. \; l} H^{(F)}(i, j) = q_{i_c} p_j H^{(1,*)}(i_c, j, l)$$

Where $j$ denotes the sample id. $H^{(F)}$ was already defined in a previous chapter and indicates the experimentally found absolute frequencies for sequence $i$ in sample $j$. Analogous to the correction for the sequence and sample dependencies already considered above, the frame-averaged absolute frequency $H^{(F,*)}$ can be corrected by a factor $q_{i_c}$ and $p_j$ for the respective systematic errors, thus obtaining the corrected frame-averaged absolute frequency $H^{(1,*)}$. $H^{(0,*)}$ is the corresponding frequency for the unexposed reference samples $D = 0$. It should also be mentioned that the factor $q_{i_c}$ is not trivially related to the factor $q_i$ discussed in the last chapter for the correction of sequence dependencies within a sample. The justification for factoring out this factor results from the experimental observation that within a sample $j$ the relative frequency of frame-averaged sequences is reproducibly in a fixed relationship to each other.

##### Spatial averaging

To further improve the statistics an averaging over the different positions $l$ can be performed. This averaging is justified if the respective position of $i_c$ in the sequence $i$ is irrelevant. This is proven experimentally in as shown in Figure SI-f 7. The only restriction is to not include the positions directly adjacent to the fixed tail sequences ACAC at both ends. Quantities for which this averaging was performed are indicated by a bar on top.

(SI-e 10)

$$\bar{H}^{(1,*)}(i_c, j) := \sum_{\mathrm{l}} H^{(1,*)}(i_c, j, l)$$

$\bar{H}^{(0,*)}(i_c, l)$ represents the special case for the unexposed samples, $D = 0$.
Thus, the probability $\bar{S}^*(i_c, D_j)$ can now be determined, which represents the averaged survival of DNA strands of length 16 with fixed outer sequences ACAC at both ends and with the sequence $i_c$ in the middle range upon exposure to dose $D_j$:

(SI-e 11)

$$\bar{S}^{*}\left(i_c, D_j\right)=\frac{\bar{H}^{(1,*)}(i_c,j)}{\bar{H}^{(0,*)}(i_c,j)}=\frac{\sum_l H^{(1,*)}(i_c,j,l)}{\sum_l H^{(0,*)}(i_c,l)}=\frac{p_0 q_{i_c}\sum_l \sum_{\{i\ with\ i_c\ at\ l\}} H^{(F)}(i,j)}{p_j q_{i_c}\sum_l \sum_{\{i\ with\ i_c\ at\ l\}} H^{(F)}(i,0)}$$

$\bar{S}^{*}$ is still too specifically defined and is not yet suitable for the general determination of survival or damage probabilities for any sequences. In SI-e 11 $q_{i_c}$ cancels but the dependence on the respective sample $\frac{p_0}{p_j}$ has not yet been corrected. This will be done in the following section.

**Calculation of sub-mer damage**

In order to obtain the survival probability of a certain isolated sequence $i_c$ and not only the rather special probability of survival of DNA strands of length 16 with the edge sequences ACAC, which contain $i_c$, we can use the factorization approach again.

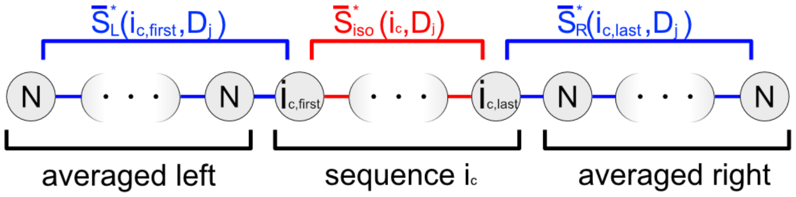

**Figure SI-f 4**. Factorization of survival probabilities according to equation SI-e 12.

We divide $\bar{S}^{*}$ into three sub-probabilities (see Figure SI-f 4). The key quantity is the survival probability $\bar{S}^{*}_{iso}\left(i_c, D_j\right)$ of the isolated sequence $i_c$ under the dose $D_j$ in the arbitrary central part of the oligonucleotides. The survival probability of the first and last arbitrary base N is biased due to the vicinity of the frame sequences ACAC at the 5’ and 3’ end of the sequence. Therefore, the survival probabilities of the frame sub-sequences ACAC and one adjacent N (arbitrary base) left and right of the sequence $i_c$ are combined to $\bar{S}^{*}_{L}\left(i_{c,first}, D_j\right)$ and $\bar{S}^{*}_{R}\left(i_{c,last}, D_j\right)$ , respectively. Analogous to the previous chapter the probabilities are assumed to be independent. This results in:

(SI-e 12)

$$\bar{S}^{*}\left(i_c, D_j\right)=\bar{S}^{*}_{L}\left(i_{c,first}, D_j\right)\cdot\bar{S}^{*}_{iso}\left(i_c, D_j\right)\cdot\bar{S}^{*}_{R}\left(i_{c,last}, D_j\right),$$

Note that the quantities $\bar{S}^{*}_{L}$, $\bar{S}^{*}_{R}$ and $\bar{S}^{*}_{iso}$ are not themselves averages over the frame sequences and position, but the asterisk and the bar here indicate that they were derived from a corresponding average value to differentiate them from the corresponding quantities in chapter SI-2. Since we are entering the range of several hundred photons/base exposure dose in our experiments, this approximation only allows the determination of the initial damage rates, as shown later by corresponding fits. Repair rates, which can only be determined via the further course and the equilibrium survival probabilities for $D_j \rightarrow \infty$, can therefore not be determined with this method.

To obtain the isolated survival probability of $i_c$, it is necessary to determine $\bar{S}_L^*$ and $\bar{S}_R^*$ beforehand. This can be achieved by treating the special case where $i_c$ corresponds to only one single base $i_c = i_c^1$. Thus $i_{c,first}$ and $i_{c,last}$ are identical and equal to $i_c^1$. The left sub strand is assumed to be $k$ bases longer than the right sub strand. The position of $i_c^1$ does not influence $\bar{S}^*$ as already discussed, but it is included in $\bar{S}_R^*$ and $\bar{S}_L^*$ with respect to $k$. One obtains from equation SI-e 12:

(SI-e 13) $$\bar{S}^*\left(i_c^1, D_j\right) = \bar{S}_L^*\left(i_c^1, D_j\right) \cdot \bar{S}_{iso}^*\left(i_c^1, D_j\right) \cdot \bar{S}_R^*\left(i_c^1, D_j\right),$$

which can be reduced to

(SI-e 14) $$\bar{S}^*\left(i_c^1, D_j\right) = \bar{S}_{L,R}^*\left(i_c^1, D_j\right)^2 \cdot \Delta s^{-k} \cdot \bar{S}_{iso}^*\left(i_c^1, D_j\right)$$

where the averaged survival probability $\bar{S}_{L,R}^*$ is defined by:

(SI-e 15) $${\bar{S}_{L,R}^*}^2\left(i_c^1, D_j\right) \coloneqq (\bar{S}_L^* - \epsilon) \cdot (\bar{S}_R^* + \epsilon) \ = \bar{S}_L^* \bar{S}_R^* + \epsilon^2.$$

where $\epsilon$ is half the difference between $\bar{S}_L^*$ and $\bar{S}_R^*$ : $2\epsilon \coloneqq |\bar{S}_L^* - \bar{S}_R^*|$ . To obtain equation SI-e 14, we assume that the survival probability of the sequence-averaged edge parts decreases exponentially with their length. This is plausible in so far as with every nucleotide added there is an additional possibility of damage. The validity of the assumption is also shown experimentally in Chapter SI-4 from the available sequence data. If the decrease in the probability of survival from an n to an (n+1)-mer is now designated Δs, the length of the edge parts, which initially differs by k as assumed, can be brought to the identical length by multiplication with $\Delta s^{-k}$. This way one gets the survival probability for an edge strand of identical length, which starts or ends with $i_c^1$.

For sufficiently small $\epsilon$, therefore, SI-e 13 follows directly from SI-e 14. $\epsilon$ is also dependent on the sequence or base $i_c^1$, but can be estimated by considering an upper limit value, by comparing the CPD quantum yields for the dimers TC and CT as shown in[2]. Here, a minimal base sequence “C” has a thymidine-nucleotide either at the 3’ or 5’ end and was irradiated with UVB light, showing a difference in quantum yield of approximately $0.4 \cdot 10^{-2}$ per photon and base. This corresponds to a maximum error $\epsilon_{max}^2 \sim 2 \cdot 10^{-5}$, which accordingly can be neglected in further considerations. In this work, the framing sequence has a length between $4$ and $7$, which reduces the numerical value of $\epsilon$ even more.
In equation SI-e 14 we can set the survival probability of the isolated monomer $\bar{S}_{iso}^*\left(i_c^1, D_j\right) = 1$ without loss of generality, because we assume that only dimer damage leads to the termination of the polymerase and thus to a detectable signal (see chapter SI-1 and SI-2). Thus, we obtain from equation SI-e 12 and SI-e 14:

(SI-e 16) $$\bar{S}_{iso}^*\left(i_c, D_j\right) = \frac{\bar{S}^*\left(i_c, D_j\right) \cdot \Delta s^{k-1}}{\left(\bar{S}^*\left(i_{c,first}, D_j\right) \cdot \bar{S}^*\left(i_{c,last}, D_j\right)\right)^{\frac{1}{2}}}.$$

The quantities $\bar{S}^*_{iso}$ on the right side of the equation can all be determined via equation SI-e 11, whereby the sample bias $\frac{p_0}{p_j}$ cancels out and only Δs needs to be determined:

$$\text{(SI-e 17)} \qquad \bar{S}^*_{iso}\left(i_c, D_j\right) = \Delta s^{k-1} \cdot \frac{\left(\frac{p_0 \sum_l \sum_{\{i\ with\ i_c\ at\ l\}} H^{(F)}(i,j)}{p_j \sum_l \sum_{\{i\ with\ i_c\ at\ l\}} H^{(F)}(i,0)}\right)}{\sqrt{\left(\frac{p_0 \sum_l \sum_{\{i\ with\ i_{c,first}\ at\ l\}} H^{(F)}(i,j)}{p_j \sum_l \sum_{\{i\ with\ i_{c,first}\ at\ l\}} H^{(F)}(i,0)}\right) \cdot \left(\frac{p_0 \sum_l \sum_{\{i\ with\ i_{c,last}\ at\ l\}} H^{(F)}(i,j)}{p_j \sum_l \sum_{\{i\ with\ i_{c,last}\ at\ l\}} H^{(F)}(i,0)}\right)}}$$

The sequence independent value of $\Delta s$ can be found by consideration of a presumably undamaged sequence $i_s$ for which the survival probability is, by definition $\bar{S}^*_{iso}\left(i_s, D_j\right) = 1$. In our analysis, we select the least damaged dimer sequence found by downstream processing of the sequencing data which reveals the oligomers with the lowest drop in sequence frequency upon irradiation. For this sequence $i_s$ of length $k_s$, we get:

$$\text{(SI-e 18)} \qquad \Delta s = \left( \frac{\left(\frac{\sum_l \sum_{\{i\ with\ i_{s,first}\ at\ l\}} H^{(F)}(i,j)}{\sum_l \sum_{\{i\ with\ i_{s,first}\ at\ l\}} H^{(F)}(i,0)}\right) \cdot \left(\frac{\sum_l \sum_{\{i\ with\ i_{s,last}\ at\ l\}} H^{(F)}(i,j)}{\sum_l \sum_{\{i\ with\ i_{s,last}\ at\ l\}} H^{(F)}(i,0)}\right)}{\left(\frac{\sum_l \sum_{\{i\ with\ i_s\ at\ l\}} H^{(F)}(i,j)}{\sum_l \sum_{\{i\ with\ i_s\ at\ l\}} H^{(F)}(i,0)}\right)^2} \right)^{\frac{1}{2\cdot(k_s-1)}}$$

Together with SI-e 17, SI-e 18 determines the normalized isolated survival probability for a sequence $i_c$. This isolated survival probability can be calculated with the measured sequence frequencies from next generation sequencing as it only includes values of $H^{(F)}$, see Figure SI-f 5. It should be mentioned here that the sequence $i_c$ used in SI-e 17 should not contain the edge sequences ACAC and the respective adjacent N (arbitrary base), since otherwise a strong sequence dependence would be induced, which would falsify the calculated damage rates.

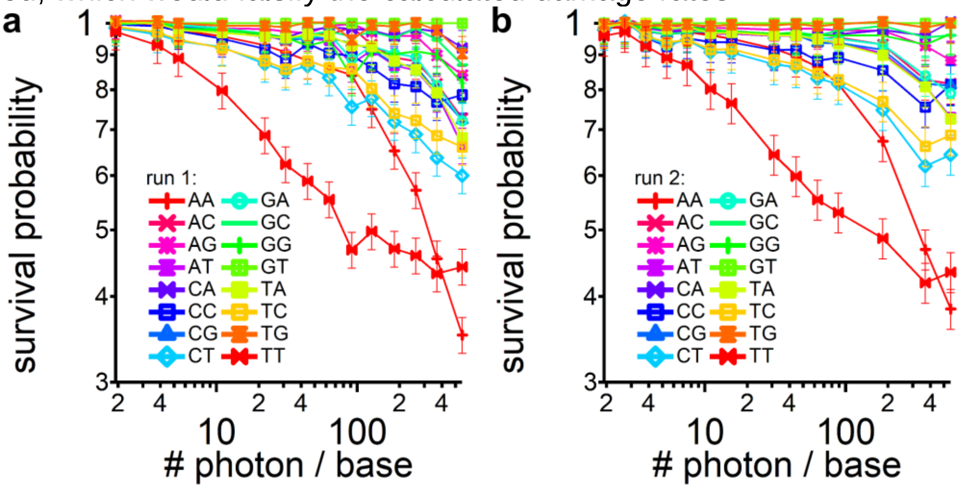

**Figure SI-f 5**: The result of equation SI-e 17 for k=2 (dimers) with the normalizing sequence $i_s = GT$ and averaged over positions 1 - 5 is plotted against the irradiation dose (# photon / base) for two experimental runs (**a** and **b**).

#### SI-4 Exponential length dependence of the survival probability

To experimentally validate Equation SI-e 16/17 and SI-e 14 and therefore the assumption of an exponential dependence of survival probability on the sequence length, we average over all $M$ possible dimer sequences $i_c$ that must not contain parts of the edge sequences ACAC and the respective adjacent N (arbitrary base):

(SI-e 19) $$\frac{1}{M}\sum_{i_c}\bar{S}^{*}_{iso}\left(i_c,D_j\right)=\frac{1}{M}\sum_{i_c}\frac{\bar{S}^{*}(i_c,D_j)\cdot\Delta s}{\sqrt{\bar{S}^{*}(i_{c,first},D_j)\bar{S}^{*}(i_{c,last},D_j)}}$$

(SI-e 19a) $$\sum_{i_c}\bar{S}^{*}_{iso}\left(i_c,D_j\right)=\Delta s\cdot\sum_{i_c}\frac{\bar{S}^{*}(i_c,D_j)}{\sqrt{\bar{S}^{*}(i_{c,first},D_j)\bar{S}^{*}(i_{c,last},D_j)}}$$

The left side averages over the survival probabilities of all possible dimer sequences and hence can be described as the mean dimer survival probability. If our assumption is correct and the change in survival probability due to an additional dimer is described on average by an additional factor $\Delta s$, and thus the overall survival probability of the strand decreases exponentially with length, $\Delta s$ would have to correspond to the average survival probability of a dimer $\Delta s=\sum_{i_c}\bar{S}^{*}_{iso}\left(i_c,D_j\right)$. The experimental values of the sum on the right-hand side of Equation SI-e 19a should therefore be close to unity:

(SI-e 20) $$\sum_{i_c}\frac{\bar{S}^{*}(i_c,D_j)}{\sqrt{\bar{S}^{*}(i_{c,first},D_j)\bar{S}^{*}(i_{c,last},D_j)}}\overset{!}{\approx}1$$

Figure SI-f 6 shows the left-hand side of SI-e 20 of all samples $j$ is indeed close to unity within 3% which validates the assumption of an exponential length dependence of survival probability reasonably well.

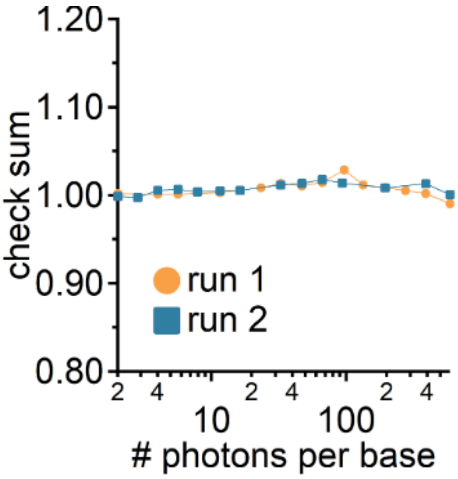

**Figure SI-f 6**: The averaged sum shown in the left-hand side of Equation SI-e 20 is plotted for all used samples for $i_c = dimers$. It is 1 within a 3% range, showing the experimental validity of approximations done for SI-e 14.

#### SI-5 Validation for sequence-averaging over all positions

Especially long exposure times and therefore high dosages decrease the number of intact strands and sequence frequencies which leads to low signal to noise ratios.

We improved the situation by averaging over all positions within the central, randomized hexamer ($N_1 - N_6, N \in \{A, G, C, T\}$). The randomized bases $N_0$ and $N_7$ at the edge positions were not considered, because of the sequence-bias introduced by the adjacent bases C and A of the ACAC sequence on the left and right end of the strand, respectively. To exclude a position-related effect on the survival probability $\bar{S}^*_{iso}$, we evaluated SI-e 17 for all possible positions within the central hexamer. The results were compared to the position average for sequencing runs 1 (Figure SI-f 7a) and 2 (Figure SI-f 7b) which showed an insignificant dependence on the position of 1.1% and 0.2% deviation from the average value, respectively. This is small compared to the drop of survival probabilities induced by irradiation of up to 60%.

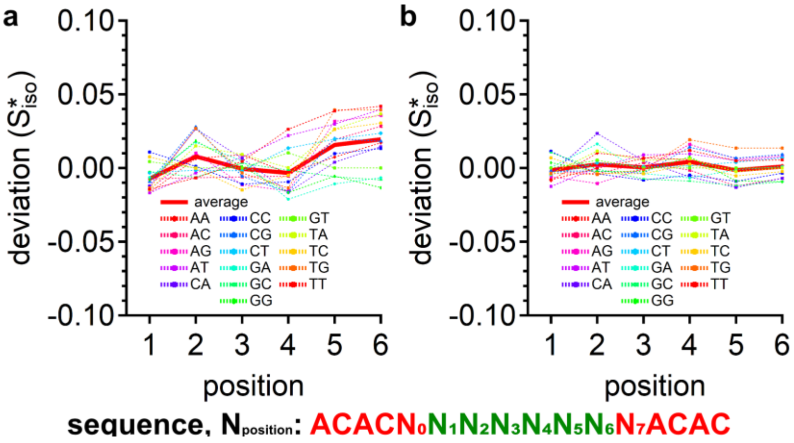

**Figure SI-f 7**: The deviation of isolated survival probabilities of dimers ($k = 2$) at positions $l = 1 \ldots 6$ is evaluated and compared against the position average ($l = 1\ to\ 5$) value for all sequences using Equation SI-e 17 for run 1 (**a**) and run 2 (**b**). It shows only a small difference of 1.1% and 0.2% for run 1 and 2, respectively. Since the deviation is well below the statistical error of ~6%, averaging of the survival probabilities over all positions is performed to increase the signal to noise ratio without masking position-dependent effects on the damage rates.

#### SI-6 Sequence symmetry of survival probabilities

The assumption of sequence symmetric damage can be made if the relative survival probabilities $\bar{S}^*_{iso}(i, D_j)$ of a sequence $i$ is equivalent to the survival probability of its reversal sequence $\bar{i}$, e.g. $i = ACGGTC, \bar{i} = CTGGCA$. Analysis of this correlation for the obtained sequencing data sets shows a clear dependency $\bar{S}^*_{iso}(i) \sim \bar{S}^*_{iso}(\bar{i})$, as the mean square displacement over $M$ different sequences $i$, $\frac{1}{M}\sum_i \left(\bar{S}^*_{iso}(i) - \bar{S}^*_{iso}(\bar{i})\right)^2$ is below 0.2% for hexamers (see Figure SI-f 8d, h). Examples are shown for data set 1 in Figure SI-f 8a-c and data set 2 in Figure SI-f 8e-g, with the black line indicating the relative frequencies of the sequences i and the blue dots denoting the relative frequency of its reversed sequence $\bar{i}$, plotted at position i. For better visibility, the sequences represented by the black line were arranged in ascending order according to their frequency. If both frequencies would be identical, the black and blue curve would coincide and the mean square displacement between both values would vanish. Despite of experimental error, both data sets still show a strong correlation as shown in Figure SI-f 8.

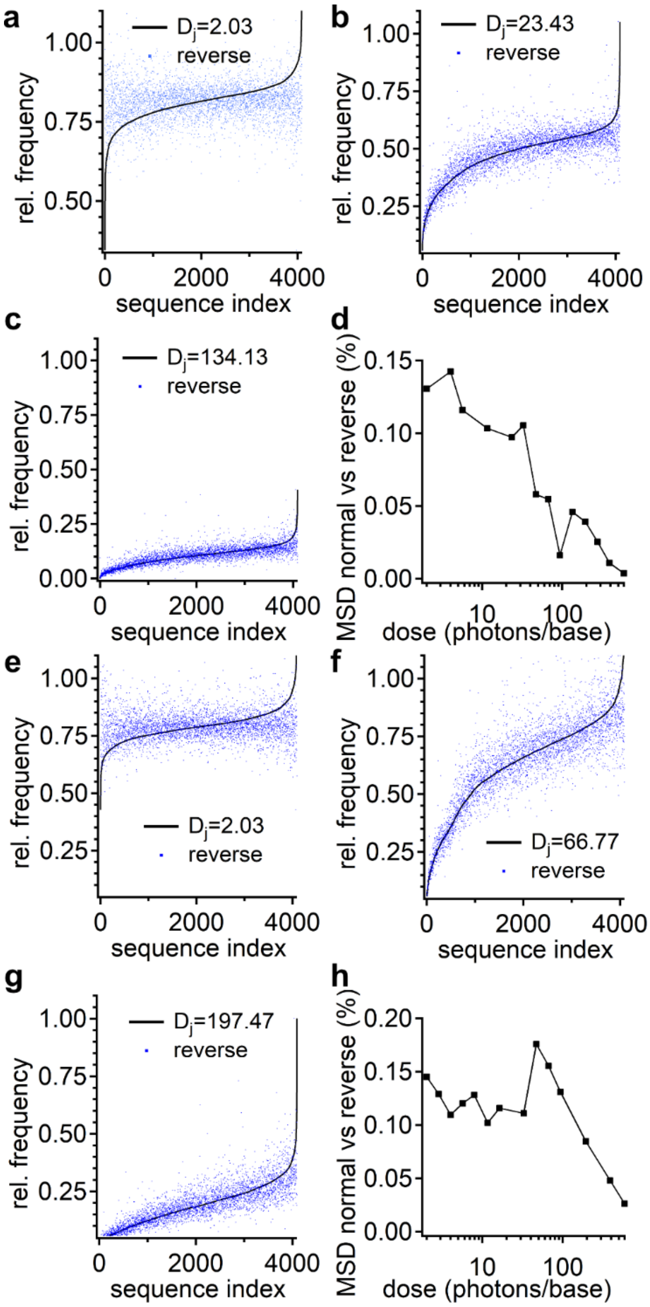

**Figure SI-f 8:** Comparison of survival probabilities of sequences and their reverse counterparts: Data set 1 with doses 2.03 photons/base (**a**), 23.43 photon/base (**b**), and 134.13 photon/base (**c**). (**d**) MSD of relative survival probabilities for arbitrary sequences $i$ against its reversal sequences $\bar{i}$ for data set 1. Corresponding survival probabilities for data set 2 with doses 2.03 photon/base (**e**), 66.77 photon/base (**f**), and 197.47 photon/base (**g**). (**h**) MSD of relative survival probabilities for arbitrary sequences $i$ against its reversal sequences $\bar{i}$ for data set 2.

#### SI-7 Absorbed Dose

**The average number of absorbed photons per base**, we denominate it the dose D, can be determined as the ratio of the numbers of absorbed photons and the number of bases $N_{base}$ in the sample. We use this number to characterize the excitation of the sample. In the irradiation experiment the power of the UV-radiation is measured by calibrated power meters for the radiation impinging on the sample and transmitted through the sample. Together with the known coefficients of reflection at the sample windows these values allow to calculate the power $W_{abs}$ absorbed in the sample. The numbers of absorbed photons $N_{abs}$ is obtained by integration over the irradiation time and division by the photon energy $h\nu$.

(SI-e 21) $$N_{abs} = \frac{\int W_{abs} dt}{h\nu}$$

The number of bases $N_{base}$ in the sample of volume V is estimated via the Lambert-Beer law (see equation (SI-e 23), which relates the absorbance with the concentration [C], the extinction coefficient $\varepsilon$ and the sample thickness d. For the sample containing oligomers with a defined sequence the extinction coefficient of the oligomer is determined according to the nearest neighbor model (see e.g. Ref. 3, parameters given for a wavelength of 260 nm). For the irradiated 16-mer ACACNNNNNNNNACAC the randomized bases N are considered by using the parameters $\varepsilon_N, \varepsilon_{C,N}, \varepsilon_{N,A}, \varepsilon_{N,N}$ obtained by appropriately averaging the coefficients $\varepsilon_i, \varepsilon_{i,j}$ given in Table 2 of Ref. 3.

$$\varepsilon_N = \frac{\varepsilon_A+\varepsilon_C+\varepsilon_G+\varepsilon_T}{4};\ \varepsilon_{C,N} = \frac{\varepsilon_{C,A}+\varepsilon_{C,C}+\varepsilon_{C,G}+\varepsilon_{C,T}}{4};\ \varepsilon_{N,A} = \frac{\varepsilon_{A,A}+\varepsilon_{C,A}+\varepsilon_{G,A}+\varepsilon_{T,A}}{4};$$

$$\varepsilon_{N,N} = \frac{1}{16} \cdot \sum_{i,j \in \{A,C,G,T\}} \varepsilon_{i,j}$$

The average extinction coefficient $\varepsilon_{16av}$ of a base in the 16-mer is calculated using equation 9 of Ref. 3:

(SI-e 22) $$\varepsilon_{16av} = \frac{\sum_{i=1}^{15} \varepsilon_{i,i+1} - \sum_{i=2}^{15} \varepsilon_i}{16} = \frac{1}{16} \cdot \big(\epsilon_{A,C} + \epsilon_{C,A} + \epsilon_{A,C} + \epsilon_{C,N} + 7\epsilon_{N,N} + \epsilon_{N,A} + \epsilon_{A,C} + \epsilon_{C,A} + \epsilon_{A,C} - 3\epsilon_A - 3\epsilon_C - 8\epsilon_N\big)$$

**Table SI-t1:** Values of parameters used in the nearest neighbor model for the calculation of the average extinction coefficients of single stranded oligomers containing randomized bases N. The parameters not presented in the table are given in Ref. 3, Table 2. All parameters are determined for a wavelength of 260 nm.

| | $\varepsilon_{260nm}$[1/(M cm] |
|---|---|
| $\varepsilon_N$ | 10750 |
| $\varepsilon_{C,N}$ | 17250 |
| $\varepsilon_{N,A}$ | 24300 |
| $\varepsilon_{N,N}$ | 20325 |

For the determination of base concentration [$C_{base}$] or number of bases $N_{base}$ we calculate the average extinction coefficient $\varepsilon_{16av}$ from equation SI-e 22 to be $\varepsilon_{16av}$ = 9789 1/(M cm), and use a measurement of the absorbance $A_{260}$ at λ = 260 nm, via a spectrophotometer and a sample cell with a defined path-length d.

(SI-e 23) $A_{260} = \varepsilon_{16av} \cdot [C_{base}] \cdot d$ Lambert-Beer Law

(SI-e 24) $[C_{base}] = \frac{A_{260}}{\varepsilon_{16av} \cdot d}$

(SI-e 25) $N_{base} = [C_{base}] \cdot V \cdot N_A = \frac{A_{260}}{\varepsilon_{16av} \cdot d} \cdot V \cdot N_A$

Here $N_A$ denotes Avogadro's number. The dose D, i.e. average number of absorbed photons per base, can now be estimated.

(SI-e 26) $D = \frac{N_{abs}}{N_{Base}}$

**The average number of photons absorbed per base of an oligomer**, $D_{olig}$ is the important quantity when one is interested in the dose-resistance of an oligomer with a defined sequence. While the average number of absorbed photons per base of the irradiated 16-mer D was determined via equations (SI-e 24 - 26) from the extinction coefficient and the absorption taken at a wavelength of 260 nm, $D_{olig}$ depends on the ratio of extinction coefficients taken at the irradiation wavelength 266nm.

(SI-e 27) $D_{Olig} = D \cdot \frac{\varepsilon_{Olig}(266)}{\varepsilon_{16av}(266)} \cdot \frac{1}{N_{Olig}}$

Here $N_{Olig}$ is the number of bases in the oligomer. The averaged extinction coefficient of the 16-mer at a wavelength of 266 nm is obtained from the measured absorption spectrum of the 16-mer and the extinction coefficient determined at 260 nm.

(SI-e 28) $\varepsilon_{16av}(266) = \varepsilon_{16av}(260) \cdot \frac{A_{16av}(266)}{A_{16av}(260)} = 8984$ /(M cm)

The extinction coefficient of the oligomer $\varepsilon_{Olig}(266)$ can be calculated via the nearest neighbor model with the parameters determined at a wavelength of 266 nm. These parameters may be obtained from the parameters given at 260 nm combined with the absorption properties of all possible DNA dimers. An evaluation of the dose $D_{Olig}$ for a series of tetramers indicated that the dose $D_{Olig}$ of adenine-rich oligomers is up to ca. 15% above the average dose D, while adenine-poor oligomers have dose values of ca. 5 % below the average dose D. These numbers indicate that the average dose D yields a good measure for the qualitative determination of dose dependences of different oligomers.

#### SI-8 Molecular Model for Dose-Dependence

Here, we consider a sample containing many oligomers of different types which will be exposed to UV-radiation. The oligomers in the sample have the same length L. In addition, we assume that all oligomers have the same extinction coefficient (when the difference in extinction coefficients should be considered explicitly, the individual dose values of the different oligomers from Section SI-7, equations SI-e 27, 28, have to be used below instead of the average value ΔD·L). We assume that the absorption of the sample should not change during our irradiation experiment. This is justified for the irradiation doses used in the experiment. The damage of a single dimer in the 16-mer is sufficient to destroy the integrity of the 16-mer but changes only the absorption of a small fraction of the absorbing bases. Indeed, even at the highest irradiation doses used in this paper, the absorption of the sample is changed by less than 20%.

In the following we consider one specific type of oligomers with sequence $i$ in the sample containing many other oligomers. The frequency of the specific sequence of type $i$ of intact oligomers in the sample after irradiation with a dose $D$ is denominated $Q_i(D)$. Prior to irradiation we have $Q_i(0) = H^{(0)}(i)$ (see Equation (SI-e 1)). Here, we use the denomination $Q_i(D)$ to differentiate from the various measured and normalized quantities H used in section SI-2**.**

When a small dose $\Delta D$, $\Delta D << 1\, absorbed\, photon/base$, is applied to the sample, a fraction of the oligomers will be excited. $\Delta Q_i^*$, the number of oligomers of type $i$ excited by the dose $\Delta D$, is proportional to the number $Q_i$ of intact oligomers present at the time of irradiation times the number of photons absorbed by the oligomer ($L \cdot \Delta D$).

(SI-e 29) $$\Delta Q_i^* = L \cdot \Delta D \cdot Q_i$$

When a certain small percentage of these excited oligomers becomes damaged with the damaging quantum yield $\Phi_i$, the number of intact oligomers will decrease by an amount $\Delta Q_i$..

(SI-e 30) $$\Delta Q_i = -\Phi_i \cdot \Delta Q_i^* = -\Phi_i \cdot L \cdot \Delta D \cdot Q_i$$

(SI-e 31) $$\Delta Q_i = -\mu_i \cdot \Delta D \cdot Q_i$$

Here, $\mu_i$ is the damage constant, describing the response of the specific oligomer to the applied UV-radiation. With infinitesimally small dose changes $\Delta D \rightarrow 0$ Equation (SI-e 31) can be converted into a differential Equation with the corresponding solution:

(SI-e 32) $$\frac{dQ_i}{dD} = -\mu_i \cdot Q_i$$

(SI-e 33) $$Q_i(D) = Q_i(0) \cdot \exp(-\mu_i \cdot D)$$

Thus the survival probability becomes $S(i,D) = Q_i(D) \,/\, Q_i(0) = \exp(-\mu_i \cdot D)$ i.e. a single exponential decay describes the dose dependence of the special oligomer with sequence $i$. The decay constant $\mu_i$ represents the damaging coefficient for the special oligomer. For an applied dose $D_\mu = \frac{1}{\mu_i}$, the survival dose, the survival probability is reduced to $\frac{1}{e}$. $D_\mu$ gives a qualitative measure for the dose resistance of certain oligomer $i$.

Quite often the recorded data show dependences, which deviate from a single exponential form (see Figure 3, main text). This may be due to a more complex reaction scheme. Equation (SI-e 32) represents the reaction out of a defined state, describing the intact oligomer. The UV radiation damage converts the intact oligomer into the damaged one. There is no further reaction involved. Especially there is no backreaction (see reaction scheme in SI-f 9a). In pyrimidine dimers however, several types of photolesions are possible. An example for a reaction scheme describing these processes is given in SI-f 9b, the corresponding rate equation system (SI-e 34). For the CPD-lesion we know that appropriate UV-irradiation can reestablish the undamaged, intact oligomer (damage Type 2, frequency $Q2_i$). For other lesions, e. g. the (6-4)-lesion a direct back-conversion to the intact oligomer is does not occur (damage Type 1).

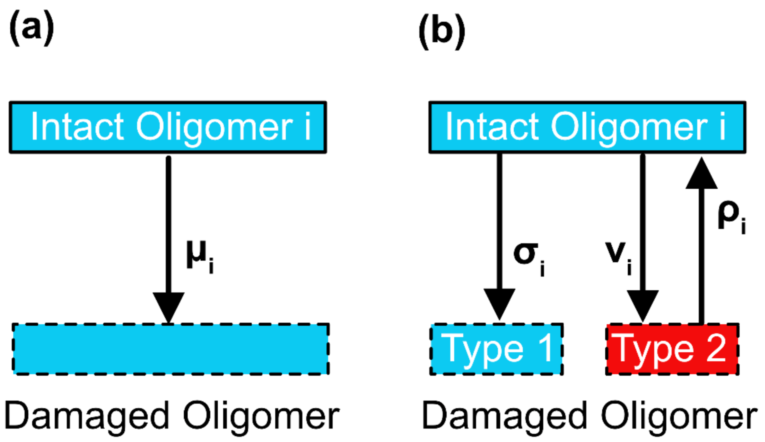

**Figure SI-f 9**: Different reaction models used in the description of the UV-induced damage. (a) Simple reaction scheme. From the intact oligomer $i$ UV-radiation induced transition to the damaged structure with the damage coefficient $\mu_i$ (there is no back-reaction). The population of the intact oligomers decays exponentially with the decay constant $\mu_i$. (b) A prototype reaction scheme with back-reactions leads to a bi-exponential dose-dependence. In this scheme there is the irreversible damage coefficient $\sigma_i$, the reversible damage coefficient $\nu_i$, and the repair coefficient $\rho_i$.

(SI-e 34) $$\frac{dQ_i}{dD} = -(\nu_i + \sigma_i)Q_i + \rho_i Q2_i,\ \frac{dQ2_i}{dD} = +\nu_i Q_i - \rho_i Q2_i$$

Since the total number of oligomers $i$ (intact and damaged of both types) is constant the solution of the rate equation system for the number of intact oligomers becomes a biexponential function of the absorbed dose $D$.

(SI-e 35) $$\frac{Q_i(D)}{Q_i(0)} = A_{i1}\exp(\gamma_{i1}D) + A_{i2}\exp(\gamma_{i2}D)$$

The decay constants depend on the damage and repair coefficients.

(SI-e 36)

$$\gamma_{i1} = \frac{-(\nu_i + \rho_i + \sigma_i) + \sqrt{(\nu_i + \rho_i + \sigma_i)^2 - 4\rho_i\sigma_i}}{2}$$

$$\gamma_{i2} = \frac{-(\nu_i + \rho_i + \sigma_i) - \sqrt{(\nu_i + \rho_i + \sigma_i)^2 - 4\rho_i\sigma_i}}{2}$$

The faster decay constant ($\gamma_i$) is visible at low doses, the slower ($\gamma_i$) at high dose levels. When the system starts initially from intact oligomers with frequency $Q_i(0)$, one obtains further relations: $A_{i1} + A_{i2} = 1$ and $A_{i2} = 1 - A_{i1}$ . A fit of the experimental data with a bi-exponential function should yield the amplitudes $A_{i1}$, $A_{i2}$ and the decay coefficients $\gamma_i$ and $\gamma_i$. Often, the initial slope of the dose dependence of the frequency of intact oligomers can be obtained without elaborate fitting. It gives information on the total damaging, i.e. on the sum of the damaging coefficients $\nu_i + \sigma_i$ :

(SI-e 37) $$\frac{dQ_i}{dD}/Q_i\Big|_{D=0} = \nu_i + \sigma_i = -(A_{i1}\gamma_{i1} + A_{i2}\gamma_{i2})$$

The other damage or repair coefficients may be obtained from the fitting results using the following relations.

(SI-e 38) $\rho_i = A_{i1}(\gamma_{i1} - \gamma_{i2}) - \gamma_{i1}$

(SI-e 39) $\sigma_i = \frac{\gamma_{i1}\gamma_{i2}}{A_{i1}(\gamma_{i1}-\gamma_{i2})-\gamma_{i1}}$

(SI-e 40) $\nu_i = -(\gamma_{i1} + \gamma_{i2} + \rho_i + \sigma_i) = -\left(\gamma_{i1} + \gamma_{i2} + A_{i1}(\gamma_{i1} - \gamma_{i2}) - \gamma_{i1} + \frac{\gamma_{i1}\gamma_{i2}}{A_{i1}(\gamma_{i1}-\gamma_{i2})-\gamma_{i1}}\right)$

#### SI-9 A Molecular Model for the Damage Coefficient of Oligomers

In the single stranded chain used in the present experiments the most prominent damage processes are the formation of di-nucleotide lesions like the cyclobutane pyrimidine dimer CPD) lesion or single nucleotide damage. CPD is formed by bridging two neighboring pyrimidines (thymine or cytosine) by a cyclobutane ring. Those damages have typical quantum efficiencies in the $10^{-2}$ to $10^{-3}$ range.
Quantitative information on the underlying molecular processes and a comparison with the quantum yields given in the literature may be obtained when combining the observations with a molecular model for damage formation. Here, we use a simple model, which is based on dimeric damage processes and nearest neighbor interactions and addresses only the initial step of damage formation (see scheme in Figure 5, insert). We assume (i) the same excitation of all bases in the strand, (ii) that the excitation of one base X in a WXY step is equally shared with the two neighboring bases W and Y leading to excited dimer states (WX)* and (XY)* with the same probability of $M_{WX*} = M_{XY*}$ = 0.5 and (iii) causing damages of the respective dimers with the quantum yields $\Phi_{WX}$ and $\Phi_{XY}$. As a simplifying assumption we use (iv) symmetric quantum efficiencies, $\Phi_{XY}$ = $\Phi_{YX}$. This model connects the experimentally observed damaging coefficient $\mu_{olig}$ with the quantum yields $\Phi_{XY}$ of the dimers of the oligomer.
The formation of the dimeric damage (*X*=*Y)* after the absorption of one photon in base $X$ should occur (assumption (ii) and (iii)) with the yield $\mu_{XY}$ .

(SI-e 41) $\mu_{XY} = M_{XY*} \cdot \Phi_{XY} = 0.5 \cdot \Phi_{XY}$.

Here, $\Phi_{XY}$ is the damage formation quantum yield from the excited state $(XY)^*$. Simultaneously the absorption in base $X$ causes the formation of damage (*W*=*X)* with the yield $\mu_{WX}$.

(SI-e 42) $\mu_{WX} = 0.5 \cdot \Phi_{WX}$.

Above, we assumed that the illumination with UV-radiation at 266 nm leads to the same excitation for all bases independent of the type (assumption (i)). When the exact absorption cross-section should be considered, correction factors must be applied to obtain exact quantum efficiencies. For the wavelength of 266 nm, used in the experiments, this correction should be less than 20% (see SI-7).
The damage coefficient of a complete oligomer $\ldots VWXYZ$.., may be calculated by summing over the damaging yields of the individual dimers considering the applied dose $D$. This is justified since the complete oligomer remains only intact, when all dimers, composing the oligomer are undamaged. In other words, each damage of a dimer in the oligomer adds up to the damage of the complete oligomer. For the

survival probabilities one may multiply the survival probabilities of all individual dimers to obtain the survival probability of the complete oligomer.

(SI-e 43) $$\mu_{olig} = \Phi_{VW} + \Phi_{WX} + \Phi_{XY} + \Phi_{YZ} + \Delta\mu_V + \Delta\mu_Z$$

Here, special effort must be laid on the ends of the oligomer (correction term $\Delta\mu_V + \Delta\mu_Z$). When the oligomer is isolated (sequence $VWXYZ$) no excitation transfer can occur to the outside. Thus, the damaging coefficient of the outmost dimers is increased by $\Delta\,\mu_V = 0.5\ \Phi_{VW}$ and $\Delta\mu_Z = 0.5 \cdot \Phi_{YZ}$. When the oligomer is embedded in randomized bases (...$NVWXYZN$..), the corresponding excitation will lead to an increase due to the damaging coefficient of the dimer composed of $V$ and the precursor nucleotide or of $Z$ and the subsequent nucleotide. Since these nucleotides are chosen in a randomized way, we obtain:

(SI-e 44) $$\Delta\mu_V = \frac{1}{4} \cdot (\Phi_{AV} + \Phi_{CV} + \Phi_{GV} + \Phi_{TV})$$

(SI-e 45) $$\Delta\mu_Z = \frac{1}{4} \cdot (\Phi_{ZA} + \Phi_{ZC} + \Phi_{ZG} + \Phi_{ZT})$$

Equations SI-e 43 to 45 yield the damage of the central sequence $VWXYZ$ induced by the photons absorbed in $NVWXYZN$. In addition, the outer parts (damage induced by photons absorbed in the outer parts) must also be considered when calculating the damage coefficient of the complete DNA-strand.

**Factorization of the survival probabilities.** The presented molecular model has the important property that it allows the factorization of the survival probabilities. This property may be illustrated with an example: When we take an oligomer with the sequence of bases TUVWXYZ, the decay constant of the oligomer becomes:

$$\mu_{olig} = 1.5\ \Phi_{TU} + \Phi_{UV} + \Phi_{VW} + \Phi_{WX} + \Phi_{XY} + 1.5\ \Phi_{YZ}$$

This relation allows a formal splitting of the oligomer at a base, e.g. at base W and the resulting parts have the following decay constants

$$\mu_{olig\_left} = 1.5\ \Phi_{TU} + \Phi_{UV} + \Phi_{VW}$$
$$\mu_{olig\_right} = \Phi_{WX} + \Phi_{XY} + 1.5\ \Phi_{YZ}$$

Here the sum of the decay constants of the two parts is equal to the decay constant of the complete oligomer. For the survival probabilities of the respective parts we find:

$$S(olig,D)=\exp(-\mu_{olig}D)= \exp(-\mu_{olig\_left}D)\ \exp(-\mu_{olig\_right}D)= S(olig\_left,D)\ S(olig\_right,D)$$

This factorization of the survival probabilities was used above in Sections SI-2 and SI-4.

The survival probability of the whole oligomer is the product of survival probabilities of its parts if and only if the survival chances of the parts are pairwise independent of each other. In the damage model presented above this independency originates from the transfer of the excitation of base W to the excited dimers (VW)* and (WX)* with equal efficiency. When the model is refined e. g. when considering processes due to charge transfer states there are deviations from the pure dimeric damage model and the survival probabilities of the different parts are no longer independent. However for weak excitation the dimeric mode may be taken as a first order approximation.

**Symmetry of the damage coefficients.** In the presented evaluation procedure, the damage coefficient of a specific oligomer is calculated as a sum over the dimeric damage coefficients $\Phi_{XY}$. Thus, the knowledge of the 16 coefficients $\Phi_{XY}$ ($X, Y \in \{A, C, G, T\}$) should be required to estimate the damage of arbitrary oligomers. The analysis of the experimental data has revealed that pairs of oligomer sequences with reversed order relative to the 5' or 3' ends, show similar dose dependences (see SI-6). Thus, the dimer damage coefficients can be assumed to be symmetric, $\Phi_{XY} = \Phi_{YX}$ (assumption (iv)). E. g. one may assume the same value for dimer damage coefficient for the CT as for the TC dimer, $\Phi_{CT} = \Phi_{TC}$. Under these conditions there are ten independent values $\Phi_{XY}$: $\Phi_{AA}, \Phi_{AC} = \Phi_{CA}, \Phi_{AG} = \Phi_{GA}, \Phi_{AT} = \Phi_{TA}, \Phi_{CC}, \Phi_{CG} = \Phi_{GC}, \Phi_{CT} = \Phi_{TC}, \Phi_{GG}, \Phi_{GT} = \Phi_{TG}, \Phi_{TT}$. An example for the tetramer CATT with randomly chosen neighbors is given here.

(SI-e 46) $$\mu_i = \mu_{CATT} = \Phi_{left} + \Phi_{CA} + \Phi_{AT} + \Phi_{TT} + \Phi_{right} = \frac{1}{4}\cdot(\Phi_{AC} + \Phi_{CC} + \Phi_{GC} + \Phi_{TC}) + \Phi_{AC} + \Phi_{AT} + \Phi_{TT} + \frac{1}{4}\cdot(\Phi_{TA} + \Phi_{TC} + \Phi_{TG} + \Phi_{TT})$$

Here the expressions within the parentheses are due to the left and right borders.

**Determination of the dimeric quantum yield $\Phi_{XY}$ from the experimental data.** It should be noted that in general the damage coefficients are linear combinations of the dimeric quantum efficiencies $\Phi_{XY}$. For the general case of oligomers of length $n$ one may write the corresponding relation in an appropriate vector form.

(SI-e 47) $$\vec{\mu} = \overleftrightarrow{A} \cdot \overrightarrow{\Phi} \ or\ \mu_i = \sum_{j=1}^{10} A_{ij} \cdot \Phi_j \ with\ i = 1..4^n, \Phi_1 = \Phi_{AA}, \Phi_2 = \Phi_{AC}, .., \Phi_{10} = \Phi_{TT}$$

The coefficients of the matrix $A$ can be directly determined from the base sequence of the respective oligomer together with the information on the terminal parts, as shown above.
In the experiment, the damage coefficients $\mu_i$ may be obtained from a fit of the experimental data, i.e. from the survival probabilities of the $4^n$ different intact oligomers recorded as a function of the illumination dose $D$. The solution of the linear system of Equations SI-e 47 with the known matrix $A$ and the experimental results $\mu_i$ can be used to determine the unknown molecular quantum efficiencies $\Phi_{XY}$ [2]. These numbers should be compared with the dimeric quantum efficiencies recorded by other techniques (see main part, Figure 5).

During the data analysis, we recognized that the dimeric quantum efficiency $\Phi_{GT}$ has the tendency to become negative, which would be physically unreasonable. Therefore, one may argue that one of the assumptions of the modeling procedure given above is not justified. A negative value of $\Phi_{GT}$ could point to repair (unreasonable for an intact dimer) or to the possibility that the dimer $GT$ would dump the excitation of an initially excited $T$ very efficiently (violation of assumption (i) to (iii)). The dumping could occur via the charge transfer state $G^+T^-$ which is known to be efficiently formed in $GT$ steps[4]. We discuss this point here in the context of a sequence ...$GTX$... . In the case where $T$ is excited initially (state ...$GT^*X$...), dumping the excitation via $G^+T^-$ prior to the formation of state ...$G(TX)^*$... would reduce the damage yield for the complete oligomer, since the lesion $...GT = X...$ cannot be formed via $G^{+}T^{-}$. Thus, the presence of a $GT$ step can reduce the damage yield via dumping the excitation of the thymine $T$. Similar processes could occur also for $AT$, $GC$ and $AC$ dimers. However, the redox potentials of the different bases indicate, that the excitation dumping should be most efficient in $GT$ and $GC$ steps. The dumping process can be incorporated in Equation (SI-e 47) by suitable modifications of the matrix $A$.

#### SI-10 Fitting of damage rates

The isolated survival probabilities shown in Figure SI-f 5**Figure SI-f 5** show a significant drop in the abundance of undamaged dimers over the course of the irradiation experiment. To quantify the effect, we fitted a three-state model to the experimental data, featuring one reversible and one irreversible state connected by a ground state SI-f 9b). Less complex models (e.g. two-state) were not able to fit the experimental data sufficiently well.

As shown before, this model implemented by a bi-exponential function of the absorbed dose $D_j$:

(SI-e 48) $$\bar{S}^{*}_{iso}(i_c, D_j) = A_1 \exp(\gamma_{i_c,1} \cdot D_j) + A_2 \cdot \exp(\gamma_{i_c,2} \cdot D_j)$$

with the coefficients $A_1, \gamma_{i_c,1}, A_2, \gamma_{i_c,2}$ to be determined from fitting using the nonlinear Levenberg-Marquardt algorithm. The transition rates are then obtained by evaluating Equations SI-e 38-40. Experimental data were smoothed before fitting by a standard LOWESS algorithm to stabilize the fitting procedure against experimental outliers by e.g. low sequencing counts[5]. Fit weights $w_{i,j}$ were estimated after error propagation of the standard deviation from the experimental raw data through all corrections up to the final isolated survival probabilities $\bar{S}^{*}_{iso}(i_c, D_j)$ using $w_{i_c,j} = \frac{1}{\Delta^2}$. The final error estimates for the transition rates $\rho_{i_c}, \sigma_{i_c}, \mu_{i_c}$ were obtained by averaging the fit results from run 1 and 2, see Figure SI-f10.

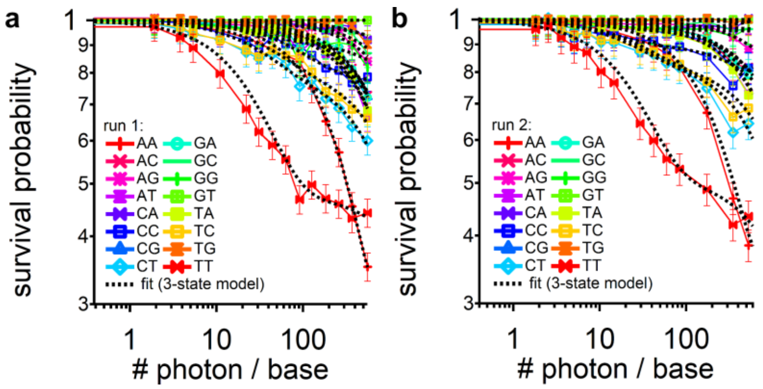

**Figure SI-f 10**. The experimentally obtained isolated survival probabilities for dimers (see) fits well a bi-exponential three-state model (**a** = run 1, **b** = run 2).

**The fitting of the dose dependences** of the dimer, tetramer and hexamer data shown in the main document (Figure 3 and 4) was performed using the routine CurveFit with the function dblexp_XOffset from Igor pro (Wavemetrics, USA). The main difficulty of the data fitting originates from the fact, that the most simple mathematical model leads to a mono-exponential dose dependence of the relative frequencies, while a more complicate model (which is supported by the experimental dose dependences of some sequences) requires a bi-exponential fit function. Thus data fitting has to decide between a mono-exponential or a bi-exponential behavior and has to determine the amplitudes and the decay constants for the different oligomer sequences.

To accomplish the data analysis we developed a multistep decision process:

1. For each sequence the complete dose dependence of the relative frequency was fitted by the double-exponential fitting tool with few constraints on the two amplitudes and the two decay constants (e.g. no rising components, sufficiently large amplitude of the fast component as compared to the data noise, clearly different decay constants). When this fit converged and supplied amplitudes and decay constants with sufficiently small errors, the data were directly used.

Otherwise:

2. Two subsets of the data were selected for small dose values (D < 35 photon/base) and large dose values (D > 60 photon/base) and each subset is fitted by a mono-exponential function. When both fits converged and yielded consistent data (no rising components, small dose values yield faster decay, sufficiently large amplitude of the fast component as compared to the data noise, clearly different decay constants) the results were combined to a bi-exponential function and the corresponding two amplitudes and two decay constants are stored.

Otherwise:

3. The complete data set was fitted by a mono-exponential function. The amplitude and the decay constant were stored.

4. Finally the results of the multistep fitting-procedure were used to calculate the initial damaging constant $\mu_i = \nu_i + \sigma_i$ using equation (SI-e 37).

When comparing the experimental data points and the modeling functions obtained by the presented procedure we find a reasonably "good" fit for nearly all sequences.

**Experimental errors in the determination of the quantum yields.**

Since the quantum yields for the dimeric damage shown in Figure 5 were obtained by the solution of the linear system of equations (SI-e 47), the statistical uncertainties of the experimental values $\mu_i$ can only be used indirectly to determine the error of the quantum yields. These errors are estimated as follows: The linear system of equations (SI-e 47) is solved repeatedly (1000 times) with $\mu_i$ values for the different sequences taken from a Gaussian distribution of the $\mu_i$ (widths according to the fitting errors of the individual $\mu_i$). The total error is obtained by combining these statistical errors with the uncertainty of the determination of the dose (16% relative error).

#### SI-11 Error propagation

The statistical error from sequencing is estimated by using the standard deviation of a Bernoulli distribution $stddev \equiv \Delta \sim \frac{1}{\sqrt{\bar{H}^{(0)}(i,k)}}$ and then propagated through all operations of the data analysis shown above. The standard error propagation for an operation $f(x_1..x_g)$ with $x_1..x_g$ being the used datasets, $\Delta_f$ being the propagated error and $\Delta_{x_1},..,\Delta_{x_g}$ being the error of the datasets $x_1..x_g$ is then given by[6]:

(SI-e 49) $$\Delta_f^2 \sim \Delta_{x_1}^2 \cdot \left(\frac{\delta f}{\delta x_1}\right)^2 + .. + \Delta_{x_g}^2 \left(\frac{\delta f}{\delta x_g}\right)^2 + 2 \cdot \sum_{r,t \in \{1..g\}} \Delta_{x_r,x_t}^2 \cdot \left(\frac{\delta f}{\delta x_r}\right) \cdot \left(\frac{\delta f}{\delta x_t}\right).$$

The propagated error is then used to calculate the weights for fitting the state models as shown above. The resulting fitting errors for the rate parameters $\Delta_{\rho_{i_c}}$ , $\Delta_{\sigma_{i_c}}$, $\Delta_{\nu_{i_c}}$are calculated via error propagation via the conversion functions SI-e 41-43. The errors of the fit parameters $\Delta_A, \Delta_\gamma$ and cross correlations are given by the fitting algorithm.

#### SI-12 Example for data handling

In this paragraph we present a typical example for the computation of the dose dependencies from the original Illumina data. The experiment yields the absolute frequencies of the oligomers with a specific sequence $i$ in the random part for the different samples $j$. The samples are numbered according to the handling number $j$ in the analysis procedure. The raw data for the frequency of the central octamer (only oligomers with intact ACAC-tails are counted) are given in Table SI-t 2(lower part). From the $4^8 = 65536$ sequencies of the oligomer we show only 20 starting with AAAAAAAA, AAAAAAAC, etc. Also, a selection of the sample numbers is given. The corresponding dose values are shown in the upper part of Table SI-t 2. Looking at the displayed values we find absolute frequencies between 0 and 4075 counts. Striking is the absence of counts for the sample with $j = 1$. Apparently, a fatal error occurred in the preparation/analysis process of this sample. This is confirmed by the analysis in chapter SI-13 where we demonstrated that the values $j = 1$ and $j = 6$ are outliers of the experiment and should be disregarded. The data in Table SI-t 2show that this is trivial for $j = 1$. For the frequencies of the sample $j = 6$ non-vanishing but small values occur. Even smaller absolute frequencies are found in the raw $j = 9$ to $j = 11$, however these small values may be explained by the strong illumination doses (see upper part).

For the central octamer, the defined ACAC-tails lead to defined neighbors at the 5' (C) and 3' (A) ends of the octamer part. Since important DNA-lesions such as the CPD-photolesion involve adjacent bases, the tail sequences do not allow to use the given octamer data as a prototype for DNA-damage in random sequences. This is only achieved when the octamer data are averaged or summed over the outmost bases at position 0 and 7. The resulting oligomer, i.e., hexamer data (see Table SI-t 3) have always a random base outside the hexamer. Due to the summation over the terminal bases of the octamer, the related absolute frequencies are larger than for the octamer (approximately by one order of magnitude). Again, the outliers and the small absolute frequencies for the samples with remarkably high doses are clearly visible. Another striking feature is strong variation of the absolute frequencies with sequence for constant sample numbers (or doses). For given sequences there is the trend that the absolute frequencies decay with increasing dose.

The absolute frequencies may further be increased upon appropriate summation of hexamer sequences leading to shorter oligomers of defined length (from pentamers to dimers). An example is given in Table SI-t 4, where the absolute frequencies for dimers are presented. The frequency values are now in the range of $10^6$. The sequence dependence is also visible in the dimer data and may be due to accidental sequence dependencies in the synthesis and the analysis/preparation procedure. It is visible for all dimer sequences. According to Equation SI-e 3 the sequence

dependence may be eliminated by normalizing the absolute frequencies of a given sample to those of a reference sample. We chose the sample with vanishing dose, $D_j = 0$, since the expected sequence dependence induced by illumination should be absent for this sample. By this normalization procedure, we obtain the normalized absolute frequencies displayed in Table SI-t 5. Here the sequence dependence is weak for the small dose values. At high doses, a pronounced sequence dependence is visible and points to a sequence dependence induced by illumination.
Until now the variations in the sample preparation/analysis steps, i.e. during library preparation and Illumina analysis are not yet corrected for. They may influence the determined relative frequencies and may differ from sample to sample. In first order approximation they should be independent of the sequence. An additional, strong, and systematic sample dependence visible in Table SI-t 5 is related to the illumination induced damage leading to very small relative frequencies at high doses. The illumination dependence of the relative frequencies is due to the ACAC-tails, the random parts outside the considered dimer and the dimer (within a surroundings of random bases) itself. According to Equation SI-e 6 the survival probabilities of these three parts must be multiplied to obtain the survival probability of the measured 16-mer.

According to Equation SI-e 4 and 5 we may eliminate the influences of tails and random parts by normalizing the relative frequencies by the relative frequencies of one sequence $i_0$ where we expect a weak damage due to the dimer part. The resulting normalized relative frequencies (see Table SI-t 6) are obtained by using $i_0$ for the dimer $GG$. These relative frequencies should represent the true relative frequencies multiplied by the inverse of the decay of the dimer part (see Equation SI-e 5). Thus, the data for the dimer $GG$ are equal to 1. To obtain the true dose dependence, one needs knowledge of the decay $F_{i0}$ of the dimer $GG$. It is well known that the related decay should be rather weak. An upper limit of the decay constant $D_{max}$ may be obtained from the fact that radiation induced processes of intact oligomers never lead to an increase of the frequencies. $D_{max}$ is reached when the multiplication of the relative frequencies of Table SI-t 6 with $\exp\left(-\frac{D_j}{D_{max}}\right)$ yields only decaying curves. Further reduction of the decay constant $D_{max}$ may be obtained when additional information is available for the damage constants of specific oligomers. Another approach is used in Chapter SI-3 where the assumption of a defined dimer damaging coefficient $\Phi_{XY}$ is used for normalization.
For the present data set we assume $D_{max-GG} = 1890$ photon/base and obtain the relative frequencies given in Table SI-t 7. For the tetramer and the hexamer we obtained: $D_{max-GGGG} = 1300$ and $D_{max-GGGGGG} = 990$ photon/base.

**Table SI-t 2.** Absolute frequencies for the raw data set for the doses $D_j$ given in the upper part. The outliers are j = 1 and j = 6.

| Sample# | j = | Dose $D_j$ (photon/base)<br>0 | 1 | 2 | 3 | 4 | 5 | 6 | 7 | 8 | 9 | 10 | 11 | 12 |
|---|---|---|---|---|---|---|---|---|---|---|---|---|---|---|
| | | 0.00 | 15.52 | 22.14 | 31.02 | 44.46 | 63.22 | 91.74 | 126.99 | 184.12 | 261.25 | 369.49 | 558.61 | 1.92 |

| Sequence number i | Sequence | Frequency in Sample j =<br>0 | 1 | 2 | 3 | 4 | 5 | 6 | 7 | 8 | 9 | 10 | 11 | 12 | ... |
|---|---|---|---|---|---|---|---|---|---|---|---|---|---|---|---|
| 1 | AAAAAAAA | 4075 | 0 | 1804 | 1933 | 553 | 405 | 25 | 82 | 32 | 5 | 0 | 0 | 3521 | ... |
| 2 | AAAAAAAC | 3928 | 0 | 1792 | 1985 | 656 | 509 | 57 | 143 | 65 | 11 | 1 | 0 | 3179 | ... |
| 3 | AAAAAAAG | 1546 | 0 | 752 | 847 | 254 | 241 | 34 | 51 | 34 | 5 | 0 | 0 | 1335 | ... |
| 4 | AAAAAAAT | 3299 | 0 | 1613 | 1726 | 501 | 430 | 9 | 130 | 44 | 2 | 0 | 0 | 2725 | ... |
| 5 | AAAAAACA | 2724 | 0 | 1280 | 1522 | 474 | 412 | 39 | 124 | 39 | 11 | 0 | 0 | 2363 | ... |
| 6 | AAAAAACC | 2088 | 0 | 1017 | 1111 | 375 | 295 | 39 | 105 | 54 | 6 | 0 | 0 | 1736 | ... |
| 7 | AAAAAACG | 1609 | 0 | 829 | 864 | 310 | 241 | 36 | 86 | 29 | 7 | 0 | 0 | 1396 | ... |
| 8 | AAAAAACT | 1927 | 0 | 866 | 918 | 294 | 228 | 8 | 87 | 29 | 14 | 0 | 0 | 1551 | ... |
| 9 | AAAAAAGA | 1813 | 0 | 886 | 905 | 286 | 271 | 28 | 76 | 35 | 8 | 0 | 0 | 1485 | ... |
| 10 | AAAAAAGC | 1663 | 0 | 770 | 897 | 280 | 228 | 31 | 102 | 43 | 9 | 0 | 0 | 1325 | ... |
| 11 | AAAAAAGG | 581 | 0 | 282 | 339 | 105 | 94 | 11 | 32 | 9 | 3 | 0 | 0 | 491 | ... |
| 12 | AAAAAAGT | 1213 | 0 | 576 | 676 | 174 | 204 | 12 | 55 | 36 | 7 | 0 | 0 | 998 | ... |
| 13 | AAAAAATA | 3150 | 0 | 1533 | 1740 | 468 | 436 | 26 | 111 | 40 | 7 | 0 | 0 | 2686 | ... |
| 14 | AAAAAATC | 2620 | 0 | 1285 | 1366 | 432 | 397 | 31 | 138 | 60 | 16 | 1 | 0 | 2264 | ... |
| 15 | AAAAAATG | 1559 | 0 | 772 | 842 | 245 | 236 | 23 | 70 | 39 | 6 | 0 | 0 | 1239 | ... |
| 16 | AAAAAATT | 2462 | 0 | 880 | 952 | 250 | 212 | 2 | 67 | 30 | 7 | 0 | 0 | 1959 | ... |
| 17 | AAAAACAA | 2021 | 0 | 1002 | 1108 | 352 | 311 | 41 | 96 | 34 | 2 | 0 | 0 | 1770 | ... |
| 18 | AAAAACAC | 1830 | 0 | 938 | 994 | 381 | 319 | 36 | 138 | 55 | 19 | 0 | 0 | 1471 | ... |
| 19 | AAAAACAG | 765 | 0 | 430 | 515 | 152 | 171 | 33 | 49 | 19 | 8 | 1 | 0 | 754 | ... |
| 20 | AAAAACAT | 1418 | 0 | 689 | 866 | 272 | 224 | 6 | 107 | 53 | 10 | 2 | 0 | 1239 | ... |
| ... | ... | ... | ... | ... | ... | ... | ... | ... | ... | ... | ... | ... | ... | ... | ... |

**Table SI-t 3.** Absolute frequencies of the central hexamer, obtained by summing over the base at position 0 and 7 of the central octamer. The doses $D_j$ are given in the upper part. The outliers are j = 1 and j = 6.

| | | Dose $D_j$ (photon/base) | | | | | | | | | | | | |
|---|---|---|---|---|---|---|---|---|---|---|---|---|---|---|
| Sample# | j = | 0 | 1 | 2 | 3 | 4 | 5 | 6 | 7 | 8 | 9 | 10 | 11 | 12 |
| | | 0.00 | 15.52 | 22.14 | 31.02 | 44.46 | 63.22 | 91.74 | 126.99 | 184.12 | 261.25 | 369.49 | 558.61 | 1.92 |

| Sequence | | Frequency in Sample j = | | | | | | | | | | | | | |
|---|---|---|---|---|---|---|---|---|---|---|---|---|---|---|---|
| number | Sequence | 0 | 1 | 2 | 3 | 4 | 5 | 6 | 7 | 8 | 9 | 10 | 11 | 12 | ... |
| 1 | AAAAAA | 32221 | 0 | 14772 | 16060 | 4642 | 3770 | 278 | 1038 | 431 | 66 | 2 | 1 | 26859 | ... |
| 2 | AAAAAC | 20509 | 0 | 9713 | 10456 | 3277 | 2759 | 292 | 965 | 405 | 75 | 3 | 0 | 16842 | ... |
| 3 | AAAAAG | 13260 | 0 | 6443 | 7222 | 2131 | 1989 | 195 | 655 | 322 | 85 | 3 | 3 | 10865 | ... |
| 4 | AAAAAT | 24656 | 0 | 11048 | 11973 | 3354 | 2900 | 196 | 933 | 394 | 84 | 6 | 0 | 20180 | ... |
| 5 | AAAACA | 14605 | 0 | 7358 | 8231 | 2662 | 2419 | 265 | 976 | 420 | 111 | 10 | 0 | 12391 | ... |
| 6 | AAAACC | 8172 | 0 | 3912 | 4203 | 1471 | 1212 | 138 | 483 | 226 | 56 | 3 | 1 | 6630 | ... |
| 7 | AAAACG | 9095 | 0 | 4683 | 5053 | 1666 | 1439 | 156 | 584 | 284 | 69 | 7 | 0 | 7557 | ... |
| 8 | AAAACT | 10544 | 0 | 4525 | 4743 | 1403 | 1226 | 105 | 478 | 185 | 46 | 4 | 2 | 8528 | ... |
| 9 | AAAAGA | 14069 | 0 | 7113 | 8035 | 2578 | 2345 | 209 | 988 | 469 | 108 | 12 | 4 | 11733 | ... |
| 10 | AAAAGC | 8282 | 0 | 4131 | 4612 | 1580 | 1308 | 179 | 605 | 278 | 75 | 8 | 0 | 6509 | ... |
| 11 | AAAAGG | 5318 | 0 | 2689 | 3118 | 925 | 908 | 83 | 355 | 191 | 44 | 10 | 0 | 4391 | ... |
| 12 | AAAAGT | 7535 | 0 | 3612 | 4187 | 1302 | 1162 | 115 | 558 | 280 | 74 | 12 | 1 | 6097 | ... |
| 13 | AAAATA | 19433 | 0 | 9534 | 10878 | 3379 | 2846 | 153 | 1175 | 489 | 111 | 20 | 0 | 15929 | ... |
| 14 | AAAATC | 11210 | 0 | 5205 | 5544 | 1878 | 1647 | 127 | 635 | 293 | 75 | 7 | 0 | 9259 | ... |
| 15 | AAAATG | 10358 | 0 | 5145 | 5976 | 1905 | 1721 | 118 | 689 | 364 | 78 | 4 | 0 | 8746 | ... |
| 16 | AAAATT | 15326 | 0 | 4988 | 5087 | 1314 | 1123 | 48 | 397 | 190 | 29 | 2 | 0 | 12077 | ... |
| 17 | AAACAA | 13508 | 0 | 6987 | 7672 | 2501 | 2320 | 243 | 994 | 496 | 93 | 9 | 1 | 11465 | ... |
| 18 | AAACAC | 8154 | 0 | 4389 | 4572 | 1730 | 1485 | 186 | 703 | 379 | 128 | 14 | 0 | 6671 | ... |
| 19 | AAACAG | 5862 | 0 | 3127 | 3527 | 1215 | 1162 | 109 | 593 | 270 | 85 | 10 | 1 | 4955 | ... |
| 20 | AAACAT | 10033 | 0 | 4833 | 5334 | 1799 | 1640 | 121 | 795 | 396 | 118 | 13 | 0 | 8412 | ... |
| ... | ... | ... | ... | ... | ... | ... | ... | ... | ... | ... | ... | ... | ... | ... | ... |

**Table SI-t 4.** Absolute frequencies of the dimers obtained by summing over the appropriate parts of the central hexamer. The doses $D_j$ are given in the upper part. The outliers are j = 1 and j = 6.

| Sample# | j = | Dose $D_j$ (photon/base) 0 | 1 | 2 | 3 | 4 | 5 | 6 | 7 | 8 | 9 | 10 | 11 | 12 ... |
|---|---|---|---|---|---|---|---|---|---|---|---|---|---|---|
| | | 0.00 | 15.52 | 22.14 | 31.02 | 44.46 | 63.22 | 91.74 | 126.99 | 184.12 | 261.25 | 369.49 | 558.61 | 1.92 |

| Sequence number i | Sequence | Absolute Frequency in Sample j = 0 | 1 | 2 | 3 | 4 | 5 | 6 | 7 | 8 | 9 | 10 | 11 | 12 ... |
|---|---|---|---|---|---|---|---|---|---|---|---|---|---|---|
| 1 | AA | 5.924e+06 | 0.000e+00 | 2.858e+06 | 3.175e+06 | 1.057e+06 | 9.604e+05 | 7.910e+04 | 4.518e+05 | 2.577e+05 | 8.068e+04 | 1.018e+04 | 8.080e+02 | 4.886e+06 |
| 2 | AC | 2.372e+06 | 0.000e+00 | 1.172e+06 | 1.300e+06 | 4.710e+05 | 4.347e+05 | 3.917e+04 | 2.357e+05 | 1.517e+05 | 5.584e+04 | 8.663e+03 | 8.590e+02 | 1.950e+06 |
| 3 | AG | 2.144e+06 | 0.000e+00 | 1.122e+06 | 1.272e+06 | 4.350e+05 | 4.222e+05 | 3.537e+04 | 2.343e+05 | 1.596e+05 | 5.955e+04 | 9.382e+03 | 9.480e+02 | 1.783e+06 |
| 4 | AT | 3.615e+06 | 0.000e+00 | 1.669e+06 | 1.863e+06 | 6.401e+05 | 5.873e+05 | 4.412e+04 | 3.143e+05 | 2.000e+05 | 7.211e+04 | 1.104e+04 | 1.008e+03 | 2.969e+06 |
| 5 | CA | 2.068e+06 | 0.000e+00 | 1.027e+06 | 1.147e+06 | 4.143e+05 | 3.887e+05 | 3.396e+04 | 2.157e+05 | 1.421e+05 | 5.430e+04 | 8.999e+03 | 9.480e+02 | 1.705e+06 |
| 6 | CC | 7.880e+05 | 0.000e+00 | 3.683e+05 | 3.988e+05 | 1.534e+05 | 1.375e+05 | 1.243e+04 | 7.616e+04 | 4.968e+04 | 1.911e+04 | 3.201e+03 | 3.960e+02 | 6.384e+05 |
| 7 | CG | 9.943e+05 | 0.000e+00 | 5.052e+05 | 5.639e+05 | 2.015e+05 | 1.903e+05 | 1.619e+04 | 1.075e+05 | 7.250e+04 | 2.796e+04 | 4.552e+03 | 4.850e+02 | 8.150e+05 |
| 8 | CT | 1.202e+06 | 0.000e+00 | 5.017e+05 | 5.469e+05 | 1.974e+05 | 1.744e+05 | 1.336e+04 | 9.335e+04 | 5.923e+04 | 2.170e+04 | 3.506e+03 | 3.890e+02 | 9.600e+05 |
| 9 | GA | 2.573e+06 | 0.000e+00 | 1.325e+06 | 1.495e+06 | 5.142e+05 | 4.924e+05 | 3.771e+04 | 2.683e+05 | 1.807e+05 | 6.654e+04 | 1.020e+04 | 9.760e+02 | 2.130e+06 |
| 10 | GC | 9.227e+05 | 0.000e+00 | 4.717e+05 | 5.262e+05 | 1.890e+05 | 1.807e+05 | 1.537e+04 | 1.059e+05 | 7.351e+04 | 2.943e+04 | 5.038e+03 | 5.420e+02 | 7.526e+05 |
| 11 | GG | 9.100e+05 | 0.000e+00 | 4.805e+05 | 5.467e+05 | 1.837e+05 | 1.801e+05 | 1.307e+04 | 1.029e+05 | 7.418e+04 | 2.916e+04 | 5.146e+03 | 5.310e+02 | 7.507e+05 |
| 12 | GT | 1.023e+06 | 0.000e+00 | 5.077e+05 | 5.746e+05 | 1.982e+05 | 1.857e+05 | 1.489e+04 | 1.077e+05 | 7.598e+04 | 2.907e+04 | 5.106e+03 | 5.830e+02 | 8.355e+05 |
| 13 | TA | 3.191e+06 | 0.000e+00 | 1.475e+06 | 1.650e+06 | 5.740e+05 | 5.253e+05 | 4.145e+04 | 2.810e+05 | 1.794e+05 | 6.382e+04 | 9.780e+03 | 9.140e+02 | 2.618e+06 |
| 14 | TC | 1.231e+06 | 0.000e+00 | 5.186e+05 | 5.646e+05 | 2.055e+05 | 1.845e+05 | 1.554e+04 | 9.878e+04 | 6.238e+04 | 2.329e+04 | 3.864e+03 | 4.380e+02 | 9.856e+05 |
| 15 | TG | 1.293e+06 | 0.000e+00 | 6.353e+05 | 7.160e+05 | 2.510e+05 | 2.352e+05 | 1.779e+04 | 1.351e+05 | 9.496e+04 | 3.680e+04 | 6.297e+03 | 6.640e+02 | 1.062e+06 |
| 16 | TT | 2.046e+06 | 0.000e+00 | 6.304e+05 | 6.373e+05 | 2.076e+05 | 1.778e+05 | 1.177e+04 | 9.065e+04 | 5.852e+04 | 2.152e+04 | 3.497e+03 | 4.110e+02 | 1.607e+06 |

**Table SI-t 5**. Relative frequencies of the dimers, obtained from the data of Table SI-T4 by normalization to the absolute frequencies of the sample j = 0 with $D_0$ = 0. The doses $D_j$ are given in the upper part. The outliers are j = 1 and j = 6.

| | Dose $D_i$ (photon/base) | | | | | | | | | | | | |
|---|---|---|---|---|---|---|---|---|---|---|---|---|---|
| Sample# | 0 | 1 | 2 | 3 | 4 | 5 | 6 | 7 | 8 | 9 | 10 | 11 | 12 ... |
| | 0.00 | 15.52 | 22.14 | 31.02 | 44.46 | 63.22 | 91.74 | 126.99 | 184.12 | 261.25 | 369.49 | 558.61 | 1.92 |

| | Frequency in Sample j = | | | | | | | | | | | | |
|---|---|---|---|---|---|---|---|---|---|---|---|---|---|
| Sequence | 0 | 1 | 2 | 3 | 4 | 5 | 6 | 7 | 8 | 9 | 10 | 11 | 12 ... |
| AA | 1 | 0 | 0.48250 | 0.53587 | 0.17843 | 0.16212 | 0.01335 | 0.07626 | 0.04350 | 0.01362 | 0.00172 | 0.00014 | 0.82470 |
| AC | 1 | 0 | 0.49397 | 0.54832 | 0.19859 | 0.18330 | 0.01652 | 0.09939 | 0.06398 | 0.02354 | 0.00365 | 0.00036 | 0.82226 |
| AG | 1 | 0 | 0.52338 | 0.59326 | 0.20292 | 0.19692 | 0.01650 | 0.10930 | 0.07444 | 0.02778 | 0.00438 | 0.00044 | 0.83164 |
| AT | 1 | 0 | 0.46170 | 0.51536 | 0.17709 | 0.16247 | 0.01221 | 0.08697 | 0.05535 | 0.01995 | 0.00305 | 0.00028 | 0.82149 |
| CA | 1 | 0 | 0.49636 | 0.55467 | 0.20033 | 0.18793 | 0.01642 | 0.10428 | 0.06870 | 0.02626 | 0.00435 | 0.00046 | 0.82455 |
| CC | 1 | 0 | 0.46735 | 0.50611 | 0.19464 | 0.17452 | 0.01578 | 0.09665 | 0.06304 | 0.02426 | 0.00406 | 0.00050 | 0.81009 |
| CG | 1 | 0 | 0.50815 | 0.56714 | 0.20269 | 0.19140 | 0.01629 | 0.10817 | 0.07292 | 0.02812 | 0.00458 | 0.00049 | 0.81973 |
| CT | 1 | 0 | 0.41720 | 0.45482 | 0.16414 | 0.14500 | 0.01111 | 0.07763 | 0.04926 | 0.01805 | 0.00292 | 0.00032 | 0.79838 |
| GA | 1 | 0 | 0.51515 | 0.58103 | 0.19988 | 0.19139 | 0.01466 | 0.10429 | 0.07023 | 0.02586 | 0.00396 | 0.00038 | 0.82806 |
| GC | 1 | 0 | 0.51121 | 0.57024 | 0.20483 | 0.19579 | 0.01666 | 0.11473 | 0.07967 | 0.03190 | 0.00546 | 0.00059 | 0.81570 |
| GG | 1 | 0 | 0.52805 | 0.60076 | 0.20193 | 0.19794 | 0.01436 | 0.11312 | 0.08152 | 0.03204 | 0.00566 | 0.00058 | 0.82499 |
| GT | 1 | 0 | 0.49613 | 0.56155 | 0.19371 | 0.18143 | 0.01456 | 0.10525 | 0.07426 | 0.02841 | 0.00499 | 0.00057 | 0.81654 |
| TA | 1 | 0 | 0.46213 | 0.51710 | 0.17989 | 0.16464 | 0.01299 | 0.08807 | 0.05622 | 0.02000 | 0.00307 | 0.00029 | 0.82054 |
| TC | 1 | 0 | 0.42135 | 0.45875 | 0.16696 | 0.14991 | 0.01263 | 0.08026 | 0.05068 | 0.01892 | 0.00314 | 0.00036 | 0.80074 |
| TG | 1 | 0 | 0.49149 | 0.55395 | 0.19415 | 0.18194 | 0.01376 | 0.10456 | 0.07347 | 0.02847 | 0.00487 | 0.00051 | 0.82127 |
| TT | 1 | 0 | 0.30802 | 0.31142 | 0.10145 | 0.08689 | 0.00575 | 0.04430 | 0.02860 | 0.01051 | 0.00171 | 0.00020 | 0.78533 |

**Table SI-t 6.** Relative frequencies of the dimers, obtained from the data of Table SI-T5 by normalization to the relative frequencies of the sequence GG, i = 10. The doses $D_j$ are given in the upper part. The outliers are j = 1 and j = 6.

| | Dose $D_j$ (photon/base) | | | | | | | | | | | | |
|---|---|---|---|---|---|---|---|---|---|---|---|---|---|
| Sample# | 0 | 1 | 2 | 3 | 4 | 5 | 6 | 7 | 8 | 9 | 10 | 11 | 12 ... |
| | 0.00 | 15.52 | 22.14 | 31.02 | 44.46 | 63.22 | 91.74 | 126.99 | 184.12 | 261.25 | 369.49 | 558.61 | 1.92 |

| | Frequency in Sample j = | | | | | | | | | | | | |
|---|---|---|---|---|---|---|---|---|---|---|---|---|---|
| Sequence | 0 | 1 | 2 | 3 | 4 | 5 | 6 | 7 | 8 | 9 | 10 | 11 | 12 ... |
| AA | 1.00000 | - | 0.91374 | 0.89199 | 0.88366 | 0.81904 | 0.92975 | 0.67415 | 0.53362 | 0.42499 | 0.30375 | 0.23359 | 0.99965 |
| AC | 1.00000 | - | 0.93546 | 0.91271 | 0.98348 | 0.92608 | 1.15006 | 0.87861 | 0.78478 | 0.73469 | 0.64585 | 0.62023 | 0.99669 |
| AG | 1.00000 | - | 0.99115 | 0.98751 | 1.00495 | 0.99488 | 1.14891 | 0.96626 | 0.91316 | 0.86684 | 0.77379 | 0.75716 | 1.00807 |
| AT | 1.00000 | - | 0.87434 | 0.85785 | 0.87699 | 0.82081 | 0.84999 | 0.76879 | 0.67889 | 0.62253 | 0.54008 | 0.47747 | 0.99575 |
| CA | 1.00000 | - | 0.93999 | 0.92327 | 0.99212 | 0.94945 | 1.14361 | 0.92188 | 0.84265 | 0.81938 | 0.76939 | 0.78490 | 0.99947 |
| CC | 1.00000 | - | 0.88504 | 0.84245 | 0.96392 | 0.88170 | 1.09855 | 0.85437 | 0.77330 | 0.75696 | 0.71838 | 0.86172 | 0.98194 |
| CG | 1.00000 | - | 0.96231 | 0.94403 | 1.00378 | 0.96698 | 1.13416 | 0.95625 | 0.89451 | 0.87747 | 0.80961 | 0.83610 | 0.99362 |
| CT | 1.00000 | - | 0.79007 | 0.75708 | 0.81290 | 0.73255 | 0.77385 | 0.68628 | 0.60419 | 0.56316 | 0.51564 | 0.55477 | 0.96775 |
| GA | 1.00000 | - | 0.97556 | 0.96716 | 0.98988 | 0.96694 | 1.02080 | 0.92194 | 0.86143 | 0.80712 | 0.70073 | 0.64956 | 1.00373 |
| GC | 1.00000 | - | 0.96811 | 0.94919 | 1.01438 | 0.98914 | 1.15978 | 1.01419 | 0.97728 | 0.99550 | 0.96551 | 1.00659 | 0.98874 |
| GG | 1.00000 | - | 1.00000 | 1.00000 | 1.00000 | 1.00000 | 1.00000 | 1.00000 | 1.00000 | 1.00000 | 1.00000 | 1.00000 | 1.00000 |
| GT | 1.00000 | - | 0.93956 | 0.93473 | 0.95932 | 0.91664 | 1.01362 | 0.93045 | 0.91085 | 0.88651 | 0.88238 | 0.97621 | 0.98976 |
| TA | 1.00000 | - | 0.87517 | 0.86074 | 0.89086 | 0.83180 | 0.90466 | 0.77853 | 0.68963 | 0.62415 | 0.54197 | 0.49052 | 0.99460 |
| TC | 1.00000 | - | 0.79794 | 0.76361 | 0.82685 | 0.75739 | 0.87935 | 0.70948 | 0.62171 | 0.59057 | 0.55518 | 0.61008 | 0.97060 |
| TG | 1.00000 | - | 0.93076 | 0.92208 | 0.96151 | 0.91918 | 0.95835 | 0.92429 | 0.90122 | 0.88840 | 0.86145 | 0.88002 | 0.99550 |
| TT | 1.00000 | - | 0.58332 | 0.51838 | 0.50239 | 0.43898 | 0.40045 | 0.39160 | 0.35080 | 0.32809 | 0.30220 | 0.34436 | 0.95193 |

**Table SI-t 7.** Relative frequencies of the dimers, obtained by multiplying the data of Table SI-T6 by the assumed decay ($\exp(-D_j/2000)$) of the frequencies of sequence GG,. The doses $D_j$ are given in the upper part. The outliers are j = 1 and j = 6.

| Sample# | Dose $D_i$ (photon/base) 0 | 1 | 2 | 3 | 4 | 5 | 6 | 7 | 8 | 9 | 10 | 11 | 12 ... |
|---|---|---|---|---|---|---|---|---|---|---|---|---|---|
| | 0.00 | 15.52 | 22.14 | 31.02 | 44.46 | 63.22 | 91.74 | 126.99 | 184.12 | 261.25 | 369.49 | 558.61 | 1.92 |

| Sequence | Frequency in Sample j = 0 | 1 | 2 | 3 | 4 | 5 | 6 | 7 | 8 | 9 | 10 | 11 | 12 ... |
|---|---|---|---|---|---|---|---|---|---|---|---|---|---|
| AA | 1 | - | 0.903093 | 0.877475 | 0.863113 | 0.792096 | 0.885695 | 0.630339 | 0.484091 | 0.370127 | 0.249812 | 0.173814 | 0.998636 |
| AC | 1 | - | 0.92456 | 0.897851 | 0.960613 | 0.895613 | 1.09557 | 0.821516 | 0.711931 | 0.639846 | 0.531159 | 0.461521 | 0.995679 |
| AG | 1 | - | 0.979607 | 0.971435 | 0.981584 | 0.962153 | 1.09447 | 0.90347 | 0.828395 | 0.754929 | 0.636384 | 0.563418 | 1.00704 |
| AT | 1 | - | 0.864153 | 0.843887 | 0.856603 | 0.793813 | 0.809715 | 0.718834 | 0.615872 | 0.542158 | 0.444178 | 0.355293 | 0.994743 |
| CA | 1 | - | 0.929037 | 0.908247 | 0.96905 | 0.918215 | 1.08943 | 0.861976 | 0.764438 | 0.713599 | 0.632764 | 0.584053 | 0.998455 |
| CC | 1 | - | 0.874729 | 0.828734 | 0.941509 | 0.852694 | 1.0465 | 0.798855 | 0.701523 | 0.659238 | 0.590813 | 0.641218 | 0.980946 |
| CG | 1 | - | 0.951097 | 0.928664 | 0.980446 | 0.935173 | 1.08043 | 0.894108 | 0.811476 | 0.764193 | 0.665841 | 0.622151 | 0.99261 |
| CT | 1 | - | 0.780863 | 0.744755 | 0.793996 | 0.708454 | 0.737185 | 0.641683 | 0.548106 | 0.49046 | 0.424073 | 0.412815 | 0.966767 |
| GA | 1 | - | 0.9642 | 0.951415 | 0.966865 | 0.935138 | 0.97243 | 0.86203 | 0.781474 | 0.702918 | 0.5763 | 0.483345 | 1.0027 |
| GC | 1 | - | 0.956831 | 0.933741 | 0.990801 | 0.956605 | 1.10482 | 0.948282 | 0.886568 | 0.866984 | 0.794057 | 0.749021 | 0.987735 |
| GG | 1 | - | 0.988352 | 0.983723 | 0.976751 | 0.967106 | 0.952619 | 0.935018 | 0.907178 | 0.870901 | 0.822425 | 0.744115 | 0.998984 |
| GT | 1 | - | 0.928611 | 0.919516 | 0.937016 | 0.886484 | 0.965591 | 0.869991 | 0.826307 | 0.772065 | 0.725689 | 0.726414 | 0.98875 |
| TA | 1 | - | 0.864972 | 0.846734 | 0.870151 | 0.804434 | 0.861793 | 0.72794 | 0.625617 | 0.543573 | 0.445731 | 0.365003 | 0.993591 |
| TC | 1 | - | 0.788642 | 0.751181 | 0.80763 | 0.732476 | 0.837684 | 0.663378 | 0.564006 | 0.51433 | 0.45659 | 0.453971 | 0.969618 |
| TG | 1 | - | 0.919914 | 0.907076 | 0.939158 | 0.888942 | 0.912944 | 0.864227 | 0.817568 | 0.773707 | 0.708479 | 0.654836 | 0.994485 |
| TT | 1 | - | 0.576527 | 0.509947 | 0.49071 | 0.424536 | 0.38148 | 0.366152 | 0.318239 | 0.285736 | 0.248533 | 0.25624 | 0.950963 |

#### SI-13 Detection of outliers

According Equation SI-e 1 the data of an experiment are represented by a two-dimensional set of numbers, the count numbers $H^{(1)}(i,j)$, obtained as a function of sequence number $i$ and sample number $j$. In general, the illumination dose is systematically varied for the different samples each having a specific dose $D_j$. However the preparation steps in the handling of the individual samples, such as e. g. the library preparation, the PCR-amplification, ... , may be subject to fluctuations in the handling procedure which may lead to variations in the count numbers. In a first order approximation one may assume that these variations are independent of the sequence and can be corrected in the data handling procedure.

The amount of the sample or preparation specific fluctuations has been investigated in a reference experiment, where two types of DNA-sequences are compared. Type 1: The standard DNA oligomers with ACAC tails and randomized octamers in the center. This solution is split into $N$ (here $N = 18$) samples illuminated with doses $D_j$. Type 2: DNA oligomers with GTTG tails and randomized octamers in the center. This solution is again split into $N$ ($N = 18$) samples but is not illuminated. Finally the DNA samples of both types are mixed (1 part of type 2 with 3 parts of type 1, in order to compensate for a reduction in oligomer numbers upon illumination) leading to $N = 18$ samples $S_j$ which are subsequently treated by the Illumina process and analyzed. For each sample $j$ we obtain the count numbers for both types of DNA oligomers, which experienced - besides illumination - the same treatment. Thus, the GTTG-tailed, non-illuminated oligomers may serve as reference molecules for the illuminated ACAC-tailed oligomers of the respective sample $S_j$. The GTTG-tailed oligomers also give an impression of the fluctuations imposed by the preparation of the samples.

In Figure SI-f 11we plotted the count numbers of the GTTG-tailed oligomers averaged over all oligomer sequences (blue) for the different sample numbers $j = 0$ to 17. The graph shows that strong variations of the average count numbers by more than an order of magnitude occur for the identically prepared GTTG-tailed DNA samples. The average count numbers $H_j$ vary between $H_1 = 0$ and $H_0 = 20.38$. Another small average $H_j$ is found for sample $j = 6$ with $H_6 = 0.072$. These samples are true outliers in the preparation and may be omitted in the data evaluation. The rest of the samples have average count numbers between 2.2 and 18.2. The role as outliers for points $j = 1$ and $j = 6$ is also supported by an analysis where the Pearson correlation coefficient referred to $j = 0$ ($D_0 = 0$) is calculated (red points in Figure SI-f 11a and c). The plots show, that the Pearson correlation coefficient may also be used to assign outliers in experiments. This observation is important for experiments without reference sequences.

In order to gain information on the additional influence of the illumination, the average count numbers $H_j$ of the ACAC-tailed oligomers (blue, filled dots) are plotted in Figure SI-f 11b versus sample number $j$. Due to the difference in concentration for oligomers of type 1 and type 2 (see above) the average count numbers for ACAC-tailed oligomers are often found at higher values. To compare both types of oligomers, the data for the average count numbers of GTTG-tailed oligomers were scaled by a factor of 4.84 (coinciding points for $j = 0$, black open dots). Again, there is a wide spread of the $h_j$ values reflecting the trend found for the non-illuminated GTTG-tailed oligomers (black). However, at some sample numbers (see e.g. $j = 7$ to

11) there are significantly reduced values. This can be understood considering the illumination induced damage which leads to a strong decrease in $H_j$ at high illumination dose $D_j$ (see violet points). When plotting the average count numbers versus dose (see Figure SI-f 3) one finds a strong decrease with dose together with overlaid fluctuations due to the preparation of the samples.

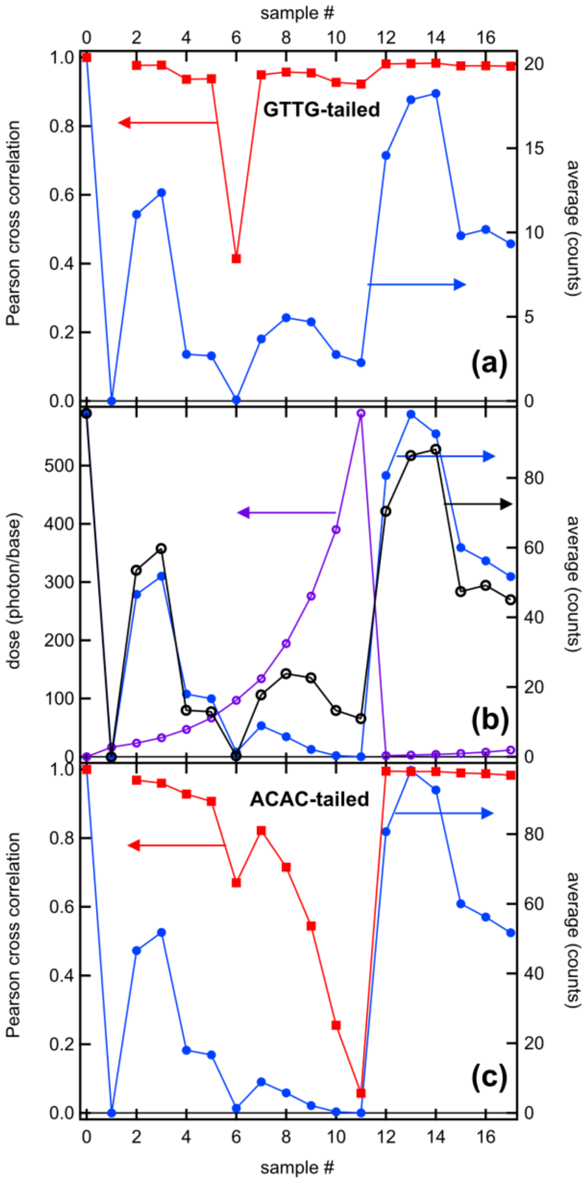

**Figure SI-f 11.** Data for the selection of outliers. (a) Average count numbers (blue dots) and Pearson correlation coefficient referred to $j = 0$ (red squares) for the reference oligomers (GTTG-tailed) plotted versus sample number $j$. (b) Dose applied to the ACAC-tailed oligomers as a function of sample number $j$ (violet circles). Average count numbers for both types of oligomers (blue dots, ACAC-tailed, black circles, GTTG-tailed, scaled by a factor of 4.84). (c) ACAC-tailed oligomers: Average count numbers (blue dots) and Pearson correlation coefficient referred to $j = 0$ (red squares). The figure clearly shows that the same samples are outliers in the experiments on GTTG-tailed and ACAC-tailed oligomers and that unusually small average count numbers and Pearson cross correlation values can be used to diagnose them.

#### SI-14 Analysis for the predominant occurrence of single-stranded DNA

For the calculations made before, it was assumed that the DNA is mostly in single strand form. Since the DNA pool contains a large number of different sequences and thus possible binding states, and the number of duplex formations will be quite small due to the short length of the DNA, a direct measurement of the fraction of bases involved in Watson-Crick (WC) pairing via UV absorption or intercalating dyes will only provide a rough estimate. Therefore, for a better understanding of the situation, we numerically calculated the binding probabilities for a large number of different sequences from the naïve pool. For this purpose, a local installation of the Nucleic Acid Package[7–9] (Nupack) was used and controlled in an automated way by means of a Labview interface in order to be able to calculate and evaluate the mutual interaction of a large number of sequences.
Nupack determines the possible binding configurations and their free energy for a group of sequences with a given concentration (1 nM per sequence) under predefined buffer conditions (0.135 M NaCl, no $MgCl_2$, 30°C). Due to the complexity of the problem, it is not possible to directly test the mutual interaction of $4^8 = 65536$ sequences against each other. Instead, 50 sequences were randomly selected from the complete pool for which all relevant binding permutations were calculated. This procedure was repeated 200 times to improve statistics and approximate reality. From the calculations, we obtained binding energies for $70 \cdot 10^3$ different duplex configurations (SI-f12a), for which a non-vanishing concentration averaging 0.5% of the strand concentration per sequence was obtained for approximately 25% (SI-f12b). Nupack also allows to determine the number of bases involved in Watson-Crick pairing per duplex formed, which in our example is on average 4 bases per 16-mer strand. Thus, only about 0.1 % of all bases are involved in WC binding, so that our assumption of mainly single-stranded DNA is plausible.

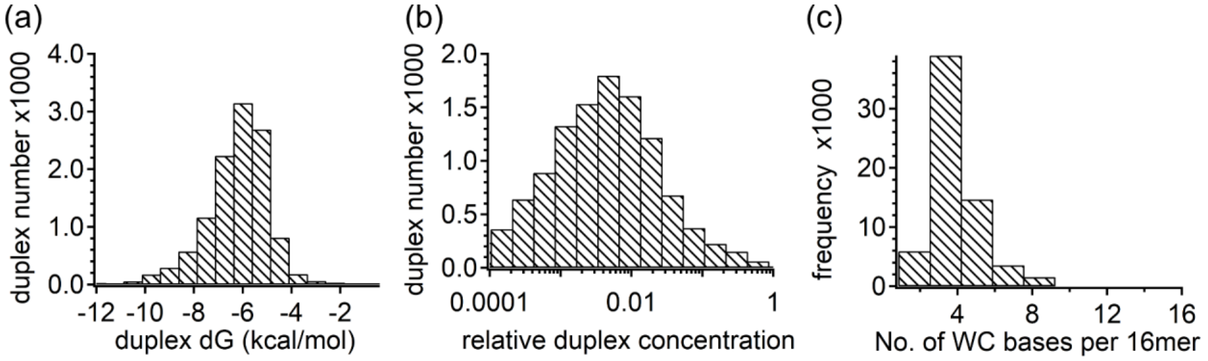

**Figure SI-f 12**: Calculated statistics on the binding probability in a randomized DNA pool for sequences ACAC NNNNNNNN ACAC. (a) After pairwise calculation of the possible binding energies of 103 different sequences each to form duplexes, a histogram is obtained for their mean binding energy. The mean free energy here is -6.3 kcal/mol. (b) From $70 \cdot 10^3$ different binding states considered, the concentration distribution shown here is obtained. At a strand concentration of 1 nM per sequence (1 mM base concentration), the average relative concentration of duplexes is 0.5%. (c) Histogram showing the number of bases participating in Watson-Crick binding for all duplexes. If one is formed, on average 4 bases are involved in the binding.

**SI-15 Determination of the damage yield for GG-dimers by ultraviolet spectroscopy**

Information on the damage yield of the reference dime GG was obtained from an experiment, where the change of the UV-spectrum of a GG-dimer sample was recorded for different illumination doses.
The sample: The dimer sample was obtained from Biomers, Germany. The sample was dissolved in phosphate buffer (10mM) to yield an absorbance of A ≈ 1 OD in the sample cell with an optical path length of 1 cm. The corresponding concentration $C_{GG}$ was estimated using the extinction coefficients given in Table SI-t 1: $C_{GG}$ = 47 μM.
The experimental system: Illumination of the sample was done with a Nd-based laser system (AOT-YVO-25QSP/MOPA) from Advanced Optical Technology, UK, operated at a repetition rate of 6.5 kHz. The UV radiation (4th harmonic of the laser) had an average power of ca. 20 mW with a beam diameter at the sample of ca. 1 mm. Double pass through the sample cell ascertained the essential part of the incoming light was absorbed. The residual transmission of the UV-radiation was considered when calculating the absorbed dose. For a set of illumination doses the absorption spectrum of the sample (1 cm path length) was recorded using a Shimadzu UV1900 spectrophotometer.
Results: In Figure SI-f13 we plotted the UV-absorption spectra of the GG sample before illumination (red) and after illumination with increasing dose values. The different spectra recorded at dose values until several 1000 photon/base show a remarkably similar shape. This indicates that there is only one process in this dose range, namely the decay of the initial absorption band towards a state with a weaker absorption and a wing extending to ca 350 nm. It requires more the 5000 photon/base to reduce the original absorption (e.g. at 266 nm, Figure SI-f 13) by 50%. The spectrum recorded at a dose of 650 photon/base shows, that on the dose level used in the illumination of the oligomers in the main paper, the concentration of the GG-dimer is essentially constant. This observation supports the assumption, that poly-G containing oligomers are well suited to serve as reference oligomers with smooth and very slowly decaying frequencies.

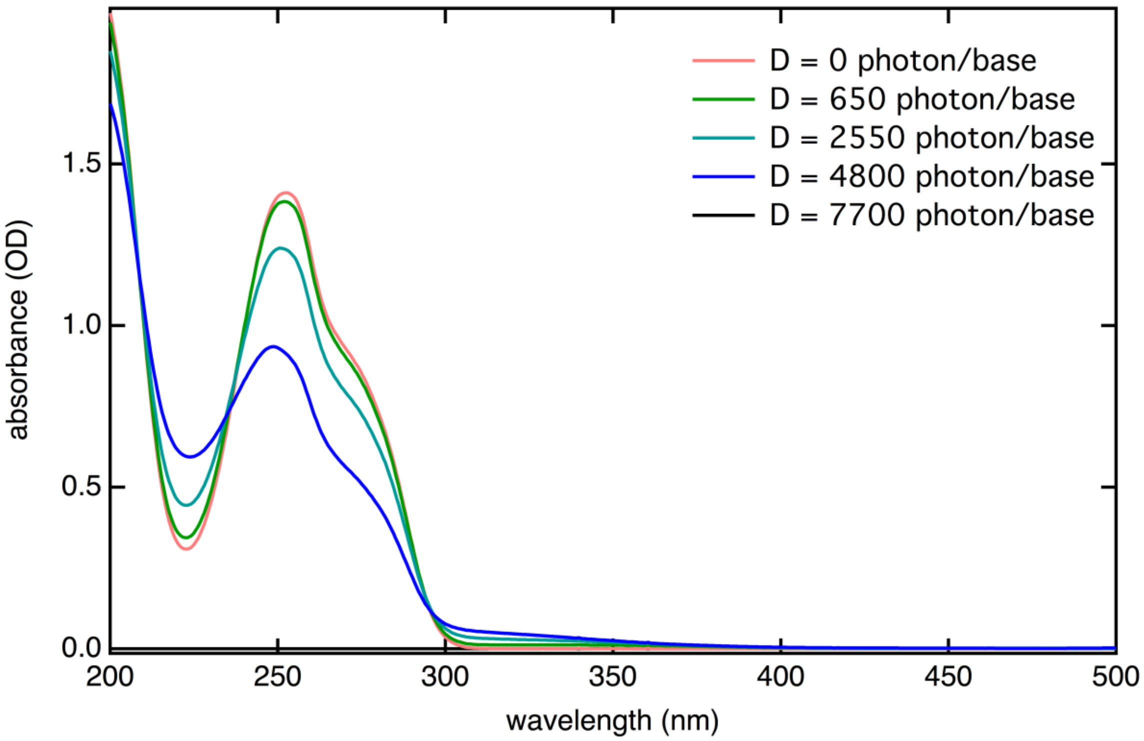

**Figure SI-f 13.** Plot of the absorption spectra of the GG-dimer sample in the near UV and visible range for different illumination doses indicated in the figure.

#### SI-16 Assessment of the sensitivity of the method for detection of 8-oxoguanine

To evaluate the sensitivity of the method or the polymerases used for further known structural damage of DNA, we performed measurements with DNA strands containing an 8-oxoguanine instead of a guanine at a central site.
Since this damage is monomeric, i.e. it involves only one nucleotide, it should be significantly less detectable by the polymerases used. Two samples were prepared for the measurement: One sample contains the modified target sequence ACAC AGAGAG ACAC and a reference sequence ACAC ACAGAGAC ACAC, each at a concentration of 5 µM in 1x PBS. The second sample also contains the reference sequence at identical concentrations and the modified target sequence ACAC AGA(8-oxo)GAGAG ACAC which has an 8-oxo guanine instead of the second guanine. Both samples were prepared for sequencing in the same way as the measurements shown throughout this work and were finally sequenced. For the evaluation, the sequence frequencies were each normalized by the frequencies of the reference sequence contained in the respective sample, so that both samples could be related to each other. Finally, all relative frequencies obtained in this way were normalized by the frequency of the unmodified target sequence. Figure SI-f14 shows on the left how the relative frequency of the target sequence drops from 1 in the sample without 8-oxo modification to 0.026 in the sample containing the 8-oxo modified target sequence. This detection amplitude of ca 30-times larger than the CPD damage detection in the case of a T=T CPD-lesion (main text Figure 2b). Moreover, the frequency of another sequence (ACAC AGA**T**AGAG ACAC) simultaneously increases from $10^{-6}$ (i.e. a single readout event) in the sample without

8-oxo-guanine in the  sequence to 0.7 in the sample containing 8-oxo-guanine. Therefore, it can be assumed that the polymerase used in the preparation kit does not lead to termination at the 8-oxo-guanine modification, but causes an erroneous readout (T instead of G). In the detection of CPD lesions performed in the main text, no increase of certain sequences with increasing exposure dose was found. Thus, the sequence preparation actually resulted in a termination of the polymerase activity and not in an erroneous readout.

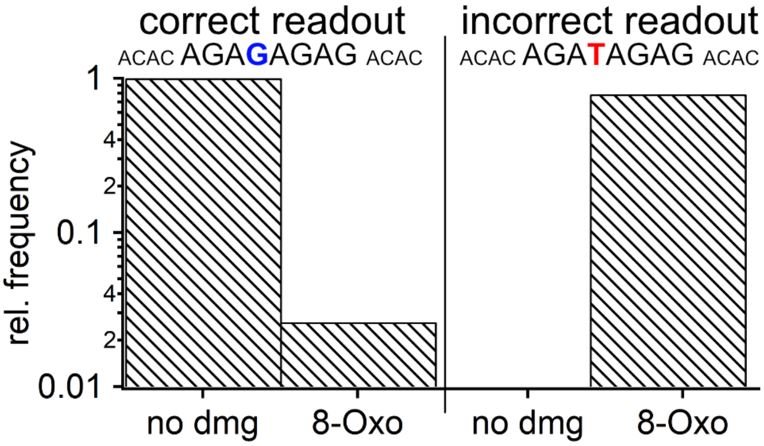

**Figure SI-f 14.** Change in the relative frequency of a target sequence ACAC AGAGAG ACAC upon insertion of an 8-oxo-guanine at position 8. While this sequence could be detected less frequently by a factor of 0.027 (left), the modification led to the appearance of an erroneous sequence in which a thymine was detected at position 8 instead of a guanine (right). The frequency of this altered sequence with the relative frequency of 0.7 corresponds approximately to the decrease in the frequency of the correct target sequence. Apparently, the polymerase used causes predominantly the replacement of 8-oxo guanine by a thymine instead of the a termination.

#### SI-17 Characteristic data on the dose dependences for tetramers and hexamers

**Table SI-t 8.** Decay parameters for tetramers. Total initial decay coefficients $\mu_{olig}$ , 50% dose values calculated from the initial decay constant $\mu_{olig}$ ($D_{50\%, mono}$) and from the fit-curves ($D_{50\%, fit}$).

$D_{50\%, fit}$ > $D_{50\%, mono}$ points to a pronounced bi-exponentiality of the dose dependence. The error $\Delta\mu_{olig}$ contains only the statistical part from the precision of the data points and the fitting procedure. It does not consider the systematic uncertainty (±20%) in the determination of the irradiation dose.

| Sequence number | Sequence | Total initial decay coefficient $\mu_{olig}$ [damage/photon] | Error $\Delta\mu_{olig}$ | $D_{50\%, mono}$ = ln2 /$\mu_{olig}$ | $D_{50\%, fit}$ |
|---|---|---|---|---|---|
| 0 | AAAA | 9.74e-03 | 4.98e-05 | 71 | 71 |
| 1 | AAAC | 7.13e-03 | 4.96e-05 | 97 | 97 |
| 2 | AAAG | 5.83e-03 | 4.85e-05 | 119 | 119 |
| 3 | AAAT | 7.83e-03 | 5.18e-05 | 88 | 88 |
| 4 | AACA | 5.26e-03 | 4.83e-05 | 132 | 132 |
| 5 | AACC | 5.60e-03 | 5.37e-05 | 124 | 124 |
| 6 | AACG | 4.60e-03 | 4.97e-05 | 151 | 151 |
| 7 | AACT | 1.59e-02 | 1.91e-03 | 43 | 89 |
| 8 | AAGA | 4.74e-03 | 4.81e-05 | 146 | 146 |
| 9 | AAGC | 3.99e-03 | 4.99e-05 | 174 | 174 |
| 10 | AAGG | 3.99e-03 | 5.07e-05 | 174 | 174 |
| 11 | AAGT | 4.16e-03 | 5.28e-05 | 167 | 167 |
| 12 | AATA | 5.67e-03 | 4.77e-05 | 122 | 122 |
| 13 | AATC | 5.75e-03 | 5.23e-05 | 121 | 121 |
| 14 | AATG | 4.10e-03 | 4.77e-05 | 169 | 169 |
| 15 | AATT | 2.88e-02 | 1.07e-03 | 24 | 35 |
| 16 | ACAA | 4.92e-03 | 4.90e-05 | 141 | 141 |
| 17 | ACAC | 3.70e-03 | 1.07e-03 | 187 | 194 |
| 18 | ACAG | 2.86e-03 | 4.97e-05 | 242 | 242 |
| 19 | ACAT | 1.02e-02 | 1.81e-03 | 68 | 163 |
| 20 | ACCA | 4.87e-03 | 1.23e-03 | 142 | 152 |
| 21 | ACCC | 7.37e-03 | 1.61e-03 | 94 | 123 |
| 22 | ACCG | 4.48e-03 | 1.21e-03 | 155 | 164 |
| 23 | ACCT | 2.37e-02 | 2.30e-03 | 29 | 107 |
| 24 | ACGA | 3.79e-03 | 4.96e-05 | 183 | 183 |
| 25 | ACGC | 3.05e-03 | 5.57e-05 | 227 | 227 |
| 26 | ACGG | 3.02e-03 | 5.49e-05 | 229 | 229 |
| 27 | ACGT | 3.81e-03 | 1.11e-03 | 182 | 190 |
| 28 | ACTA | 4.96e-03 | 5.28e-05 | 140 | 140 |
| 29 | ACTC | 2.66e-02 | 3.66e-03 | 26 | 116 |
| 30 | ACTG | 3.40e-03 | 5.26e-05 | 204 | 204 |
| 31 | ACTT | 3.56e-02 | 1.14e-03 | 19 | 28 |

| | | | | | |
|---|---|---|---|---|---|
| 32 | AGAA | 5.22e-03 | 4.87e-05 | 133 | 133 |
| 33 | AGAC | 3.06e-03 | 4.85e-05 | 226 | 226 |
| 34 | AGAG | 2.51e-03 | 4.90e-05 | 276 | 276 |
| 35 | AGAT | 3.73e-03 | 5.07e-05 | 186 | 186 |
| 36 | AGCA | 2.94e-03 | 4.94e-05 | 236 | 236 |
| 37 | AGCC | 4.13e-03 | 1.17e-03 | 168 | 182 |
| 38 | AGCG | 2.43e-03 | 5.11e-05 | 285 | 285 |
| 39 | AGCT | 1.12e-02 | 1.85e-03 | 62 | 162 |
| 40 | AGGA | 3.77e-03 | 5.01e-05 | 184 | 184 |
| 41 | AGGC | 2.71e-03 | 5.43e-05 | 256 | 256 |
| 42 | AGGG | 1.73e-03 | 5.51e-05 | 401 | 401 |
| 43 | AGGT | 3.35e-03 | 5.89e-05 | 207 | 207 |
| 44 | AGTA | 3.39e-03 | 4.90e-05 | 204 | 204 |
| 45 | AGTC | 3.15e-03 | 5.55e-05 | 220 | 220 |
| 46 | AGTG | 2.44e-04 | 5.52e-05 | 2844 | 2844 |
| 47 | AGTT | 1.54e-02 | 1.18e-03 | 45 | 81 |
| 48 | ATAA | 5.58e-03 | 4.82e-05 | 124 | 124 |
| 49 | ATAC | 3.62e-03 | 4.90e-05 | 192 | 192 |
| 50 | ATAG | 3.02e-03 | 4.82e-05 | 229 | 229 |
| 51 | ATAT | 1.36e-02 | 2.23e-03 | 51 | 143 |
| 52 | ATCA | 4.20e-03 | 5.14e-05 | 165 | 165 |
| 53 | ATCC | 1.86e-02 | 3.49e-03 | 37 | 147 |
| 54 | ATCG | 4.02e-03 | 5.37e-05 | 172 | 172 |
| 55 | ATCT | 2.42e-02 | 1.96e-03 | 29 | 92 |
| 56 | ATGA | 3.42e-03 | 4.80e-05 | 202 | 202 |
| 57 | ATGC | 2.69e-03 | 5.10e-05 | 258 | 258 |
| 58 | ATGG | 2.32e-03 | 5.02e-05 | 299 | 299 |
| 59 | ATGT | 3.16e-03 | 5.27e-05 | 220 | 220 |
| 60 | ATTA | 2.10e-02 | 1.07e-03 | 33 | 53 |
| 61 | ATTC | 3.08e-02 | 1.08e-03 | 23 | 32 |
| 62 | ATTG | 1.43e-02 | 1.15e-03 | 48 | 87 |
| 63 | ATTT | 5.12e-02 | 9.35e-04 | 14 | 16 |
| 64 | CAAA | 6.43e-03 | 5.14e-05 | 108 | 108 |
| 65 | CAAC | 4.31e-03 | 5.32e-05 | 161 | 161 |
| 66 | CAAG | 3.05e-03 | 5.12e-05 | 228 | 228 |
| 67 | CAAT | 1.55e-02 | 2.47e-03 | 45 | 141 |
| 68 | CACA | 3.35e-03 | 1.02e-03 | 207 | 226 |
| 69 | CACC | 6.13e-03 | 1.47e-03 | 113 | 193 |
| 70 | CACG | 2.88e-03 | 9.82e-04 | 241 | 262 |
| 71 | CACT | 2.20e-02 | 2.34e-03 | 32 | 154 |
| 72 | CAGA | 3.01e-03 | 5.09e-05 | 230 | 230 |
| 73 | CAGC | 3.02e-03 | 1.01e-03 | 229 | 262 |
| 74 | CAGG | 2.30e-03 | 5.68e-05 | 302 | 302 |
| 75 | CAGT | 3.40e-03 | 1.06e-03 | 204 | 225 |
| 76 | CATA | 3.46e-03 | 5.03e-05 | 200 | 200 |
| 77 | CATC | 2.38e-02 | 4.57e-03 | 29 | 182 |

| | | | | | |
|---|---|---|---|---|---|
| 78 | CATG | 2.17e-03 | 5.18e-05 | 319 | 319 |
| 79 | CATT | 3.07e-02 | 1.12e-03 | 23 | 35 |
| 80 | CCAA | 6.79e-03 | 1.46e-03 | 102 | 111 |
| 81 | CCAC | 6.16e-03 | 1.49e-03 | 112 | 196 |
| 82 | CCAG | 4.12e-03 | 1.19e-03 | 168 | 224 |
| 83 | CCAT | 2.25e-02 | 2.46e-03 | 31 | 163 |
| 84 | CCCA | 2.20e-02 | 3.48e-03 | 31 | 174 |
| 85 | CCCC | 3.13e-02 | 5.34e-03 | 22 | 173 |
| 86 | CCCG | 6.95e-03 | 1.71e-03 | 100 | 160 |
| 87 | CCCT | 3.73e-02 | 3.28e-03 | 19 | 100 |
| 88 | CCGA | 5.80e-03 | 1.39e-03 | 119 | 141 |
| 89 | CCGC | 6.12e-03 | 1.62e-03 | 113 | 177 |
| 90 | CCGG | 4.12e-03 | 1.27e-03 | 168 | 194 |
| 91 | CCGT | 1.42e-02 | 2.62e-03 | 49 | 184 |
| 92 | CCTA | 2.68e-02 | 3.75e-03 | 26 | 124 |
| 93 | CCTC | 4.15e-02 | 4.37e-03 | 17 | 106 |
| 94 | CCTG | 6.30e-03 | 1.53e-03 | 110 | 158 |
| 95 | CCTT | 4.56e-02 | 1.50e-03 | 15 | 21 |
| 96 | CGAA | 5.00e-03 | 5.28e-05 | 139 | 139 |
| 97 | CGAC | 4.05e-03 | 1.16e-03 | 171 | 191 |
| 98 | CGAG | 2.33e-03 | 5.36e-05 | 297 | 297 |
| 99 | CGAT | 6.06e-03 | 1.38e-03 | 114 | 147 |
| 100 | CGCA | 3.81e-03 | 1.13e-03 | 182 | 205 |
| 101 | CGCC | 5.53e-03 | 1.54e-03 | 125 | 190 |
| 102 | CGCG | 2.22e-03 | 6.41e-05 | 312 | 312 |
| 103 | CGCT | 2.25e-02 | 3.24e-03 | 31 | 153 |
| 104 | CGGA | 3.82e-03 | 5.78e-05 | 181 | 181 |
| 105 | CGGC | 3.75e-03 | 1.20e-03 | 185 | 209 |
| 106 | CGGG | 1.66e-03 | 6.30e-05 | 417 | 417 |
| 107 | CGGT | 4.65e-03 | 1.33e-03 | 149 | 174 |
| 108 | CGTA | 3.32e-03 | 5.45e-05 | 209 | 209 |
| 109 | CGTC | 4.33e-03 | 1.28e-03 | 160 | 187 |
| 110 | CGTG | 4.17e-04 | 6.51e-05 | 1663 | 1663 |
| 111 | CGTT | 1.72e-02 | 1.42e-03 | 40 | 82 |
| 112 | CTAA | 6.02e-03 | 5.64e-05 | 115 | 115 |
| 113 | CTAC | 2.52e-02 | 4.46e-03 | 28 | 152 |
| 114 | CTAG | 4.50e-03 | 1.21e-03 | 154 | 170 |
| 115 | CTAT | 2.46e-02 | 2.48e-03 | 28 | 118 |
| 116 | CTCA | 3.67e-02 | 4.55e-03 | 19 | 124 |
| 117 | CTCC | 3.75e-02 | 3.89e-03 | 18 | 108 |
| 118 | CTCG | 2.27e-02 | 3.80e-03 | 31 | 141 |
| 119 | CTCT | 4.29e-02 | 2.69e-03 | 16 | 64 |
| 120 | CTGA | 5.09e-03 | 1.27e-03 | 136 | 147 |
| 121 | CTGC | 5.32e-03 | 1.42e-03 | 130 | 189 |
| 122 | CTGG | 3.40e-03 | 1.11e-03 | 204 | 227 |
| 123 | CTGT | 9.52e-03 | 1.99e-03 | 73 | 208 |

| | | | | | |
|---|---|---|---|---|---|
| 124 | CTTA | 3.11e-02 | 1.16e-03 | 22 | 33 |
| 125 | CTTC | 4.17e-02 | 1.30e-03 | 17 | 21 |
| 126 | CTTG | 1.77e-02 | 1.35e-03 | 39 | 78 |
| 127 | CTTT | 6.19e-02 | 1.12e-03 | 11 | 13 |
| 128 | GAAA | 6.86e-03 | 5.04e-05 | 101 | 101 |
| 129 | GAAC | 4.10e-03 | 4.99e-05 | 169 | 169 |
| 130 | GAAG | 3.11e-03 | 4.91e-05 | 223 | 223 |
| 131 | GAAT | 5.29e-03 | 5.19e-05 | 131 | 131 |
| 132 | GACA | 2.79e-03 | 4.89e-05 | 248 | 248 |
| 133 | GACC | 3.65e-03 | 1.09e-03 | 190 | 206 |
| 134 | GACG | 2.33e-03 | 5.10e-05 | 297 | 297 |
| 135 | GACT | 1.25e-02 | 1.95e-03 | 55 | 164 |
| 136 | GAGA | 3.00e-03 | 4.94e-05 | 231 | 231 |
| 137 | GAGC | 2.01e-03 | 5.25e-05 | 345 | 345 |
| 138 | GAGG | 2.17e-03 | 5.49e-05 | 319 | 319 |
| 139 | GAGT | 2.49e-03 | 5.65e-05 | 279 | 279 |
| 140 | GATA | 3.34e-03 | 4.85e-05 | 207 | 207 |
| 141 | GATC | 3.99e-03 | 1.12e-03 | 174 | 184 |
| 142 | GATG | 2.03e-03 | 4.91e-05 | 342 | 342 |
| 143 | GATT | 2.37e-02 | 1.21e-03 | 29 | 59 |
| 144 | GCAA | 4.06e-03 | 5.21e-05 | 171 | 171 |
| 145 | GCAC | 2.98e-03 | 9.99e-04 | 233 | 303 |
| 146 | GCAG | 1.95e-03 | 5.40e-05 | 356 | 356 |
| 147 | GCAT | 1.10e-02 | 1.91e-03 | 63 | 232 |
| 148 | GCCA | 3.69e-03 | 1.12e-03 | 188 | 223 |
| 149 | GCCC | 5.59e-03 | 1.54e-03 | 124 | 200 |
| 150 | GCCG | 2.35e-03 | 6.44e-05 | 295 | 295 |
| 151 | GCCT | 2.51e-02 | 2.82e-03 | 28 | 148 |
| 152 | GCGA | 2.70e-03 | 5.35e-05 | 257 | 257 |
| 153 | GCGC | 1.83e-03 | 6.43e-05 | 378 | 378 |
| 154 | GCGG | 1.71e-03 | 6.19e-05 | 405 | 405 |
| 155 | GCGT | 2.79e-03 | 1.01e-03 | 248 | 259 |
| 156 | GCTA | 3.84e-03 | 5.71e-05 | 181 | 181 |
| 157 | GCTC | 2.64e-02 | 4.49e-03 | 26 | 166 |
| 158 | GCTG | 1.88e-03 | 5.93e-05 | 369 | 369 |
| 159 | GCTT | 3.01e-02 | 1.26e-03 | 23 | 36 |
| 160 | GGAA | 4.98e-03 | 5.18e-05 | 139 | 139 |
| 161 | GGAC | 2.36e-03 | 5.19e-05 | 294 | 294 |
| 162 | GGAG | 2.05e-03 | 5.46e-05 | 338 | 338 |
| 163 | GGAT | 3.30e-03 | 5.31e-05 | 210 | 210 |
| 164 | GGCA | 2.28e-03 | 5.39e-05 | 304 | 304 |
| 165 | GGCC | 3.11e-03 | 1.10e-03 | 223 | 289 |
| 166 | GGCG | 1.85e-03 | 6.10e-05 | 374 | 374 |
| 167 | GGCT | 1.45e-02 | 2.60e-03 | 48 | 202 |
| 168 | GGGA | 2.53e-03 | 5.49e-05 | 274 | 274 |
| 169 | GGGC | 1.04e-03 | 6.22e-05 | 670 | 670 |

| | | | | | |
|---|---|---|---|---|---|
| 170 | GGGG | 7.69e-04 | 4.66e-05 | 901 | 901 |
| 171 | GGGT | 1.71e-03 | 6.84e-05 | 406 | 406 |
| 172 | GGTA | 3.01e-03 | 5.41e-05 | 230 | 230 |
| 173 | GGTC | 2.96e-03 | 1.06e-03 | 234 | 245 |
| 174 | GGTG | 9.25e-04 | 7.53e-05 | 749 | 749 |
| 175 | GGTT | 1.42e-02 | 1.46e-03 | 49 | 106 |
| 176 | GTAA | 4.57e-03 | 5.04e-05 | 152 | 152 |
| 177 | GTAC | 2.25e-03 | 5.19e-05 | 309 | 309 |
| 178 | GTAG | 2.15e-03 | 5.22e-05 | 323 | 323 |
| 179 | GTAT | 4.31e-03 | 1.15e-03 | 161 | 184 |
| 180 | GTCA | 2.95e-03 | 5.62e-05 | 235 | 235 |
| 181 | GTCC | 5.27e-03 | 1.43e-03 | 132 | 178 |
| 182 | GTCG | 2.41e-03 | 6.27e-05 | 287 | 287 |
| 183 | GTCT | 2.23e-02 | 2.65e-03 | 31 | 137 |
| 184 | GTGA | 9.64e-04 | 5.61e-05 | 719 | 719 |
| 185 | GTGC | 1.00e-04 | 1.00e-04 | 6931 | |
| 186 | GTGG | 1.50e-04 | 7.51e-05 | 4621 | 4621 |
| 187 | GTGT | 2.31e-04 | 7.32e-05 | 2997 | 2997 |
| 188 | GTTA | 1.30e-02 | 1.20e-03 | 53 | 92 |
| 189 | GTTC | 2.00e-02 | 1.50e-03 | 35 | 76 |
| 190 | GTTG | 4.30e-03 | 1.23e-03 | 161 | 194 |
| 191 | GTTT | 3.40e-02 | 1.06e-03 | 20 | 26 |
| 192 | TAAA | 7.64e-03 | 5.25e-05 | 91 | 91 |
| 193 | TAAC | 7.35e-03 | 1.48e-03 | 94 | 102 |
| 194 | TAAG | 4.33e-03 | 5.19e-05 | 160 | 160 |
| 195 | TAAT | 1.63e-02 | 1.71e-03 | 42 | 108 |
| 196 | TACA | 5.39e-03 | 1.27e-03 | 129 | 149 |
| 197 | TACC | 1.39e-02 | 2.19e-03 | 50 | 167 |
| 198 | TACG | 4.48e-03 | 1.19e-03 | 155 | 181 |
| 199 | TACT | 2.16e-02 | 1.83e-03 | 32 | 109 |
| 200 | TAGA | 4.41e-03 | 1.15e-03 | 157 | 166 |
| 201 | TAGC | 4.54e-03 | 1.20e-03 | 153 | 198 |
| 202 | TAGG | 3.40e-03 | 1.04e-03 | 204 | 214 |
| 203 | TAGT | 7.58e-03 | 1.32e-03 | 91 | 200 |
| 204 | TATA | 1.05e-02 | 1.74e-03 | 66 | 147 |
| 205 | TATC | 1.69e-02 | 1.97e-03 | 41 | 130 |
| 206 | TATG | 4.34e-03 | 1.15e-03 | 160 | 189 |
| 207 | TATT | 3.28e-02 | 1.01e-03 | 21 | 30 |
| 208 | TCAA | 1.74e-02 | 1.88e-03 | 40 | 104 |
| 209 | TCAC | 2.15e-02 | 2.36e-03 | 32 | 167 |
| 210 | TCAG | 8.63e-03 | 1.47e-03 | 80 | 190 |
| 211 | TCAT | 2.61e-02 | 1.99e-03 | 27 | 111 |
| 212 | TCCA | 2.21e-02 | 2.21e-03 | 31 | 128 |
| 213 | TCCC | 3.07e-02 | 2.87e-03 | 23 | 110 |
| 214 | TCCG | 1.80e-02 | 2.29e-03 | 38 | 149 |
| 215 | TCCT | 3.76e-02 | 2.23e-03 | 18 | 58 |

| | | | | | |
|---|---|---|---|---|---|
| 216 | TCGA | 1.38e-02 | 1.80e-03 | 50 | 132 |
| 217 | TCGC | 1.84e-02 | 2.64e-03 | 38 | 157 |
| 218 | TCGG | 1.21e-02 | 1.99e-03 | 57 | 164 |
| 219 | TCGT | 1.36e-02 | 1.70e-03 | 51 | 134 |
| 220 | TCTA | 2.96e-02 | 2.25e-03 | 23 | 89 |
| 221 | TCTC | 4.37e-02 | 2.77e-03 | 16 | 65 |
| 222 | TCTG | 1.68e-02 | 2.03e-03 | 41 | 136 |
| 223 | TCTT | 4.85e-02 | 1.25e-03 | 14 | 18 |
| 224 | TGAA | 5.06e-03 | 5.30e-05 | 137 | 137 |
| 225 | TGAC | 4.74e-03 | 1.23e-03 | 146 | 191 |
| 226 | TGAG | 2.58e-03 | 5.39e-05 | 269 | 269 |
| 227 | TGAT | 1.07e-02 | 1.56e-03 | 65 | 180 |
| 228 | TGCA | 4.63e-03 | 1.23e-03 | 150 | 192 |
| 229 | TGCC | 1.02e-02 | 2.26e-03 | 68 | 245 |
| 230 | TGCG | 2.85e-03 | 1.02e-03 | 243 | 277 |
| 231 | TGCT | 1.60e-02 | 1.79e-03 | 43 | 149 |
| 232 | TGGA | 4.50e-03 | 1.19e-03 | 154 | 165 |
| 233 | TGGC | 4.15e-03 | 1.22e-03 | 167 | 253 |
| 234 | TGGG | 2.11e-03 | 8.64e-04 | 328 | 352 |
| 235 | TGGT | 6.99e-03 | 1.58e-03 | 99 | 204 |
| 236 | TGTA | 4.32e-03 | 1.16e-03 | 161 | 168 |
| 237 | TGTC | 9.41e-03 | 1.95e-03 | 74 | 205 |
| 238 | TGTG | 5.32e-04 | 6.14e-05 | 1302 | 1302 |
| 239 | TGTT | 1.78e-02 | 1.24e-03 | 39 | 71 |
| 240 | TTAA | 2.80e-02 | 1.09e-03 | 25 | 37 |
| 241 | TTAC | 2.88e-02 | 1.19e-03 | 24 | 41 |
| 242 | TTAG | 2.01e-02 | 1.21e-03 | 34 | 74 |
| 243 | TTAT | 3.27e-02 | 1.03e-03 | 21 | 30 |
| 244 | TTCA | 3.35e-02 | 1.05e-03 | 21 | 28 |
| 245 | TTCC | 4.15e-02 | 1.35e-03 | 17 | 22 |
| 246 | TTCG | 2.99e-02 | 1.21e-03 | 23 | 33 |
| 247 | TTCT | 5.10e-02 | 1.27e-03 | 14 | 17 |
| 248 | TTGA | 2.04e-02 | 1.15e-03 | 34 | 63 |
| 249 | TTGC | 2.13e-02 | 1.34e-03 | 33 | 73 |
| 250 | TTGG | 1.46e-02 | 1.22e-03 | 47 | 102 |
| 251 | TTGT | 1.89e-02 | 1.14e-03 | 37 | 65 |
| 252 | TTTA | 4.89e-02 | 9.89e-04 | 14 | 17 |
| 253 | TTTC | 6.11e-02 | 1.05e-03 | 11 | 13 |
| 254 | TTTG | 3.78e-02 | 9.80e-04 | 18 | 23 |
| 255 | TTTT | 7.71e-02 | 9.22e-04 | 9 | 10 |

**Table SI-t 9.** Hexamer sequences with largest and smallest survival probabilities, determined from the fit curves at large irradiation dose values D = 500 photon/base i.e. 3000 photon/oligomer.

Oligomers with small irradiation damage, large survival probability > 60%:

GGGTGG, GTGCGC, GGTGGC, GTGTGG, GTGTCC, AGGGGC, CACGTG, GTGCCA, GTGGCC, AGTGGC, GGTGGT, GTGTGC, TAGTGG, GTGCGA, AGGTGG, AGTGCA, GTGGTC, GGTGCA, CAGTGG, GTGTAC, CTGTGG, GTGTGT, TGTGTC, AAGTGG, GTGCAC, CATGTG, GAGTGG, GCAGTG, AGTGTC, TCAGTG, GTGGAC, CCGTGG, GGGGGC, CCGTGA, GTGCCC, CGTGCA, CGTGCC, AGTGGT, CGTGGC, ACGTGC, TGTGCA, GTGCGT, CGGTGG, GTGGCA, GTGTCT, GTGCGG, AGCGTG, ATAGTG, TGTGCG, GAGTGA, GTGTCG, CGGTGC, GACGTG, GTGGCG, TGTGGC, AGTGCC, CAGGGG, GCGTGG, ATGTGC, CAAGTG, GTGAGC, CGGTGT, TGTGCC, CCTGTG, CGTGTG, GCTGGC, GCCGTG, CTAGTG, TGTGTG, GCGGTG, GTGATC, CGTGTC, GAGTGC, TGGGGC, GGGGGG, GATGTG, CAGGTG, GCGGGG, CGAGTG, AGTGCG

Oligomers with large irradiation damage, small survival probability < 0.75 %:

AAACTA, GGAATT, CTCCCT, AATTTA, TAAACT, AAAACA, CATTTT, ACAAAA, TCCTCG, TGAAAA, CTCTTA, TTTTAT, TCTACT, ATTCTC, GCTTTT, TTAAAG, ACTTTT, CGAAAA, AATTAT, TTTAAA, GAAAAC, TAAATT, TTAATT, TACTCT, ATAAAA, AGCCTT, TAAAAT, TTTCTT, CCTTCT, TTGAAA, TCAAAA, AGAAAA, AAACTC, TCCCTT, AAAACC, TTATCC, TTTTAA, AAAATC, ACTCCT, TCTAAA, GGGCTT, AAAATA, CTTTTC, TTACTT, CTAAAA, TTAAAC, CTTAAA, CCTTTC, AAATTC, TCCTTC, GGAAAT, AATTCT, AAATCT, ATCTCT, GTTAAA, TTTTTT, TTCTTA, AAATTT, TCTTGA, AACTTT, AAACCT, TTCTTT, AACTCT, AAAAAG, ATTTTT, AATTTT, GGAAAA, ATTGTT, AAACTT, AAATTG, TTAAAA, CAAAAA, CTTCTC, AAAGTT, TCTCCG, GAAAAT, TCTTCT, CTTATT, AAAACT, AATTAA, AAAAAC, TAAAAA, GAAAAA, GAAATT, ATTAAA, AAATTA, AAAAAT, GTTCTC, AAAAAA, CTTTCT, AAAATT
